# The three-dimensional landscape of chromatin accessibility in Alzheimer’s disease

**DOI:** 10.1101/2021.01.11.426303

**Authors:** Jaroslav Bendl, Mads E. Hauberg, Kiran Girdhar, Eunju Im, James M. Vicari, Samir Rahman, Pengfei Dong, Ruth Misir, Steven P. Kleopoulos, Sarah M. Reach, Pasha Apontes, Biao Zeng, Wen Zhang, Georgios Voloudakis, Ralph A. Nixon, Vahram Haroutunian, Gabriel E. Hoffman, John F. Fullard, Panos Roussos

## Abstract

Much is still unknown about the neurobiology of Alzheimer’s disease (AD). To better understand AD, we generated 636 ATAC-seq libraries from cases and controls to construct detailed genomewide chromatin accessibility maps of neurons and non-neurons from two AD-affected brain regions, the entorhinal cortex and superior temporal gyrus. By analyzing a total of 19.6 billion read pairs, we expanded the known repertoire of regulatory sequences in the human brain. Multi-omic data integration associated global patterns of chromatin accessibility with gene expression and identified cell-specific enhancer-promoter interactions. Using inter-individual variation in chromatin accessibility, we define *cis*-regulatory domains capturing the 3D structure of the genome. Multifaceted analyses uncovered disease associated perturbations impacting chromatin accessibility, transcription factor regulatory networks and the 3D genome, and implicated transcriptional dysregulation in AD. Overall, we applied a systematic approach to understand the role of the 3D genome in AD and to illuminate novel disease biology that can advance diagnosis and therapy.

---

Alzheimer’s Disease (AD) is a chronic neurodegenerative disorder characterized, clinically, by cognitive decline and, neuropathologically, by accumulation of amyloid beta (Aβ) plaques and intracellular neurofibrillary tangles. Although a growing number of common and rare genetic risk variants have been identified (*1*), the neurobiological causes and substrates of AD in the vast majority of the population remain unknown. As such, additional approaches should be considered in an attempt to better understand the molecular mechanisms that mediate this debilitating disease. Abnormalities in the epigenomic regulation of brain function could result from primary genetic and non-genetic causal factors and epiphenomena, including changes secondary to disease progression. Thus, the epigenome could elucidate disease mechanisms, especially in late-onset AD, where there can be a gap of multiple decades before the initial changes in brain function become clinically apparent.

A number of recent studies have identified AD-associated epigenomic changes (*2–6*). These studies, however, have been limited to bulk tissue and have only examined one brain region at a time. This, naturally, impedes identification of brain region and cell-type specific disease signatures. Furthermore, although analyses of bulk tissue can identify apparent changes in gene expression and the epigenome, bulk tissue level changes can reflect alterations in cell composition rather than changes in the function of individual cells. This is particularly important for neurodegenerative diseases, including AD, where disease progression involves neuronal loss.

Due to the inherent difficulty in obtaining fresh specimens, most molecular studies of the human brain are restricted to frozen post-mortem samples. Working with frozen material is not without its challenges, including the loss of cytoplasm (and, with it, many cell-specific antigens) as a consequence of freeze-thawing. Nevertheless, an increasing number of human brain studies have employed cell-type specific nuclear markers to isolate nuclei of interest via Fluorescence-Activated Nuclear Sorting (FANS) (*7–9*). Using FANS to study individual cell-types increases the power to identify cell-specific, disease-associated changes and, importantly, mitigates the aforementioned changes in cell type composition.

In this study, we expanded the panel of genomic assays in the Mount Sinai Brain Bank AD (MSBB-AD) cohort (*10*), a collection of human post-mortem brain samples from individuals who have been deeply phenotyped, both clinically and histopathologically. In particular, we generated genome-wide maps of chromatin accessibility using ATAC-seq from neuronal and non-neuronal nuclei isolated by FANS from the entorhinal cortex (EC) and superior temporal gyrus (STG) of AD cases and controls. Both brain regions are affected in AD, with the EC showing earlier and more severe pathological changes and neuronal loss (*11*). We initially used these data to expand the repertoire of identified cell type specific regulatory regions and studied their relationship to gene expression. We then examined the shared and distinct molecular mechanisms associated with clinical dementia and neuropathological lesions. Subsequently, we identified regulatory genomic signatures associated with AD, including variability in discrete open chromatin regions (OCRs), transcription factor (TF) regulatory networks and *cis*-regulatory domains (CRDs). AD-associated signatures showed brain-region and cell-type specificity, implicating non-coding regulatory regions within AD genetic risk loci that participate in a variety of biological pathways as well as changes in transcription factor regulation.

## Results

### Population-scale datasets of brain region and cell type specific chromatin accessibility expand the repertoire of known human brain regulatory elements

We performed ATAC-seq profiling in neuronal (NeuN+) and non-neuronal (NeuN-) nuclei isolated by FANS from the STG and EC of AD cases (n=153) and controls (n=56) derived from the Mount Sinai NIH Brain and Tissue Repository (Fig. 1A). All samples were part of the MSBB-AD cohort, which has been extensively analyzed as part of Accelerating Medicines Partnership - Alzheimer’s Disease (AMP-AD) project and have additional functional omics data, including whole genome sequencing and multiregional RNA-seq profiling (*10*) (Data S1). The individuals were selected to represent the full spectrum of clinical and pathological severity (Data S2) based on the following phenotypes: **(1)** case-control status defined using the Consortium to Establish a Registry for Alzheimer’s Disease (CERAD) criteria (*12*); **(2)** Braak AD-staging score for progression of neurofibrillary neuropathology (Braak & Braak-score or BBScore) (*13, 14*); **(3)** mean density of neuritic plaques (plaque mean); and **(4)** assessment of dementia and cognitive status based on clinical dementia rating scale (CDR) (*15*). These phenotypes were moderately correlated (fig. S1), indicating shared and distinct disease processes.

**Fig. 1.**
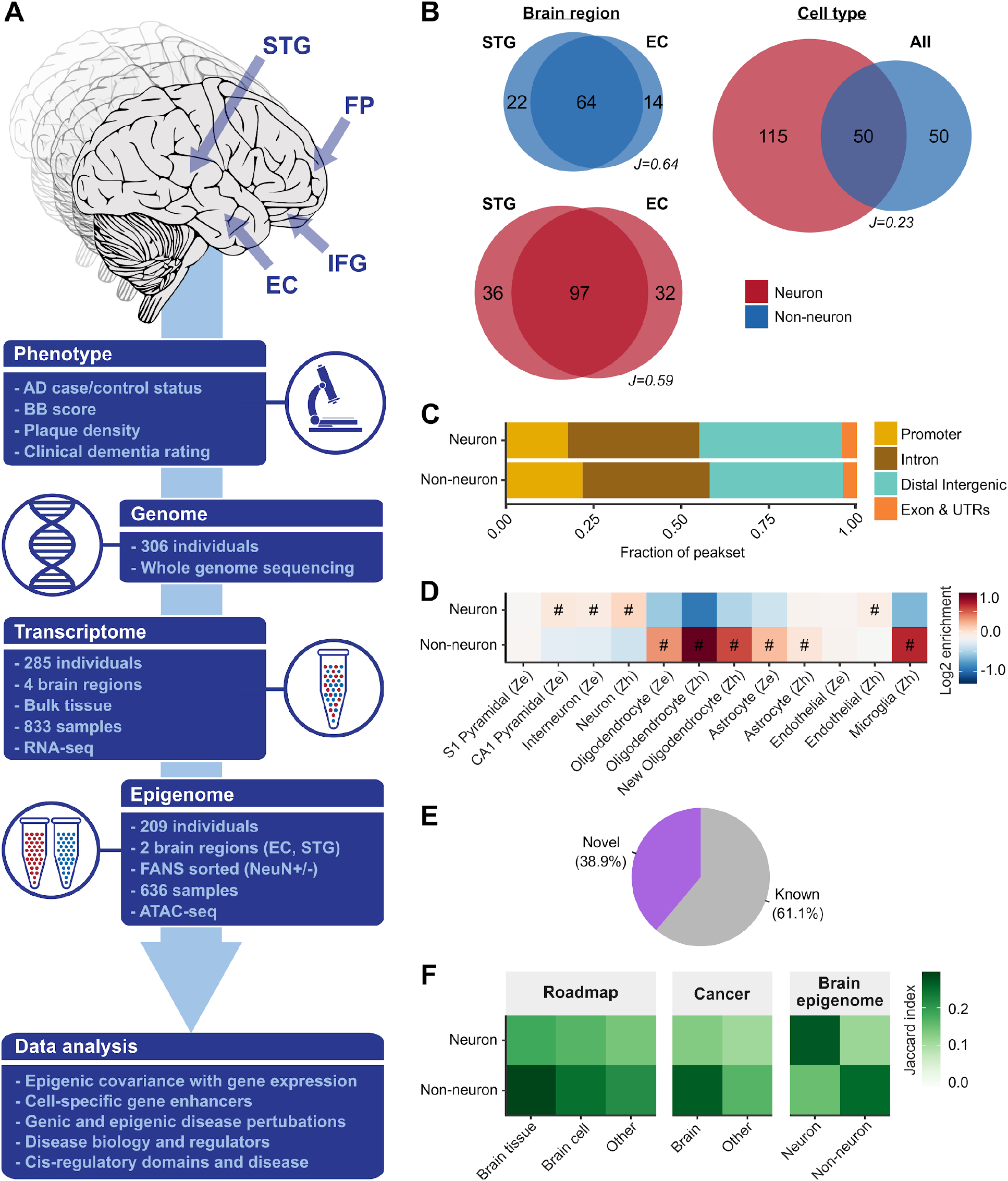
Large-scale chromatin accessibility analysis in the human brain. (**A**) Description of the study design. ATAC-seq was performed on neuronal (NeuN+) and non-neuronal (NeuN-) nuclei isolated from two different brain regions (STG and EC) in MSBB-AD samples. Existing RNA-seq profiling in the same cohort was performed on bulk tissue derived from four different brain regions (EC, IFG, FP and STG). All samples have genomic and AD-related phenotypic data. (**B**) Venn diagrams by cell type and brain region summarizing the overlap in megabases of OCRs. “J” indicates the Jaccard index between the respective OCRs. (**C**) Proportions of neuronal and non-neuronal OCRs stratified by genomic context. (**D**) Heatmap showing enrichment of neuronal and non-neuronal OCRs with cell type specific markers from “Ze”: Zeisel (*16*) and “Zh”: Zhang (*17*). “#”: Test wide significant at FDR<0.05 “·“: Nominally significant at *P*-value<0.05. (**E**) Novelty of OCRs compared to known OCRs from the union of three broad OCR resources (Roadmap Epigenomics Consortium, The Cancer Genome Atlas, and a brain epigenome atlas). (**F**) Overlap of neuronal and non-neuronal OCRs with sets of reference studies (as in panel E). For this, we considered only the known OCRs.

We processed 403 brain dissections from 2 brain regions to yield a total of 773 neuronal (NeuN+) and non-neuronal (NeuN-) FANS-sorted samples (fig. S2). Chromatin accessibility in each sample was then determined through ATAC-seq profiling. Extensive quality control of ATAC-seq libraries based on cell-type, sex, and genotype concordance, as well as sample quality metrics and sequencing depth, yielded a total of 636 samples constituting a total of 19.6 billion read pairs with an average of 30.8 million non-duplicated read pairs per library (fig. S3 and S4, Data S3). Given the large differences in chromatin accessibility profiles between the two cell types (Fig. 1B, fig. S5), neuronal and non-neuronal samples were considered separately for subsequent downstream analysis. A total of 315,630 neuronal and 205,120 non-neuronal open chromatin regions (OCRs) were identified, respectively. One reason for this high number of OCRs is the sequencing depth in the aggregated read files used for peak calling. The OCRs were proximal to genes (Fig. 1C) and, as expected, showed enrichment in known cell type markers (Fig. 1D).

Next, we compared our extensive catalog of OCRs to previous reference studies from the Roadmap Epigenomics Project (*18*), Cancer Atlas (*19*), and the Brain Open Chromatin Atlas (*20*). We identified 157,944 novel OCRs (126,903 neuronal and 68,054, partially overlapping, non-neuronal OCRs), thereby expanding the repertoire of known OCRs in human brain tissue/cells by 58%. In addition, 61.1% of the OCRs identified in this study overlapped with previously observed regulatory elements detected in the OCR reference repositories (Fig. 1E), highlighting the consistency of our results with published datasets. We found a higher (median Jaccard *J*=0.21) overlap between OCRs from the reference studies and non-neuronal OCRs compared to neuronal OCRs (median Jaccard *J*=0.14), indicating that our open chromatin atlas significantly expands our knowledge of the neuroepigenome (Fig. 1F, fig. S6). Overall, we have generated the largest resource of chromatin accessibility in the central nervous system to date and expanded the repertoire of identified cell type specific regulatory sequences in the human brain.

### Proximal and distal chromatin accessibility explains variation in gene expression

Chromatin structure is integral to transcriptional regulation, with chromatin accessibility regulating gene expression by facilitating, or inhibiting, binding of the transcriptional machinery. Given the availability of RNA-seq data from the same individuals and brain region homogenates from which we generated our cell-type specific ATAC-seq data, we sought to quantify the relative contribution of proximal (i.e. promoter) and distal (i.e. enhancer) chromatin accessibility to transcriptional variance. We applied variance decomposition models to each of 20,709 expressed RNA-seq genes using the covariance of OCRs at transcription start sites (TSSs) and distal regulatory elements as inputs (fig. S7). We analyzed neuronal and non-neuronal OCRs separately, but included OCRs and gene expression from two brain regions (STG and EC) and corrected for donor effects by adding inter-individual covariance in the models.

In this model, more than 70% of expression variance (76.1% for neuronal and 71.7% for nonneuronal) was explained by promoter and enhancer OCRs, confirming that gene expression is broadly associated with chromatin accessibility (Fig. 2, A to B). To assess the validity of these findings, we permuted our dataset and, as expected, found that little variance was attributed to the epigenome in this shuffled analysis (fig. S8 and S9). The proportion of expression attributed to enhancer OCRs was twice as large in neuronal samples (Fig. 2C, Data S4), confirming a larger impact of distal regulatory mechanisms in neuronal cell types (*20*). In contrast, a higher proportion of expression variance in non-neuronal samples was attributed to promoter OCRs. Gene set enrichment analysis of the genes with the highest proportion of variance explained by promoter OCRs showed enrichment with known cell type markers (fig. S10A).

**Fig. 2.**
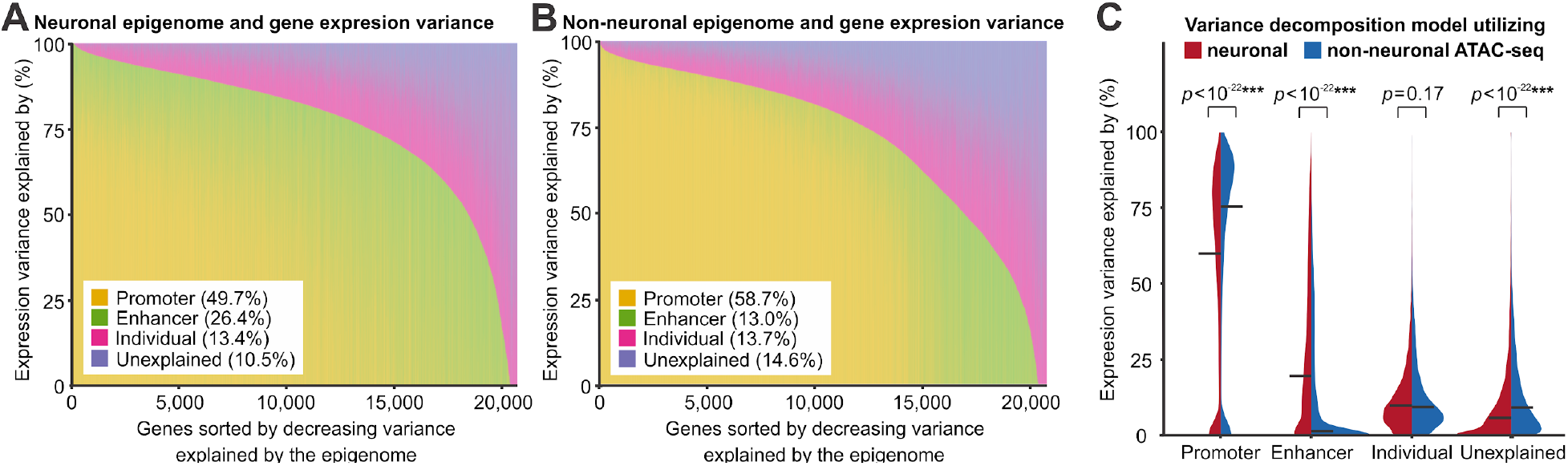
Variance component analysis of gene expression. The analysis was built upon (**A**) neuronal and (**B**) and nonneuronal open chromatin dataset. Genes, i.e., columns, are sorted by decreasing proportion of variance explained by epigenome (enhancer and promoter), with the mean variance explained by each component shown in parentheses. (**C**) Comparison of variance explained by each component across all RNA-seq genes. A Wilcoxon signed-rank test was used to test the differences between models using neuronal and non-neuronal ATAC-seq. Horizontal lines in the violin plot indicate median values.

The proportion of expression attributed to inter-individual covariance was similar for neuronal (mean 13.4%) and non-neuronal (mean 13.7%) samples. We hypothesized that the inter-individual covariance was, in part, driven by genetic regulation of gene expression (*21*). In order to test this hypothesis, we estimated, for each gene, the fraction of gene expression variation explained by the *cis*-genetic component via training transcriptomic imputation models with the EpiXcan method (*17*) in an independent RNA-seq + SNP-array dataset, comprised of human postmortem brains from the PsychENCODE Consortium (*22*); the cross validation R-squared (R^2^CV) model is known to be a good proxy for how much of the expression variance is driven by *cis*-genetic variation (*23*). Here we observed significant correlations between per-gene inter-individual covariance and R2CV in neuronal (Spearman *ρ*=0.22, *P*-value=1.3×10^-54^, fig. S10B) and non-neuronal (Spearman *ρ*=0.16, *P*-value=3.7×10^-30^, fig. S10C) OCRs. The remaining covariance was unexplained and was found to be larger in non-neuronal (mean 14.6%) compared to neuronal (mean 10.5%) samples. Overall, these results suggest that gene expression in human brain tissue is associated with global patterns of chromatin accessibility that differ between neuronal and non-neuronal cells.

### Neuronal and non-neuronal enhancer–promoter interactions regulate the majority of human brain expressed genes

Having shown that our data capture the transcriptional regulation associated with global patterns of proximal and distal chromatin accessibility, we next sought to determine enhancer-promoter (EP) interactions. E-P contacts have, with some success, been inferred from genomic distance and Hi-C derived chromatin loops (*24*). To improve upon this, however, the “activity-by-contact” (ABC) approach (*25*) model was employed, which quantifies the regulatory impact of enhancers quantitatively by assuming proportionality with both E-P contact frequency (inferred from Hi-C) and enhancer activity (inferred from chromatin accessibility and H3K27ac histone modification). To apply this approach using our ATAC-seq data, we generated cell type-specific ChIP-seq and Hi-C data.

Across the neuronal and non-neuronal datasets, we identified 37,056 and 38,233 E-P interactions, respectively. We determined that at least 63% of the expressed genes (13,135 of 20,709) were linked to one or more distal OCR (OCRABC) (Fig. 3A, fig. S11A, Data S5). While the majority of these enhancers were predicted to interact with a single gene, about one quarter seemingly regulated two or more genes (Fig. 3B, fig. S11B). Jointly, the enhancers that participate in E-P interactions cover 13.2-13.3Mb in each cell type (0.45-0.46% of the genome). On average, 43-47% of the E-P links were shared between neurons and non-neurons, whereas 83-90% of E-P links were shared across brain regions when comparing within the same cell type, with a high correlation of ABC score (fig. S11F). Among the predicted OCRABC enhancers, only 25% were linked to the nearest gene, clearly demonstrating the shortcomings of purely distance based regulatory annotation (Fig. 3F, fig. S11C). Still, the frequency of E-P links decreased sharply with distance and 86% were within 100kb (Fig. 3, C to E, fig. S11, D to E) (*26*).

**Fig. 3.**
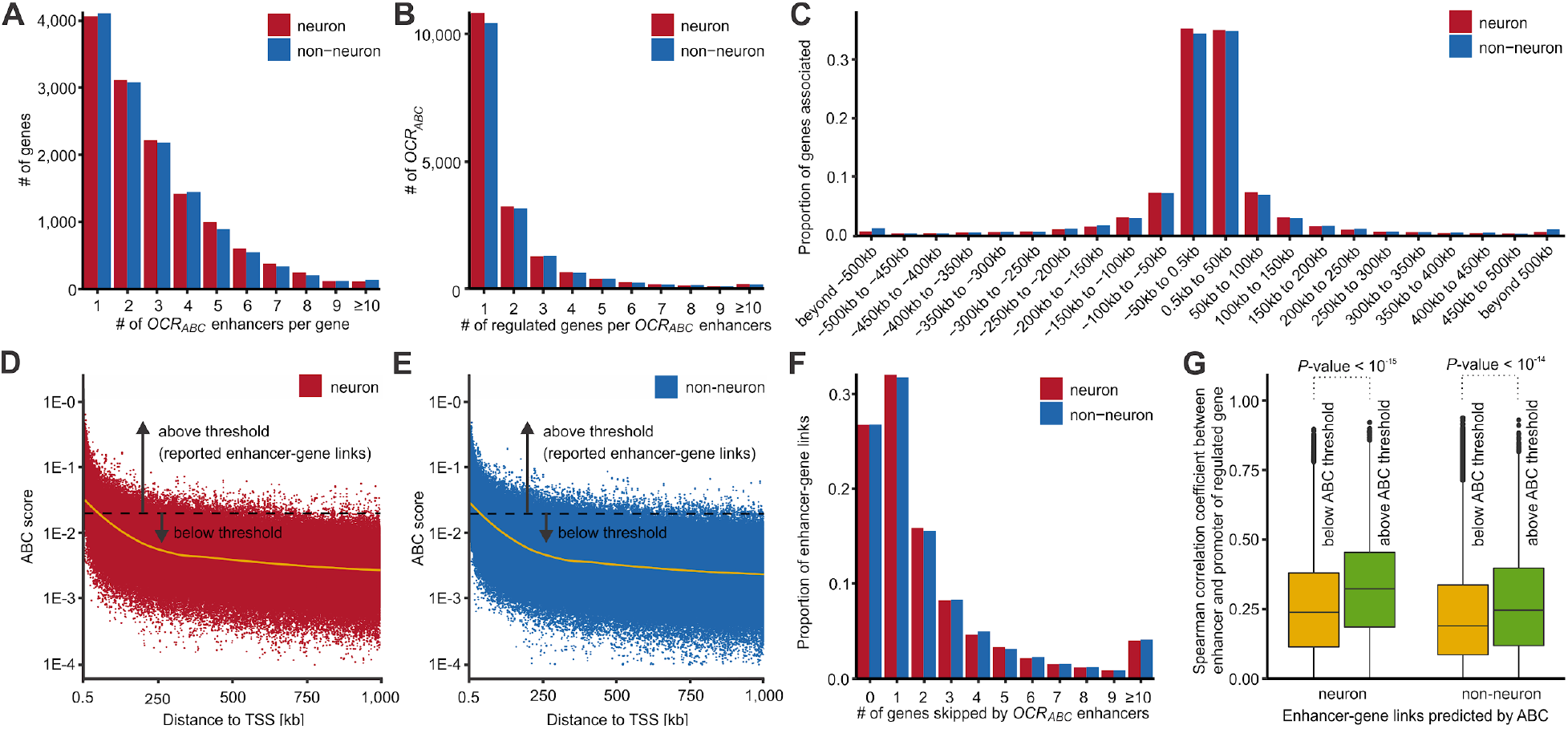
Linking distal regulatory OCRs (OCRABC) to genes using Activity-By-Contact (ABC) method. (**A**) Histogram of the number of OCRABC linked per gene. (**B**) Histogram of the number of genes linked per OCRABC. (**C**) Histogram of distance of OCRABC to the TSS of regulated genes. (**D-E**) Scatterplot of genomic distance vs ABC score of enhancer-gene links for (D) neurons and (E) non-neurons. The dashed black line denotes the minimum ABC score for an enhancer-gene link to be reported as a valid association. The solid yellow line is LOESS (local regression) fit. (**F**) Histogram of the number of genes “skipped” by an OCRABC to reach their linked genes. (**G**) Correlation between enhancer OCRs and promoter OCRs of linked genes from the ABC method, for links that did not meet the cut-off (low scores, OCRother) versus those that did (high scores, OCRABC). *P*-values were calculated by t-test from correlation values converted to Z-scores. The center line indicates the median, the box shows the interquartile range, whiskers indicate the highest/lowest values within 1.5x the interquartile range, and potential outliers from this are shown as dots.

To corroborate the results of our regulatory analyses, we compared distal OCRs predicted to participate in E-P interactions (OCR_ABC_) with a subset of distal OCRs not predicted to take part in such interactions but matched by distance to nearest TSS and OCR width (OCR_other_). We used orthogonal evidence for E-P interactions assigned based on fine mapping of GTEx eQTL analysis. OCR_ABC_, but not OCR_other_, were found to be enriched in fine-mapped eQTLs from GTEx (*27*) (OR 1.5-1.8, *P*-value < 10^-45^, fig. S12A, Data S6). When compared based on chromatin states from the Roadmap Epigenomics Project, OCR_ABC_ showed a relative depletion in repressed chromatin states as opposed to OCR_other_ (fig. S12, B to C). Finally, chromatin accessibility at predicted promoter regions was significantly more highly correlated with accessibility at OCR_ABC_ than accessibility at OCR_other_ (Fig. 3G, fig. S11G).

### AD associated epigenetic changes vary markedly by cell type and brain region

To explore AD-associated changes in chromatin accessibility, we performed differential analyses across multiple phenotypes (case/control status, BBScore, plaque mean and CDR) considering the brain regions (STG/EC) separately, or in combination, to increase statistical power (Fig. 4A, fig. S13, Data S7). EC neurons showed the highest number of associations across all AD phenotypes, with 19,336 OCRs (6.1%) associated with one or more AD phenotype(s) (Fig. 4B). For STG neurons, the corresponding number was 10,490 (3.3%), highlighting the regional specificity of AD-associated epigenetic changes. For non-neurons, the number of significant associations was markedly smaller, with STG and EC non-neurons jointly showing an association in 4,625 OCRs (2.3%) for one or more AD phenotype(s). Thus, the epigenomic changes in AD vary markedly by cell type and brain region. In terms of phenotypes, CDR and PlaqueMean identified the highest number of differential OCRs (Fig. 4B, Data S7 and S8) and, furthermore, yielded the highest π1 estimates, i.e. the highest estimates of the true proportion of differential OCRs (Data S9). These phenotypes, therefore, appear better powered to assay disease changes than the dichotomous casecontrol status.

**Fig. 4.**
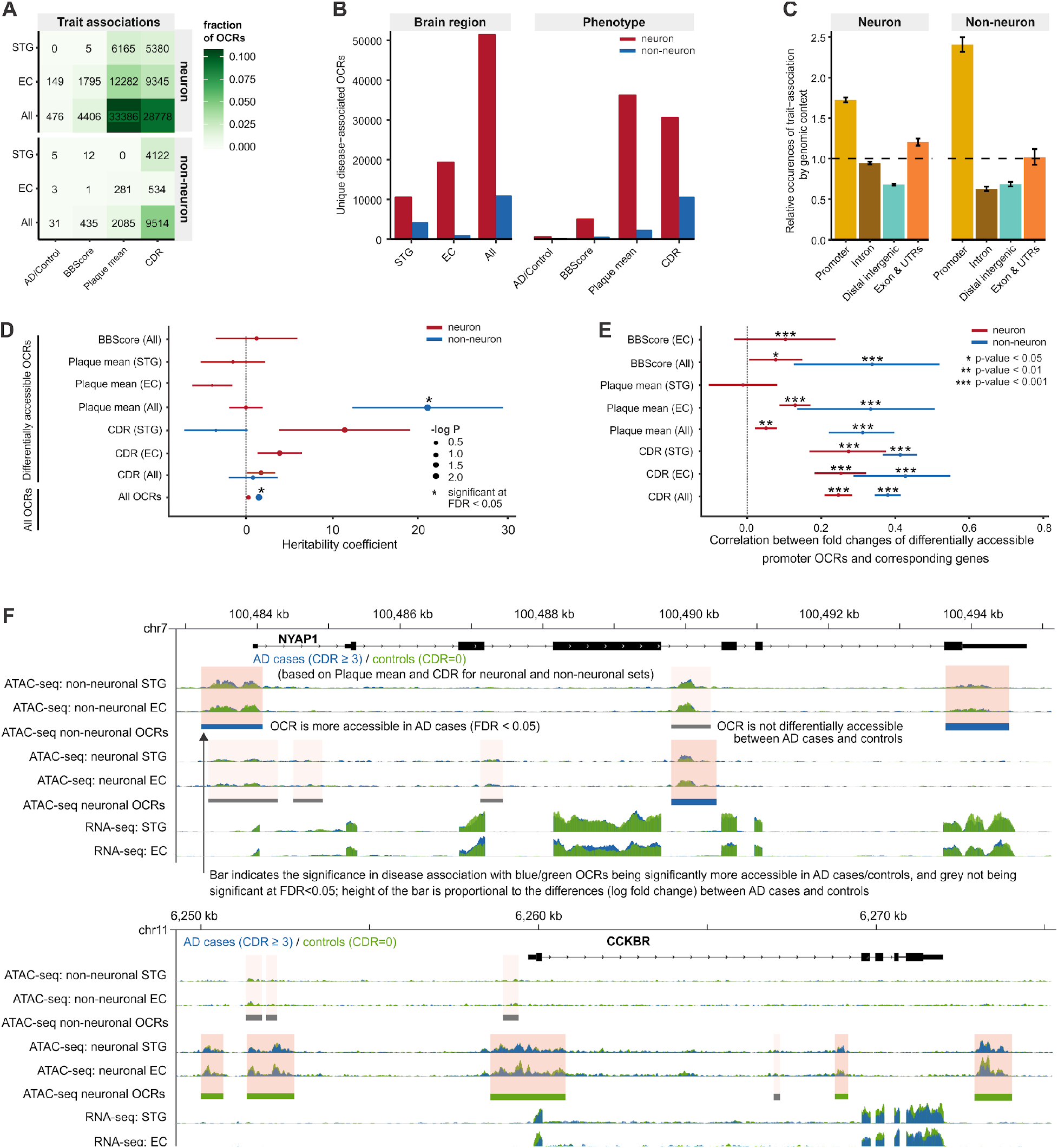
Disease-associated chromatin changes. (**A**) Disease-associated OCRs stratified by cell type, brain regions, and AD-related phenotypes. (**B**) Disease-associated OCRs stratified by brain regions or AD-related phenotypes (**C**) genetic context of disease-associated OCRs. (**D**) Enrichment of common genetic variants in AD with disease-associated OCRs when assayed by LD-score regression. Only sets of OCRs covering at least 0.05% of the genome were tested. Positive coefficients signify enrichment. “*”: FDR significant after correction across all tests. (**E**) Correlation between disease-associated chromatin changes at promoters (ATAC-seq) and disease-associated changes in gene expression (RNA-seq) at the corresponding genes. Pearson correlations were used and only comparisons with over 100 differential OCRs and genes were tested. (**F**) Examples of changes in chromatin accessibility and gene expression near the genes encoding *NYAP1* and *CCKBR*.

Epigenetically perturbed regulatory regions in AD were located near transcripts as well as in intergenic regions, suggesting that a combination of proximal and distal regulatory elements contribute to AD (Fig. 4C, Data S7). For both neurons and non-neurons, we found a higher fraction of AD-associated OCRs in the promoter area compared to all OCRs tested (Fig. 4C, fig. S14). However, a higher proportion of epigenetically perturbed regulatory regions in AD were located in proximity to the transcription start sites in non-neuronal compared to neuronal OCRs (Fisher’s exact one-sided *P*-value=1.07×10^-169^). We then examined the robustness of our differential chromatin accessibility analysis by checking the concordance (based on correlation of OCR foldchange) with three epigenetic studies of AD in human postmortem brains (*3*) and iPSC-derived neurons (*3, 28*) (fig. S15). We found significant concordance with an average Pearson coefficient of 0.31, while higher correlations were observed in neuronal compared to non-neuronal analysis (mean Pearson correlation of 0.38 vs 0.22).

While it is well-established that AD genetic risk variants are enriched in OCRs in microglia and astrocyte genes (*29*), the relationship with quantitative regulation of chromatin accessibility in AD is less clear. Besides considering the relationship between AD risk variants and OCRs genomewide, the detailed phenotype data of our samples allowed us to explore the relationship between genetic risk variation and AD-associated OCRs using LD-score partitioned heritability (*30*). Here, we found the set of all non-neuronal OCRs and the PlaqueMean-associated non-neuronal OCRs to be significantly enriched in AD genetic variants, even when accounting for the general genetic context of the OCRs (Fig. 4D). Interestingly, disease-associated non-neuronal OCRs showed a higher heritability coefficient than other OCRs highlighting the overlap between AD genetic risk variants and the subsequent epigenetic perturbations seen in the disease. Finally, we saw moderate enrichment when we overlapped OCRs with common variants of AD co-heritable traits, such as neuroticism, insomnia or bipolar disorder (*29*) (fig. S16, Data S10).

### AD associated epigenetic changes are concordant with gene expression alterations

To investigate changes in gene expression, we reprocessed 833 homogenate RNA-seq samples from four brain regions (including STG, EC, IFG - inferior frontal gyrus, and FP - frontal pole) in a highly overlapping set of individuals (fig. S17). The differential analysis was performed across multiple phenotypes as with the ATAC-seq data. A deconvolution parameter was added to account for differences in cell type composition in the bulk tissue, driven by the neuronal loss associated with disease progression (fig. S18 and S19). A higher proportion of genes was differentially expressed in EC and FP (24.4% and 21.5% of expressed genes, respectively), followed by STG and IFG (8.1% and 3.0% of expressed genes, respectively) (fig. S20, Data S11).

Comparing the epigenomic changes at transcription start sites with the changes in gene expression, revealed strong correlations (Fig. 4E, fig. S21). This concordance was higher for non-neuronal than neuronal samples (Pearson correlation of 0.31 and 0.14, respectively), suggesting that bulk tissue transcriptome studies in AD capture expression changes in non-neurons better than in neurons. Two illustrative examples demonstrating decreased chromatin accessibility and gene expression for AD cases through differentially regulated OCRs in promoters and distal intergenic regions are shown in Fig. 4F. *NYAP1* (Neuronal Tyrosine Phosphorylated Phosphoinositide-3-Kinase Adaptor 1), represents a late onset AD GWAS-linked candidate gene (*31*) that plays a role in the regulation of PI3K signaling pathway in neurons and controls cytoskeletal remodeling in outgrowing neurites (*32*). Loss of *CCKBR* (Cholecystokinin B Receptor) negatively affects spatial reference memory (*33*) and has repeatedly been identified among the top hits in differential gene expression analyses between AD cases and controls (*34–36*).

### Epigenome and transcriptome AD perturbations are enriched for biological processes and canonical pathways

To elucidate biological processes implicated in AD, we performed gene set enrichment analyses for our two ATAC-seq cell types, bulk RNA-seq, as well as the aforementioned GWAS study (29). We did this separately for each phenotype (fig. S22 and S23), and aggregated these by taking the most significant association across all phenotypes (Fig. 5, fig. S24).

**Fig. 5.**
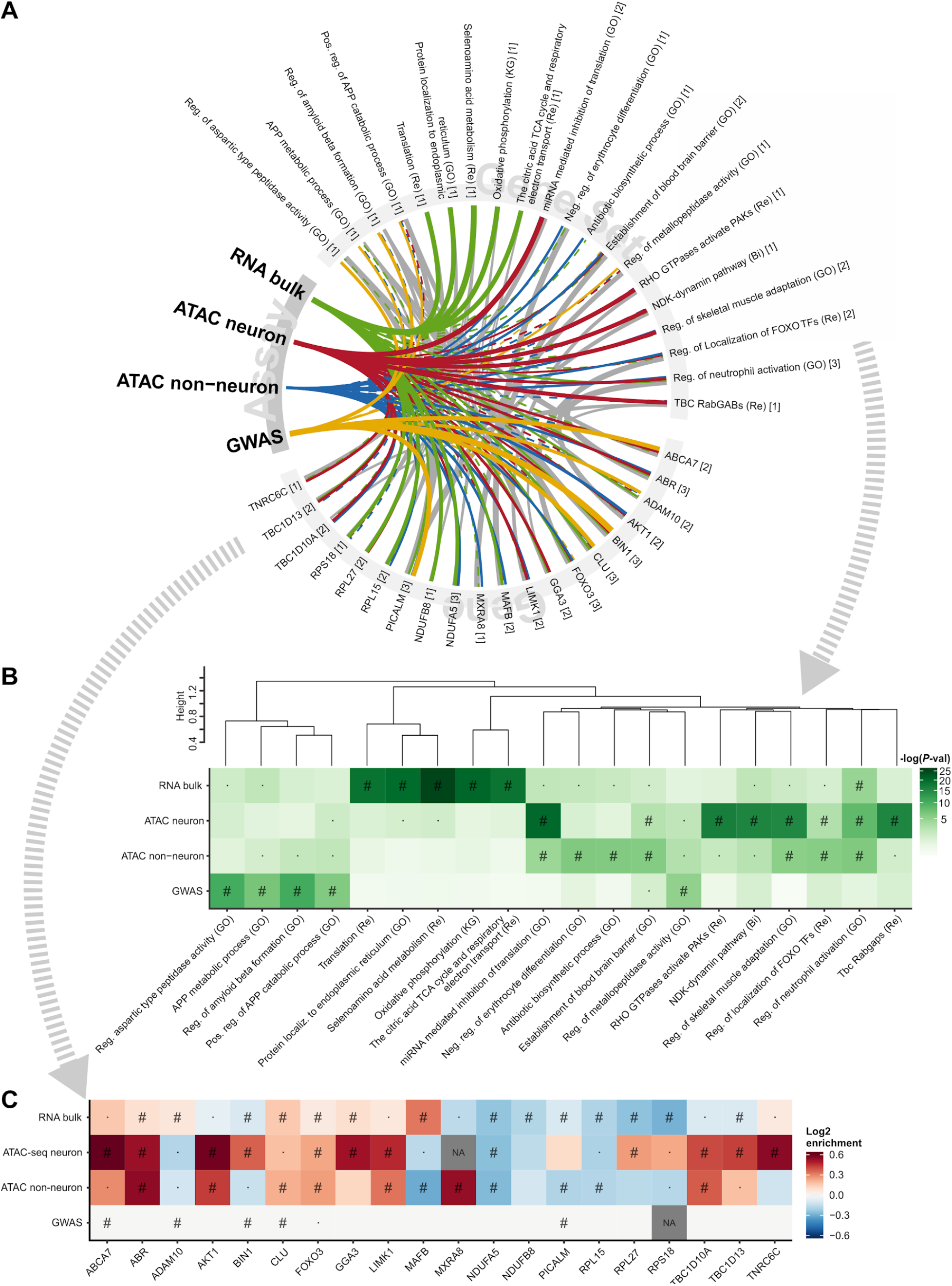
Gene set enrichment analysis using general gene sets. (**A**) Top five gene sets of the four overall assays and the top genes within these. Thickness indicates the strength of association. A full line indicates significance at FDR<5%, whereas a dashed line indicates nominal significance. Grey lines between genes and gene sets indicate gene set membership and the thickness is inversely proportional to the size of the gene set. The numbers in square brackets indicate the number of assays in which the gene or gene set was found significant at FDR<5%. The gene sets are clustered based on the constituent member genes. FDR throughout is calculated for all tests within one assay (e.g. all contrasts analyzed in ATAC Neuron times the number of gene sets tested). (**B**) heatmap of *P*-values for the associated gene sets. (**C**) Heatmap of fold changes in chromatin accessibility/gene expression and significance of change. The GWAS associations are genetic variants, which might increase or decrease or otherwise alter gene function. Thus no colors are applied to these. “#” Significance at FDR<5%. “·“: Nominal significance. “NA”: not available. “Bi”: Biocarta. “GO”: Gene Ontology. “KG”: Kegg. “Re”: Reactome. The ApoE- and MHC-loci were excluded from the GWAS analysis, as is customary. Not all genes have an OCR directly linked to it in neuron and non-neuron ATAC-seq.

The AD-associated changes in gene expression seen in RNA-seq implicated very general molecular pathways, such as “Oxidative Phosphorylation” and “Translation”. On the other hand, chromatin accessibility pointed to more specific molecular pathways. For example, using neuronal ATAC-seq we identified “the endocytotic role of NDK, phosphins and dynamin” and “RHO GTPase activation of PAKs”, among the top associations. Interestingly, PAKs (p21-activated kinases) are abundant in the brain where they have been associated with cell death and survival signaling (*37*). The non-neuronal ATAC-seq samples implicated “establishment of the blood-brain barrier”, which was also nominally significant with neuronal ATAC-seq, RNA-seq, and the AD GWAS. The blood-brain barrier has a purported role in the initiation and maintenance of chronic inflammation during AD (*38*). Other pathways associated with multiple assays included “regulation of neutrophil activation” and “regulation of localization of FOXO transcription factors”. Of note, the latter is involved in cell death and longevity (*39, 40*), and one member of the gene set, FOXO3 (*41*), showed at least a nominal association across all assays. Within these top gene sets, the top genes (e.g. *ABCA7*, *BIN1*, and *PICALM*) often showed an association across multiple assays (Fig. 5). We further highlight the top genes regardless of whether they were a member of one of the top gene sets or not (fig. S25, Data S12).

Finally, to examine changes in brain function, we performed a targeted gene set enrichment analysis using the synaptic gene ontology resource (*42*) (fig. S23 and S24). This implicated both pre- and post-synaptic dysfunction in post-mortem brain function. For the AD GWAS, synaptic dysfunction was also implicated, which, to the best of our knowledge, has not previously been reported. Of particular interest, “synaptic vesicle cycle” was significant in both the GWAS and RNA-seq, as well as in neuronal ATAC-seq, highlighting the shared enrichment of fundamental neuronal functions between the genetic drivers and subsequent chromatin accessibility and gene expression profiling in post-mortem brain.

### Cell type-specific transcription factor regulatory networks are perturbed in AD

Using footprinting analyses (*43*) we systematically examined transcription factor (TF) activity patterns underlying cell type differences and AD-related chromatin changes. We utilized 431 TF motifs, which, due to sharing of binding motifs, represented 798 TFs. TFs clustered into two cell type-specific groups with predominantly neuronal and non-neuronal regulatory patterns, complemented by a third group lacking cell-type specificity (Fig. 6, A to B, fig. S26A, Data S13). The TF cell specificity identified here was broadly concordant with existing literature (Data S14). For instance, Fos and Jun family members form an AP-1 heterodimer that is implicated in neuronal activity linked to learning and memory processes (*44, 45*). In contrast, Activating Transcription Factor 3 (ATF3) transcriptionally represses and regulates pro-inflammatory astrocytic and microglial activity (*46*). Genomic screen homeobox (GSX2), as a further example, acts as a gatekeeper of adult neurogenesis by mediating the recruitment in to the cell cycle of multipotent NSCs that can give rise to both neuronal and non-neuronal cells (*47, 48*). Corroborating our findings, the TF cell type specificities identified here were also highly concordant (Fig. 6A, Spearman *ρ* = 0.87, *P*-value < 10^-15^) with TF footprinting analyses performed in the BOCA study (Brain Open Chromatin Atlas, (*20*)). Next, we leveraged differences in footprinting signals from neuronal and non-neuronal samples (bound/unbound status per transcription factor binding site), to identify a set of 1,432 and 1,834 genes whose transcription were controlled exclusively by TFs in our neuronal and non-neuronal TF clusters, respectively (Data S15). Convincingly, these genes showed enrichment in known cell type markers of the appropriate cells (fig. S26B).

**Fig. 6.**
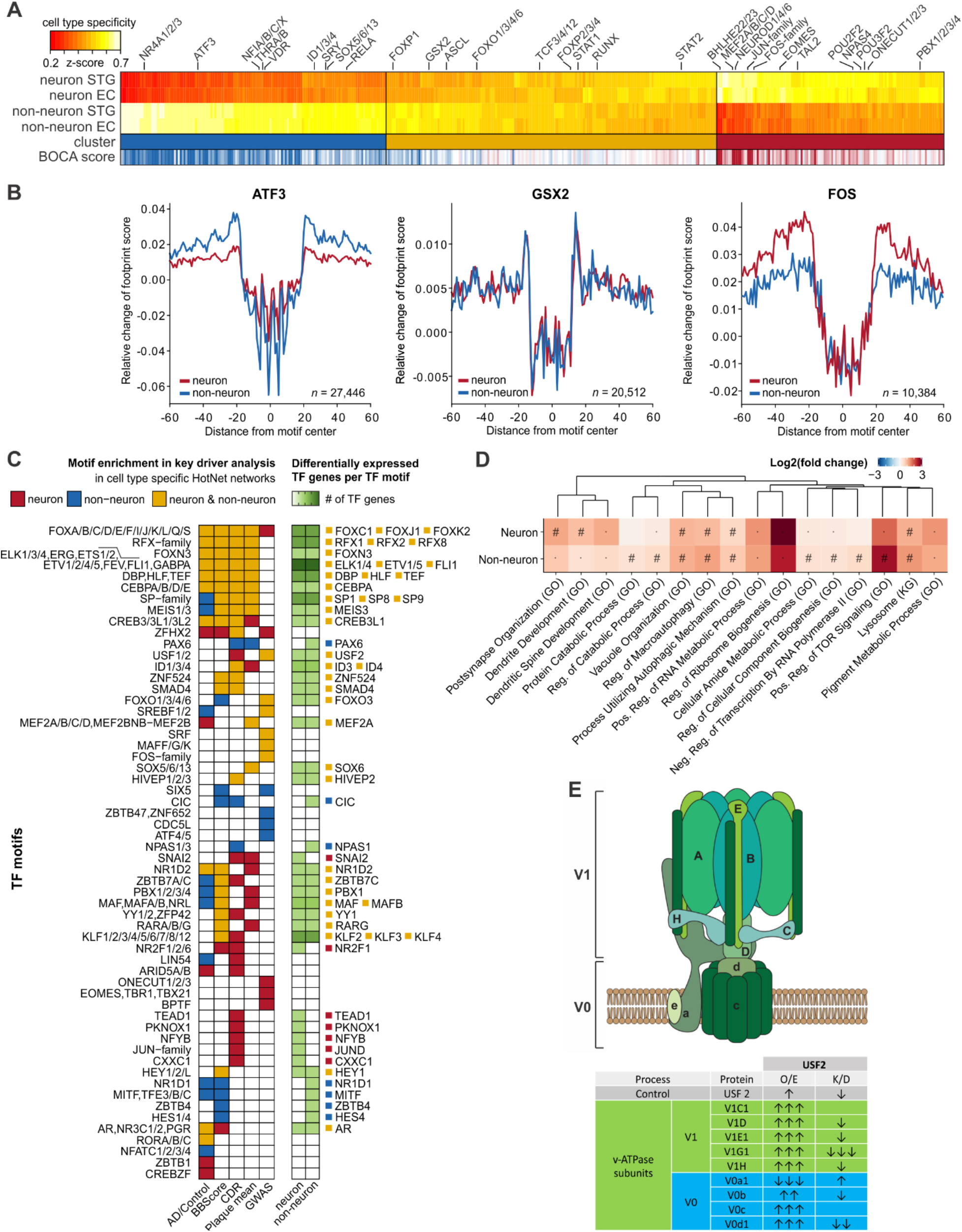
Mapping of transcription factors to cell types and AD-related phenotypes. (**A**) Hierarchical clustering of transcription factor motifs based on a cell type specificity score across neuronal and non-neuronal promoter OCRs. High specificity scores indicate high specificity to the given cell type. The concordance with Brain Open Chromatin Atlas is indicated as “BOCA score” ranging from blue (strong non-neuronal signal) to red (strong neuronal signal). Select motifs are further described in Data S14. (**B**) Aggregated footprint scores across all potential transcription factor binding sites for three motifs that represent the three major clusters of TFs according to cell type specificity. (**C**) Left: Overview of TF motifs whose neuronal and/or non-neuronal TFRNs show enrichment in AD genes. Right: subsequent prioritization of specific TF genes. The TF genes highlighted here meet the criteria of (1) significant correlation of their RNA-seq with TF regulatory targets and (2) significantly different TF gene expression between AD cases and controls. “TF”: transcription factor. (**D**) Heatmap showing top ten pathways for neuronal and non-neuronal genes regulated by USF2 transcription factor with general gene sets. “#”: FDR < 0.05 “·”: nominal *P*-value < 0.05. (**E**) Top: Schematic of the V0 and V1 subunits of the v-ATPase complex responsible for maintaining lysosomal pH. Bottom: Summary of western blot data in SH-SY5Y human neuroblastoma cells transfected with siUSF2 or stable overexpression of USF2. Arrow direction indicates protein level increase or decrease while the number of arrows represents statistical significance by Student’s two-tailed unpaired t-test, i.e. one arrow: *P*-value<0.05; two arrows: *P*-value<0.005; three arrows: *P*-value<0.0005.

Having broadly characterized TF activity in the two cell types, we then sought to examine TF dynamics in AD. For this, we constructed TF regulatory networks (TFRNs) for each cell type and brain region by analyzing actively bound TF motifs within proximal regulatory regions of genes representing the aforementioned 798 TFs. Due to high similarity of the network topology across brain regions (Jaccard *J* of 0.84 and 0.93 for neuronal and non-neuronal TFRNs, respectively), we subsequently merged TFRNs from the same cell type, resulting in a neuronal and a non-neuronal TFRN. Here, we identified a total of 26,976 and 40,348 unique, directed TF-to-TF interactions among the 431 analyzed TF motifs for neuronal and non-neuronal TFRNs, respectively, with a moderate overlap among the two networks (Jaccard *J*=0.56).

Using the neuronal and non-neuronal TFRN topology of directed TF-to-TF interactions and TF risk scores based on RNA-seq (across four AD-related phenotypes) and GWAS (*29*) data, we identified high-scoring subnetworks associated with AD (based on empirical *P*-value<0.05) (Fig. 6C). For each cell type, we subsequently combined the resulting subnetworks derived from the five TF risk scores into consensus subnetworks. Overall, we identified 46 and 45 TF motifs enriched in AD risk genes for the neuronal and non-neuronal TFRNs, respectively. To whittle down a list of candidate TF genes represented by the TF motifs prioritized by the TFRN analysis, we filtered out genes that were not differentially expressed between AD cases and controls (Fig. 6C, fig. S27). The resultant set of 56 TF genes shows a notable overlap with three studies that sought to identify AD regulatory hubs based on transcriptomics and epigenomics data (*49–51*). We found the TF gene USF2 to be highlighted in all three studies, as well as in our data, where AD perturbation on TFRN was supported by both GWAS and RNA-seq evidence.

Genes regulated by USF2 were enriched primarily in lysosomal dependent protein degradation pathways, which are processes known to be affected in AD (*52*) (Fig. 6D). While the literature on USF2 is very limited, there is evidence that USF2 plays a role in regulating lysosomal gene expression (*53*). To further support the link between USF2 and lysosomal dysfunction, we evaluated the effects of USF2 knockdown and over-expression on key components of the lysosomal pathway essential for maintaining lysosomal pH (*54*) and activity in a human neuronal cell line, SH-SY5Y. We found a reduction in the abundance of the members of the V0 (V0b, V0d1) and V1 (V1D, V1E1, V1G1, V1H) subunits of v-ATPase, which are responsible for maintaining intra-lysosomal pH (fig. S28, A to D), indicating impairments in lysosomal enzyme function. Convincingly, we observed changes in protein abundance in the opposite direction when USF2 was overexpressed (fig. S28, E to H). v-ATPase hydrolyzes ATP via its V1 domain and uses the energy released to transport protons across membranes via its V0 domain. This activity is critical for pH homeostasis and generation of a membrane potential that drives cellular metabolism (*55*). Accordingly, reduced levels of V1 sector v-ATPase subunits on lysosomes isolated from USF2 knockdown cells lowered the rate of ATP hydrolysis of v-ATPase by ~60% (fig. S28, I). The ATPase activity decline is expected to block proton translocation via the V0 sector (*56*) and impair acidification of lysosomes, as we observed (fig. S28, J to K). The increased level of V0a1 may be a compensatory, but incomplete, response to the loss of V1 subunits. Taken together, these findings suggest that the TF USF2 plays a role in maintaining pH dependent lysosomal function via v-ATPase activity.

### Three-dimensional chromatin interactions from population-level maps of chromatin accessibility

Studying correlation structures in chromatin accessibility at the population level has yielded high-resolution maps of *cis*-regulatory domains (CRDs) in lymphoblastoid cell lines (*57*). We employed this approach to explore the coordinated activity and chromatin organization of neuronal and non-neuronal OCRs in the adult human brain and its perturbations in AD (Fig. 7A, fig. S29 to S31). Initially, we explored the correlation of chromatin accessibility between pairs of OCRs and found that they decrease in frequency with genomic distance, which is in accordance with our ABC analyses of enhancer-promoter interactions (fig. S32A). Subsequently, we used adjacency constrained hierarchical clustering (*58, 59*), across both brain regions, to identify 13,671 STG neuronal, 13,334 EC neuronal and 8,861 STG non-neuronal, 8,688 EC non-neuronal CRDs, which included 37-39% of all OCRs (Fig. 7B, Data S17). Each CRD contained an average of 8 OCRs, but this varied substantially (29-30% with 5 or fewer OCRs; 44-45% with 9 or more OCRs). As expected, CRDs showed higher similarity across brain regions (Jaccard *J* of 0.33 for neurons and 0.34 for non-neurons) than across cell types (Jaccard *J* of 0.10 for STG and 0.10 for EC) (Fig. 7C). Next, we examined the composition of our CRDs and found 22% of OCRs within neuronal CRDs to be promoters, whereas 27% of OCRs within non-neuronal CRDs were promoters (fig. S32B). The remaining OCRs were enhancers. Further, a higher fraction of neuronal CRDs contained only enhancers, compared to non-neurons (fig. S32C). For example, ~40% of CRDs that include 8-10 OCRs are only enhancers in neurons, while non-neurons have a lower fraction (~30%) (fig. S32D). On average, within each neuronal CRD, a promoter coordinated with ~3.5 enhancers, whereas an enhancer coordinated with ~0.93 promoters. For non-neuronal CRDs, a promoter coordinated with ~3 enhancers, whereas an enhancer coordinated with ~1.1 promoters. Thus, the coordinated activity of OCRs within a CRD included a higher proportion of putative enhancers in neurons when compared to non-neuronal samples.

**Fig. 7.**
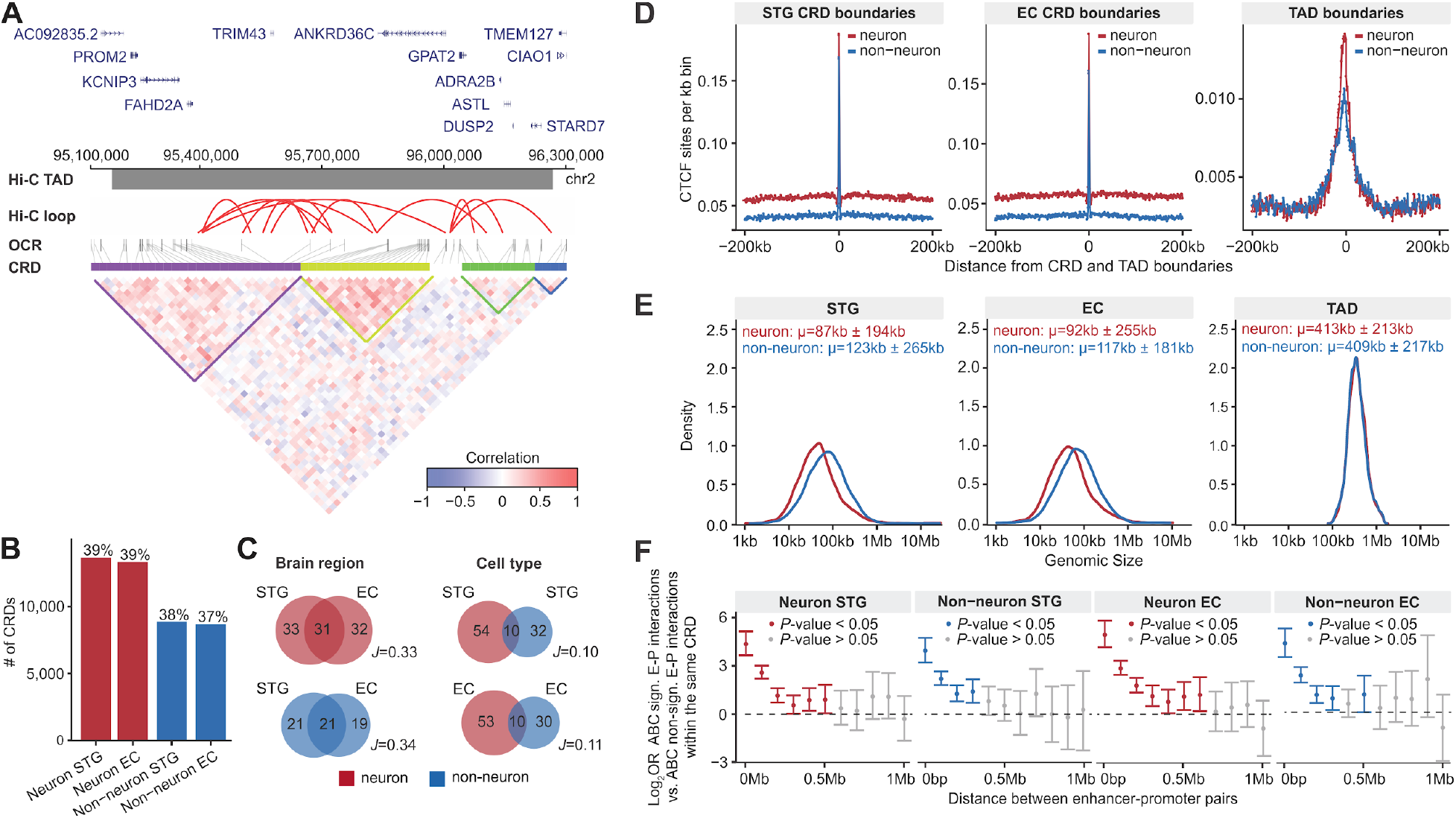
Definition of neuronal and non-neuronal *cis*-regulatory domains. (**A**) An example of *cis*-regulatory domains in STG neurons identified by estimating the interindividual correlation structure between nearby OCRs. From top to bottom: genes that are present in the locus; a single Hi-C TAD (dark gray horizontal bar) and Hi-C loops (red arcs); OCRs (gray vertical bars); CRDs, where each colorbar represents a different regulatory domain; correlation matrix of OCRs including colored triangles that highlight CRDs. (**B**) Number of CRDs stratified by cell type and brain region. The percentage is the fraction of OCRs that was within CRDs to the total number of OCRs. (**C**) Venn diagrams by cell type and brain region summarizing the overlap of CRDs. “J” indicates the Jaccard index between the respective CRDs. (**D**) CTCF density at and around CRD and TAD boundaries. (**E**) Size distribution of cell type and region specific CRDs and cell type specific TADs. (**F**) Associations of enhancer-promoter regions (ABC method) that are within the same CRDs. The odds ratios with their 95% confidence intervals are plotted as a function of the distance between enhancer and gene TSSs. *P*-values are estimated based on a two-sided Fisher’s exact test. CRD: *cis*-regulatory domain; TAD: topologically associating domain; kb: kilobase; and Mb: megabase.

Hi-C has been considered the gold standard in establishing the three-dimensional (3D) genomic structure (*60*). We compared CRDs to cell type-specific Hi-C-derived Topologically Associated Domains (TADs) to investigate the efficiency with which CRD captured the 3D genome. CCCTC-binding factor (CTCF) binding sites at TAD boundaries enhance contact insulation between regulatory elements of adjacent TADs (*61*). Both CRD and TAD boundaries were enriched in CTCF binding sites, with the former showing the highest density (Fig. 7D). Furthermore, there was a marked overlap between CRD and TAD boundaries (fig. S32, E and G). While TADs capture the spatial organization of the genome at the scale of hundreds of kb (mean of 413kb in neurons; 409kb in non-neurons), CRDs represent finer-scale regulatory clusters on the order of tens of kb (mean of 92kb in EC neurons; 87kb in STG neurons; 117kb in non-neurons; 123kb in non-neurons) (Fig. 7E). We leveraged Hi-C to divide the genome into A and B compartments, which are known to be associated with active and inactive chromatin, respectively (*62*), and confirmed that CRDs were enriched in type A compartments (fig. S32H). To explore the regulatory interactions within each CRD, we leveraged the outcome of the ABC analysis and found enrichment of E-P interactions that are within the same CRDs (Fig. 7F). Enrichment of CRDs with E-P interactions decreased sharply with distance, and interactions were not significantly enriched beyond ~500kb for neurons and ~250kb for non-neurons. Additionally, the correlation was higher for OCR interactions that have support for Hi-C loops and diminished with distance (fig. S32I).

In conclusion, interrogating inter-individual correlation between OCRs elucidates the coordinated activity of *cis*-regulatory elements and identified thousands of CRDs. These CRDs are highly concordant with, but substantially smaller than Hi-C-defined TADs, indicating that a CRD is a functional unit of the 3D genome, capturing intra-chromosomal interactions of active regulatory elements with higher spatial resolution.

### Differential analysis of CRDs defined perturbations of the three-dimensional genome in AD

We next took advantage of the CRD outcome to explore the association of AD with perturbations of the 3D genome. We performed differential analyses of CRDs across all AD-related phenotypes considering the brain regions (STG/EC) separately (fig. S29, Data S17). The AD epigenomic changes in CRDs varied markedly by cell type, brain region, and phenotype (Fig. 8A, Data S18), and showed similar patterns with the changes observed in the single OCR analysis (Fig. 4A). For instance, EC neurons showed the highest number of associations across all AD phenotypes, followed by STG neurons and then non-neuronal CRDs. The highest number of significant CRDs (n=2,603) was observed for the CDR phenotype in EC neurons and involved 26,365 OCRs (21.6%) that were also dysregulated (Fig. 8A). On average, across every phenotype, brain region, and cell type, dysregulation of the 3D genome in AD spanned 92.5 Mb or 3% of the genome (fig. S33, A to B). For the Plaque mean phenotype, perturbations of CRDs in EC neurons spanned 362.7 Mbp, or 12.1% of the genome, suggesting that restructuring of the 3D genome is widespread in these regions of the AD brain.

**Fig. 8.**
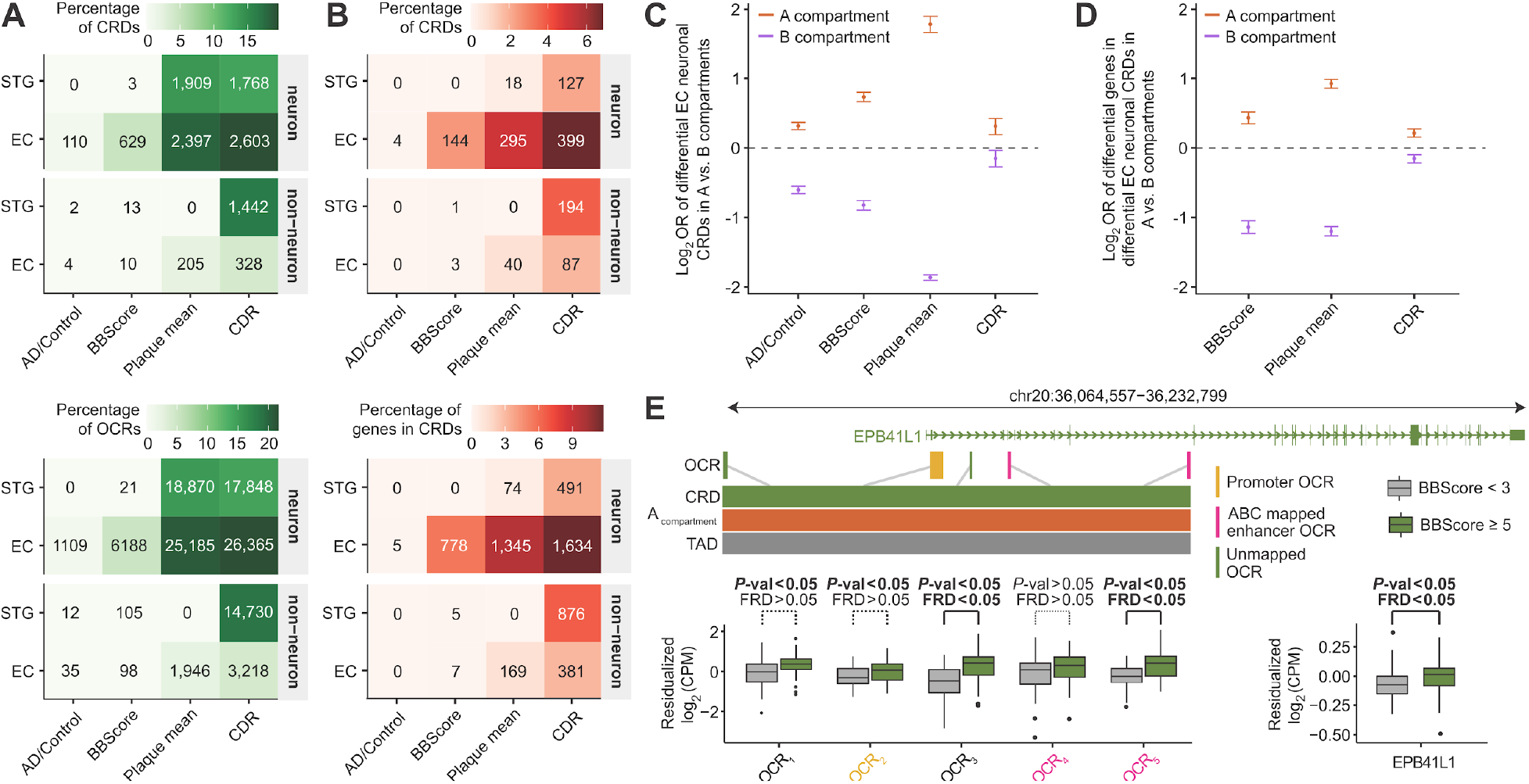
Disease-associated perturbations in CRDs. (**A**) Counts of CRDs and OCRs within CRDs that are associated with AD-related phenotypes stratified by cell type and brain regions. (**B**) Counts of CRDs and genes mapped to CRDs using the ABC model, that showed association with AD-related phenotypes in both epigenomic and transcriptomic levels using differential CRD test, stratified by brain region and cell type. (**C**) Odds ratios of disease associated EC neuronal CRDs to be in A compartments vs. B compartments, measured for each AD related phenotype. (**D**) Odds ratios of disease associated genes in EC neuronal CRDs to be in A compartments vs. B compartments, measured for each AD related phenotype. (**E**) Examples of changes in OCR expression and gene expression near the *EPB41L1* gene in BBScore associated CRD in EC neurons. Promoter and enhancer annotations of OCRs were obtained from ChIPSeeker and ABC results.

Having shown that the 3D genome is significantly altered in AD, we asked if AD-associated structural epigenomic perturbations cause transcriptional changes in nearby genes. For this, we mapped OCRs within CRDs to genes using the E-P interactions of our ABC analysis (Data S5). We next correlated changes in OCR accessibility with changes in gene expression and found our ABC-mapped CRD-derived OCRs to show a similar degree of correlation to genes when compared to the single OCR analysis (fig. S33, C to D). Using CRDs as the functional unit, we identified dysregulation of the transcriptome in AD (Fig. 8B, Data S18), which show similar cell type, brain region, and phenotype specificity as the epigenetically defined perturbations (Fig. 8A). Transcriptome-defined CRD perturbations were highly reproducible (range of π1 estimates from 0.39 to 0.88) in an independent gene expression dataset from the Religious Orders Study and Memory and Aging Project cohort (*63*) (fig. S34A, B). Gene set enrichment analysis for the transcriptome-defined CRD perturbations identified significant associations for multiple canonical pathways, including AD, cell adhesion molecules from the immunoglobulin family, chemokine receptors CXCR3 and CXCR4, SHC adaptor proteins, glutamate metabolic processes, and G protein coupled receptors (fig. S34, C to D).

A previous study reported a larger effect for tau-related changes in H3K9ac peaks located in Hi-C defined type A (active) compared to B (inactive) compartments (*3*). To elaborate on this finding, we characterized the spatial organization of differential CRDs in higher order chromatin structures i.e. type A vs. B compartments. Consistent with previous results (*3*), differentially upregulated CRDs in AD were enriched for type A compartments (Fig. 8C); in contrast, downregulated CRDs were enriched in B compartments. Spatial organization of differential CRDs was further corroborated by a negative correlation between the magnitude of AD-related phenotypic effects of CRDs that had no overlap with segments of nuclear lamina compared to those that had an overlap (lamina is associated with repressive chromatin) (*64*) (fig. S34E). An illustrative example of a differentially upregulated CRD in EC neurons associated with BBScore is shown in Fig. 8D. The spatial organization of this CRD was observed in type A (active) compartments in neurons and it involved proximal and distal OCRs regulating *EPB41L1*, which were all upregulated in more severe tau pathology (based on BBScores). The *EPB41L1* gene, encoding the Band 4.1-like protein 1 (also known as Neuronal protein 4.1) (*65*), links cell adhesion molecules to G protein coupled receptors at the neuronal plasma membrane (*66*) and has been associated with neurofibrillary tangles in AD brains (*67*).

## Discussion

Here we provide an extensive characterization of the cell type and regional chromatin regulatory landscape in human brain tissue derived from AD cases and controls. This resource increases our current understanding of the gene expression regulatory mechanisms in the human brain and deepens our understanding of AD etiopathogenesis. We initially utilized 19.6 billion read pairs from 636 individual ATAC-seq libraries to identify hundreds of thousands of cell type specific regulatory regions, expanding the annotation of the chromatin accessibility landscape in the brain from previous large-scale efforts (*18–20*).

Studying the epigenome at the population level allowed for broad inferences about gene regulation in the human brain. In fact, chromatin accessibility explained, globally, over 70% of the variance in gene expression. Building on this observation, we generated additional epigenome datasets and performed integrative analysis to define cell-specific enhancer-promoter interactions. This analysis identified putative links for more than half of the protein-coding genes in the genome, expanding previous efforts to catalogue E-P interactions in the human brain (*7*) and further increasing our understanding of the cell type-specific regulome in brain tissue. This valuable resource can be leveraged in future studies to better understand how genetic variation can affect regulatory regions, and to link these risk variants to specific genes.

AD is a highly complex disease, where both genetic and environmental factors, either directly or indirectly, shape the molecular perturbations that drive disease progression. In this study we explored the shared and distinct molecular mechanisms associated with clinical dementia and neuropathological lesions. We identified regulatory genomic signatures associated with AD, including variability in discrete open chromatin regions, TF regulatory networks and *cis*-regulatory domains.

Disease-associated changes to chromatin accessibility were extensive, involving thousands of regulatory sequences, many of which displayed specificity for a given cell-type and/or brain region. In addition to overlapping with AD common risk variation and gene expression perturbations revealed by bulk tissue analysis, these epigenetic changes also revealed additional cell-type specific AD associated molecular perturbations. Of note, common genetic variants related to the non-neuronal epigenome showed the most pronounced changes in AD, particularly so for non-neurons of the entorhinal cortex. Importantly, our FANS strategy accounts for the neuronal loss in AD and more precisely quantifies epigenetic dysregulation compared to bulk tissue studies, which, without applying deconvolution approaches, cannot distinguish molecular signatures arising through changes in cell type composition.

By applying footprinting analyses (*43*), we generated TFRNs and identified AD perturbations related to TF activity patterns that captured changes in gene expression and genetic variation in AD. This approach has the potential to elucidate biology as an aggregate of the multitudinous disease signals telling the story of gene dysregulation and dysfunction. As an example, we highlighted a putative role for USF2 in Alzheimer’s disease. This TF was predicted, in silico, to affect lysosomal genes, an observation that was supported by subsequent validation experiments using a human neuronal cell line.

Further, we used covariance of epigenetic changes to infer *cis*-regulatory domains, where OCRs work in concert to regulate nearby or more distant genes. These domains also recaptured chromosomal conformation information as evidenced by their overlap with Hi-C derived TADs. Using these domains to investigate changes in gene co-regulation structures turned out to be a powerful tool for interrogating epigenetic changes in AD. Of note, perturbed domains were enriched in active “A” compartments of the genome and depleted in inactive “B” compartments. On a more general level, we saw a disease associated decrease in both chromatin accessibility and gene expression for the lamina associated domains. This is in accordance with previous reports (*3*), but we do not yet understand the biological significance of this finding.

Collectively, this study augments our knowledge of the pathogenesis of AD, but also represents a valuable omics resource. With this cell specific map of chromatin accessibility, the MSBB-AD cohort contains information at the level of genetics, epigenomics and gene expression, which can be applied to downstream studies both genome wide and the single gene level, and, in time, should contribute to improved diagnosis and/or treatment of the disease.

## Materials and methods summary

The dataset of 636 ATAC-seq samples generated from postmortem human brains of 209 individuals represents a new addition to the panel of genomic assays in the Mount Sinai Brain Bank AD cohort. Brain tissue dissections spanning two brain regions were FANS-sorted to isolate DAPI positive neuronal (NeuN+) and non-neuronal (NeuN-) nuclei. ATAC-seq libraries were generated using an established protocol (*68*) and processed through our bioinformatics pipeline (fig. S2). We applied the variance component analysis (*69*) to quantify the proportion of gene expression variation that is attributable to promoter, enhancer and individual covariance. To predict enhancer-promoter interactions, we employed the “activity-by-contact” method that is based on the combination of frequency (derived from Hi-C) and enhancer activity (derived from ATAC-seq and H3K27ac ChIP-seq). To avoid the inflation of false discovery rate in the differential chromatin accessibility analysis, we applied *dream* (*70*), which properly accounts for correlation structures of repeated measures in the study designs utilizing cross-individual testing. For gene sets enrichment analysis with general (*71*) and synaptic (*42*) ontology gene sets, we employed *cameraPR* (*72*) that is taking into account test statistics from the differential analysis. Transcription factors involved in the regulation of gene expression were identified by footprinting analysis in *TOBIAS* (*43*) and modeled as a regulatory network. Then, *HotNet* (*73*) was used to find altered subnetworks containing TF motifs that are highly dysregulated between AD cases and controls based on transcriptomics and GWAS weights. Lastly, *decorate* (*59*) was used to identify co-regulated clusters of OCRs and run the differential analysis to find dysregulated AD-related clusters.

## Acknowledgments

We thank the patients and families who donated material for these studies. We thank the members of the Roussos laboratory for thoughtful advice and critique and the computational resources and staff expertise provided by the Scientific Computing at the Icahn School of Medicine at Mount Sinai.

## Author contributions

R.A.N., V.H. and P.R. conceived of and designed the project. V.H. provided human brain tissue. J.F.F. and P.R. designed experimental strategies for epigenome profiling in human postmortem tissue. R.M., S.K., S.M.R. and J.F.F. performed ATAC-seq data generation. S.R. performed Hi-C data generation. P.A. performed ChIP-seq data generation. E.I., J.V. and R.A.N. performed USF2 in vitro validation studies. J.B., M.E.H., K.G., G.E.H. and P.R. designed analytical strategies. J.B., M.E.H., K.G., and G.E.H conducted initial bioinformatics sample processing and quality control. J.B., M.E.H., K.G., and G.E.H. developed and performed all downstream omics data analyses and interpreted results. B.Z. performed eQTL fine-mapping analysis. W.Z. and G.V. performed transcriptome imputation analysis. J.B., M.E.H., K.G., G.E.H., J.F.F and P.R. wrote the manuscript with the help of all authors. G.E.H., J.F.F., and P.R. supervised overall data generation and analysis.

## Data and materials availability

Raw (FASTQ files) and processed data (BigWig files, peaks, and raw / normalized count matrices) are available via the AD Knowledge Portal (https://adknowledgeportal.org). The AD Knowledge Portal is a platform for accessing data, analyses, and tools generated by the Accelerating Medicines Partnership (AMP-AD) Target Discovery Program and other National Institute on Aging (NIA)-supported programs to enable open-science practices and accelerate translational learning. The data, analyses and tools are shared early in the research cycle without a publication embargo on secondary use. Data is available for general research use according to the following requirements for data access and data attribution (https://adknowledgeportal.org/DataAccess/Instructions). External validation sets: MSBB RNA-seq of postmortem brains (Synapse ID: <u>syn3157743</u>), H3K9ac ChIP-seq of postmortem brains (Synapse ID: <u>syn4896408</u>), ATAC-seq iPSC-derived neurons overexpressing MAPT gene (GEO <u>GSE97409</u>), fine-mapped eQTLs (https://alkesgroup.broadinstitute.org/LDSCORE/LDSC_QTL/, version “FE_META_TISSUE_GTEx_Brain_MaxCPP”), CTCF ChIP-seq on human neural cell (GEO <u>GSE127577</u>), The Cancer Genome Atlas (https://gdc.cancer.gov/about-data/publications/ATACseq-AWG), BOCA - brain epigenome atlas (https://icahn.mssm.edu/boca).

## Supplementary Materials

Materials and Methods

Figures S1-S34

Data S1-S18

References (74-119)

