## Supplement for "The three-dimensional landscape of chromatin accessibility in Alzheimer’s disease"

**This PDF file includes:**

Materials and Methods
Figs. S1 to S34
Captions for Data S1 to S18
References

**Other Supplementary Materials for this manuscript include the following:**

Data S1-S18 (.xlsx)

### Materials and Methods

Description of the post-mortem brain samples

Frozen brain tissue samples derived from STG (Superior Temporal Gyrus / Brodmann area 22) and EC (Entorhinal cortex / Brodmann area 36) were obtained from the Mount Sinai/JJ Peters VA Medical Center Brain Bank (MSBB–Mount Sinai NIH Neurobiobank), which holds over 2,000 human brains. All neuropsychological, diagnostic, and autopsy protocols were approved by the Mount Sinai and JJ Peters VA Medical Center Institutional Review Boards. AD cases and controls were selected to include donors with either no discernable neuropathology or cognitive complaints (controls) or those with only AD-associated neuropathology (cases; excluding donors with comorbid lesions such as significant cerebrovascular disease or Lewy bodies, etc.). Neuropathological assessments, cognitive, medical status, and neurological status was performed according to established procedures (*74*). Here neuropathological assessments was performed according to the Consortium to Establish a Registry for Alzheimer's Disease (CERAD) protocol (*12*) and included assessment by hematoxylin and eosin, modified Bielschowski, modified thioflavin S, and anti-β amyloid (4G8), anti-tau (AD2) and anti-ubiquitin (Dakoa Corp.). Further, a Braak AD-staging score for progression of neurofibrillary neuropathology was assigned to each sample (*13,14*). Additionally, the mean density of neuritic plaques in the middle frontal gyrus, orbital frontal cortex, superior temporal gyrus (STG), inferior parietal cortex and calcarine cortex (plaque mean) was calculated (*74*). For assessing cognitive function, the clinical dementia rating (CDR) was applied (*15*). Data S1 summarizes the demographic information of the present study population, including sex, age at the time of death, mean plaque density, CDR, and Braak & Braak score, stratified by cell type, brain region, and AD diagnosis. AD diagnosis was based on CERAD with cases consisting of "definite", "probable", and "possible" cases. The complete demographic information of the present MSBB AD study population is provided at https://dx.doi.org/10.7303/syn3159438 and https://dx.doi.org/10.7303/syn7392158.

FANS sorting of neuronal and non-neuronal nuclei

50 mg of frozen brain tissue was homogenized in chilled lysis buffer (0.32 M Sucrose, 5 mM $CaCl_2$, 3 mM Magnesium acetate, 0.1 mM, EDTA, 10 mM Tris-HCl, pH 8, 1 mM DTT, 0.1% Triton X-100) and filtered through a 40 µm cell strainer. Filtered lysate was underlaid with sucrose solution (1.8 M Sucrose, 3 mM $Mg(CH_3COO)_2$, 1 mM DTT, 10 mM Tris-HCl, pH 8) and subjected to ultracentrifugation at 107,000 xg for 1 hour at 4 ˚C. Pellets were resuspended in 500 µl DPBS containing 0.1% BSA. anti-NeuN antibody (1:1000, Alexa488 conjugated, Millipore, Cat# MAB377X) was added and samples incubated, in the dark, for 1 hr at 4 ˚C. Prior to FANS sorting, DAPI (Thermoscientific) was added to a final concentration of 1 µg/ml. DAPI positive neuronal (NeuN+) and non-neuronal (NeuN-) nuclei were isolated using a FACSAria flow cytometer with FACSDiva Version 8.0.1 software (BD Biosciences).

Generation of ATAC-seq libraries and sequencing

ATAC-seq libraries were generated using an established protocol (*68*). Briefly, 55,000 to 75,000 sorted nuclei were pelleted at 500 xg for 10 min at 4 ˚C. Pellets were resuspended in transposase reaction mix (22.5 µL Nuclease Free $H_2O$, 25 µL 2x TD Buffer; Illumina, Cat # FC-121-1030) and 2.5 µL Tn5 Transposase (Illumina, Cat # FC-121-1030) on ice and the reactions incubated at 37 ˚C for 30 min. Following incubation, samples were purified using the MinElute Reaction Cleanup kit (Qiagen Cat# 28204, and libraries generated using the Nextera index kit; Illumina Cat #FC-121-1011). Following amplification, libraries were resolved on 2% agarose gels and fragments ranging in size from 100-1000 bp were excised and purified (Qiagen Minelute Gel Extraction Kit – Qiagen, Cat# 28604). Next, libraries were quantified by quantitative PCR (KAPA Biosystems, Cat# KK4873) and library fragment sizes estimated using Tapestation D5000 ScreenTapes (Agilent technologie, Cat# 5067-5588). ATAC-seq libraries were subsequently sequenced on Hi-Seq2500 (Illumina) machines yielding 50 bp paired-end reads.

Generation of Hi-C libraries and sequencing

Hi-C data was generated from frozen postmortem human brain tissue using the *in situ* Hi-C protocol (*61*) with the following modifications. Frozen prefrontal cortex tissue was thawed at room temperature (RT) and dounce homogenized in HBSS (Hank's balanced salt solution). Homogenized tissue was fixed with 0.5% formaldehyde for 10 min and then quenched with 0.125 M glycine for 5 min at RT. Cross-linked tissue was then placed on ice for a further 15 min to quench crosslinking completely. Tissue was pelleted at 800 xg for 10 min at 4 ˚C and then resuspended in lysis buffer (0.32 M Sucrose, 5 mM $CaCl_2$, 3 mM $Mg(CH_3COO)_2$, 0.1 mM EDTA, 10 mM Tris-HCl, pH 8, 1 mM DTT, 0.1 % Triton X-100, 1 Roche cOmplete mini EDTA-free protease inhibitor tablet, Roche Cat# 4693159001) to isolate cross-linked nuclei. Neuronal nuclei were FANS sorted using the Alexa488 conjugated anti-NeuN antibody (1:1000) (Millipore Cat# MAB377X) on the BD FACS Aria II sorter. FANS sorted neuronal and non-neuronal nuclei were pelleted at 2500 xg for 5 min at 4 ˚C, frozen on dry ice for 20 min, and then stored at -80 ˚C.

Approximately 1 M crosslinked neuronal and non-neuronal nuclei were thawed on ice, washed with ice cold 1x CutSmart buffer (New England Biolabs (NEB), Cat# B7204S) and split into 4 aliquots to generate technical replicate libraries per sample, with 250k nuclei per library. Nuclei were pelleted at 2500 xg for 5 min at 4 ˚C, and resuspended in 342 µl 1x CutSmart buffer and then conditioned with 0.1% SDS at 65 ˚C for 10 min. Nuclei were immediately placed on ice and the SDS quenched with 1% Triton X-100. Chromatin was digested with 100 U of the 4 base pair cutter MboI (NEB Cat# R0147L) overnight at 37 ˚C with shaking at 400 rpm. MboI was heat inactivated at 65 ˚C for 20 min, and then nuclei were cooled down on ice. MboI cut sites were end labeled with biotin by adding 52 µl of biotin fill-in reaction mix (15 µl of 1 mM biotin-14-dATP (Jena Bioscience, Cat# NU-835-BIO14-L), 1.5 µl each of 10 mM dCTP, dGTP, dTTP (Sigma-Aldrich, Cat# DNTP10-1KT), 10 µl of 5 U/µl Klenow DNA Pol I (NEB, Cat# M0210L), 22.5 µl of 1x CutSmart buffer) and shaking at 37 ˚C for 1.5 h at 400 rpm. Blunt ended sites were proximity ligated by adding 948 µl ligation reaction mix (150 µl of 10x T4 DNA ligase buffer (NEB, Cat# B0202S), 125 µl of 10% Triton X-100, 15 µl of 10 mg/ml BSA, 10 µl of 400 U/µl T4 DNA ligase (NEB, Cat# M0202L), 648 µl $ddH_20$) and rotating tubes, end-over-end, at RT for 4 h. Nuclei were reverse crosslinked with 100 µl proteinase K (10 mg/ml) overnight at 65 ˚C.

Proximity ligated DNA was purified through phenol:chloroform extraction and sodium acetate/ethanol precipitation. Purified DNA was sheared using a Covaris S220 sonicator to generate a peak size of 400 bp with the following settings (peak incident power: 140 W, duty cycle: 10%, cycles per burst: 200, time: 55 sec). Biotin labeled ligation junctions were purified with Dynabeads MyOne Streptavidin C1 beads (ThermoFisher, Cat# 65001) by incubating for 1 hr at RT. Illumina compatible libraries were prepared from the sonicated and streptavidin bead immobilized DNA using the NEBNext Ultra II Library prep kit (NEB, Cat# E7645L), following manufacturer's instructions, amplifying libraries for 6-10 PCR cycles. Libraries were purified by 2-sided size selection (300-800 bp) using Ampure XP beads (Beckman Coulter, Cat# A63881).

All libraries were analyzed on a TapeStation using Agilent D5000 ScreenTapes, and quantified using the KAPA Library Quantification Kit prior to sequencing. Uniquely barcoded Hi-C libraries were pooled and deep sequenced on the Illumina NovaSeq S4 platform obtaining 100 bp paired end reads, to generate approximately 250 million reads per library.

<u>Generation of ChIP-seq libraries and sequencing</u>

Ultra-low input Native Chromatin Immunoprecipitation sequencing (ULI-NChIP-seq) libraries were prepared as follows. After FANS-sorting of neuronal (NeuN+) and non-neuronal nuclei (NeuN-), we performed ULI-NChIP assays, adapted from Brind'Amour, et al. 2015, which specifically does not require chromatin crosslinking, thereby increasing library complexity and reducing PCR artifacts. Briefly, nuclei were centrifuged at 500xg for 10 minutes at 4°C and re-suspended by gentle pipetting in residual smaller volumes of PBS/Sheath buffer. After DAPI counting of nuclei, 100K-300K were distributed into Eppendorf tubes and 0.1% Triton-X-100/0.1% Na-Deoxycholate added. The chromatin was re-suspended and placed at room temperature for 5 minutes, followed by fragmentation with micrococcal nuclease (MNase, NEB, #M0247S) for 5 min at 37°C on a ThermoMixer at 800 rpm to digest the chromatin to predominantly mononucleosomes. The MNase reaction was stopped by addition of 10% of the reaction volume of 100 mM EDTA (pipetted ~20x), followed by the addition of 1% Triton / 1% deoxycholate, (pipetted 5x) and placed on ice for at least 15 minutes. Samples were vortexed (medium setting) for ~ 30 seconds and complete NChIP buffer (20 mM Tris-HCl, pH 8.0, 2 mM EDTA, 15 mM NaCl, 0.1% Triton X-100, 1 EDTA-free protease inhibitor cocktail and 1 mM phenylmethanesulfonyl fluoride) was added to dilute the chromatin to <25% of the immunoprecipitation reaction volume, and rotated at 4°C for 1 hour. After the incubation, chromatin was vortexed (medium setting) for ~30 seconds and 5% input controls removed for DNA extraction.

To avoid non-specific binding, chromatin was next precleared by adding 10 µl/ reaction of a pre-washed 1:1 ratio of protein A to protein G Dynabeads (ThermoFisher, #10001D and #10003D), rotating the chromatin-protein A/ protein G magnetic bead mixture for 3 hours at 4°C. Antibody-bead complexes were prepared as follows: ChIP-grade histone H3K27ac antibody (Active Motif, #39133, pAB) was added to prewashed protein A/protein G Dynabeads resuspended in NChIP buffer, and the antibody-bead complexes formed by rotating the antibody and beads for 2 hours at 4°C. After the incubations, precleared chromatin and antibody-bead complexes were placed on a magnetic rack and the precleared chromatin transferred to new Eppendorf tubes, while the antibody-bead complexes were resuspended in sufficient volume of NChIP buffer to add 10 µl per MNase reaction. The chromatin was immunoprecipitated with the H3K27ac-bead complexes at 4°C overnight while rotating.

After the overnight incubation, the immunoprecipitation reactions were placed on a magnetic rack to remove unbound chromatin, washed twice with 200 µl of ChIP low salt wash buffer (20 mM Tris-HCl, pH 8.0, 0.1% SDS, 1% Triton X-100, 0.1% deoxycholate, 2 mM EDTA and 150 mM NaCl), twice with 200 µl ChIP high salt wash buffer (20 mM Tris-HCl (pH 8.0), 0.1% SDS, 1% Triton X-100, 0.1% deoxycholate, 2 mM EDTA and 500 mM NaCl), followed by elution in freshly prepared 30 µl of ChIP elution (100 mM sodium bicarbonate and 1% SDS) for 1.5 hours at 68°C on a ThermoMixer at 1000 rpm, followed by RNAse A digestion for 15 minutes at 37°C at 800 rpm. The immunoprecipitated DNA, along with the input controls, was purified using Phenol:Chloroform:Isoamyl Alcohol (25:24:1, v/v) (ThermoFisher Scientific, #15593-031), transferred to pre-spun phase lock tubes (Qiagen Maxtract #129046) to obtain the aqueous layer. An overnight ethanol precipitation was performed by adding 10 µl of 3M sodium acetate/100 µl aqueous layer of ChIP DNA, 1 µl of LPA (linear polyacrylamide, Sigma #56575) and 1 µl Glycoblue (Invitrogen, #AM9515).

NEBNext Ultra DNA Library Prep Kit (New England Biolabs, #E7370L) was used to construct NChIP libraries according to the manufacturer's directions, followed by Pippin Size selection using 2% Agarose Gel cassettes (SAGE Science) and cleanup with 1.8 volumes of SPRIselect beads (Beckman Coulter, #B23318). All libraries were analyzed on an Agilent High Sensitivity D1000 TapeStation, and quantified using the KAPA Library Quantification Kit prior to sequencing. The uniquely barcoded libraries were pooled and sequenced on a NovaSeq platform (Illumina). Approximately 40 million paired-end reads were generated per sample and subsequently aligned on hg38.

Processing of ATAC-seq data

An overview of the pipeline for the data processing of ATAC-seq samples is provided in fig. S2 and detailed in the following.

Preparing a masked reference genome and read mapping

FASTQ files delivered by the New York Genome Center were matched to the respective samples based on their pooling IDs and barcodes. It was checked that there was a one to one match and file integrity was confirmed using MD5 checksums. The raw reads files were trimmed using Trimmomatic (v0.36) (*75*) with the following settings:

```
ILLUMINACLIP:<NEXTERA_PAIR_END_ADAPTERS_FILE>:2:30:10:8:TRUE    LEADING:3
TRAILING:3 SLIDINGWINDOW:4:15 MINLEN:36.
```

To avoid biases in mapping on genomic regions with common genetic variants, the trimmed reads were mapped by STAR aligner on a modified version of hg38. Here we masked (a) pseudoautosomal regions PAR1 and PAR2 on chromosome Y and (b) biallelic SNPs that were present in WGS files of the participating individuals. The following settings of the STAR aligner (v2.5.0 (*76*)) were used:

```
--alignIntronMax 1
--outFilterMismatchNmax 100
--alignEndsType Local
--outFilterScoreMinOverLread 0.66
--outFilterMatchNminOverLread 0.66
```

For each sample, this yielded a BAM file of mapped paired-end reads sorted by genomic coordinates. From these files, reads that mapped to multiple loci were excluded using samtools (*77*), duplicated reads were excluded with PICARD (v2.2.4; http://broadinstitute.github.io/picard), and, lastly, reads mapping to the mitochondrial genome were discarded.

Genotype calling

For ATAC-seq samples, genotypes were called using GATK (v3.5.0) (*78*). In brief, these steps were performed: (1) indel-realignment; (2) base score recalibration; and (3) joint genotype calling across all samples. Variants with a phred-scaled confidence threshold < 10 were here discarded. All clustered variants, variants in ENCODE blacklisted regions of the genome, and variants not in dbSNP v151 (79) were not considered. Read depth was not used for filtering. Finally, only variants with minor allele frequencies (MAF) ≥ 25% were used. Pairwise genotype concordance was carried out amongst all ATAC-seq samples, RNA-seq samples, and whole genome sequencing (WGS) samples using the kinship coefficient from KING v1.9 (80), and considering only the variants called in the ATAC-seq data. The WGS and RNA-seq genotypes were called as previously described (10). For each assay combination (e.g. ATAC-seq vs. RNA-seq or WGS) proper cut-offs for the kinship values were selected based on the samples that were expected to be from the same brain versus those that were not. Using these cut-offs, genotype matches were categorized into three different categories: 1) Okay: matches that should be there and were there; 2) Suspicious: matches that should be there but were not there; and 3) Spurious: matches that should not be there but were there. The suspicious and spurious matches were all unambiguously resolved iteratively by a majority vote. For instance, if a sample had suspicious matches to the other samples from that person, but had spurious matches to all the samples from another brain, that sample was flagged and the person ID was corrected. When all such unambiguous sample swaps in a step were identified, the matching classification was redone, thus resolving suspicious/spurious matches and allowing the identification of further unambiguous mismatches, which could then be resolved, and so on, until all mismatches were resolved. The resulting fixed person IDs were then used to double test that the genotypes matched as expected between samples from the same brain and samples from different brains.

Sex determination of samples

Three different metrics were used to evaluate the sex of the samples: 1) the rate of heterozygotic genotyping calls on chromosome X outside the pseudoautosomal regions. For this, variants with a MAF < 5% were discarded. In samples from male individuals, a high heterozygosity rate potentially indicates an incorrect gender or sample contamination. 2) The read counts of OCRs adjacent to FIRRE and XIST, which predominantly show chromatin accessibility in samples from female individuals (81). 3) Read counts in OCRs on chromosome Y outside the pseudoautosomal regions.

Metrics used in quality control

The following quality control metrics were used for all ATAC-seq samples: the total number of initial reads; the number of uniquely mapped reads; the fraction of reads that were uniquely mapped; further mappability-related metrics from the STAR aligner; GC content, insert and duplication metrics from Picard; the rate of reads mapping to the mitochondrial genome; the PCR bottleneck coefficient (PBC), which approximates library complexity as uniquely mapped non-redundant reads divided by the number uniquely mapped reads; the relative strand cross-correlation coefficient (RSC) and the normalized strand cross-correlation coefficient (NSC), which are metrics that use cross-correlation of stranded read density profiles to evaluate the sample quality independently of peak calling; and, finally, the fraction of reads in peaks (FRiP), which is the fraction of reads that fall in called peaks, the fraction of reads in only the blacklisted peaks (wgEncodeDacMapabilityConsensusExcludable and DukeMapabilityRegionsExcludable: https://raw.githubusercontent.com/mills-lab/svelter/master/Support/GRCh38/Exclude.GRCh38.bed), and the ratio between these two metrics (for these metrics the consensus set of peaks was used). The main quality metrics are provided in Data S3.

Quality control and peak calling

On average, 29.1 million sequenced paired-end reads were obtained for each sample. Since FANS sorted nuclei were used, as opposed to whole cells, only a low fraction of reads mapped to the mitochondrial genome (mean 1.94% of the uniquely mapped reads). In our initial dataset of 773 libraries, 46 libraries had a technical or biological replicate. We decided to keep the replicates with the highest number of uniquely mapped reads (resulting into the removal of 39 libraries) or the replicate with the highest FRiP when the number of uniquely mapped reads were comparable (resulting into the removal of additional 7 samples). We also excluded libraries that had low FRiP (less than 3% in a peakset created from one hundred randomly selected samples by requiring a peak to be called in one or more of the sample BAM-files), low mappability (less than 50%), low GC content (less than 85% of cell type median, i.e., 42.52% and 46.24% for neuronal and non-neuronal libraries, respectively), low final read count (less than 5,000,000) or high PMI (more than 48 hours). The threshold of cell GC content was set empirically by testing all values between 75-95% of the cell-type median (with step of 5%) and observing changes in the clustering analysis, as samples with low GC frequently were outliers in the MDS analyses. When combining all these filters, we ended up with 636 samples, out of which none was an outlier in the MDS clustering analysis (Data S1, fig. S2, S3).

For downstream analyses, we split samples into neuronal and non-neuronal datasets. There were two main reasons for this split: 1) The differences in chromatin accessibility due to cell type were vastly greater than the effect of AD phenotypes, and we feared this could lead to subtle biases in the AD phenotype analyses. 2) In analyzing differentially accessible chromatin, we were unable to correct for the effect of markedly different chromatin compositions of these cell types. In particular the different cell types had different proportions of highly accessible OCRs (primarily promoters) and lowly accessible OCRs (primarily non-promoters).

Peak Calling

We subsequently called peaks using the model-based Analysis of ChIP-seq (MACS, v2.1) (82) per each cell type and brain region separately. To create such peaksets, the samples from the same brain region and cell types were subsampled and merged, creating 4 BAM-files with a uniform depth of 700 million paired-end reads. Subsampling ratios were calculated per each sample individually within those four respective groups (brain region by cell type) to ensure that each of them contributed the same number of reads, regardless of their overall read counts. Using these BAM-files, bigWig files were created and peaks were called with the same parameters as described in (20), but using an FDR threshold of 0.01. After removing peaks overlapping the blacklisted genomic regions, 206,701 and 353,722 peaks remained for neuronal and non-neuronal datasets, respectively.

Hi-C data analysis

Hi-C data were aligned using the HiC-Pro strategy (83). Briefly, paired-end reads were mapped independently to human genome hg38 using bowtie2 in stringent mode with parameters ("`--very-sensitive -L 20 --score-min L,-0.6,-0.2 --end-to-end`") (84). Then the chimeric reads that failed to align were trimmed after ligation sites (MboI "GATCGATC") and mapped to the genome. All the aligned reads from two ends were then merged based on reads name and mapped to MboI restriction fragments using hiclib package (85). After that, self-circles, dangling ends, PCR duplicates, and genome assembly errors were discarded. Samples of the same cell type were merged. We binned the interaction matrix at different resolutions and corrected with iterative correction (ICE) for downstream analysis.

Chromatin loops were called with HICCUPS (86) for two different cell types independently. First, we converted the filtered interaction files into juicer format with juicertools. Chromatin loops were called using juicer HICCUPS with bin sizes iterated from 10kb to 25kb by 1kb intervals and parameters "`-k VC_SQRT -p 1 -i 3`". Only reproducible loops were retained and the highest resolution of the overlapping loops were used.

Topological associated domains (TADs) were identified with Topdom (87) at 10K resolution and 200kb window size.

For compartment analysis, first, we calculated the genome-wide correlation matrix at 200Kb resolution with only interchromosomal interactions. Then the first eigenvector of the correlation matrix was obtained and corrected the sign to have a positive correlation with GC content and gene density. The signs of the eigenvector were used to assign the genome into compartment A and B.

Analysis of differentially accessible OCRs of AD-related phenotypes

To identify OCRs showing differential accessibility in AD-related phenotypes, we evaluated the accessibility statistically. Here, chromatin accessibility was estimated as the number of ATAC-seq reads that overlapped a given OCR in a given sample, calculated using RSubread (v1.22.0). The more overlaps seen with an OCR, the more accessible the OCR was considered to be. The subsequent analyses encompassed the following steps:

*Read count and OCR filtering*: The starting point here was a sample by OCR matrix (separately for neuronal and non-neuronal samples) of read counts generated as described in the previous section. From these matrices, OCRs that were lowly accessible were excluded by only keeping OCRs that had at least 2 counts per million reads in at least 10% of the samples. This removed 38,092 and 1,581 neuronal and non-neuronal OCRs, respectively, and resulted in final read count matrices of 323 neuronal samples by 315,630 OCRs and 313 non-neuronal samples by 205,120 OCRs. Next, the read counts were normalized using the trimmed mean of M-values (TMM) method (88).

*Exploration of covariates and model selection*: To explore the effect of sample level technical and biological covariates, a principal component analysis (PCA) was first performed on the normalized read counts to identify high-variance components explaining 1% or more of the variance in chromatin accessibility. We then assessed the correlation of covariates with the aforementioned PCs and selected covariates that showed a significant correlation with one or more PCs at an inclusive FDR cut-off of 0.2 as potential covariates to be used in the analysis of differential chromatin accessibility. This encompassed 63 covariates including FRiP, GC content metrics, mapping metrics, insert metrics, predicted cell type ratios, ethnicities, the rate of reads mapping to the mitochondrial genome, PBC, RSC, and barcode. These covariates were then evaluated as described in the following sections.

As a starting point for modeling chromatin accessibility, a base model was chosen with the variables "brain region by diagnosis status" (2x2=4 levels) and "sex" (2 levels). The reason for including "sex" as a covariate, was that it is known to have a large effect on a few OCRs mostly located on the gonosomes. To evaluate which covariates had to be included to have a good average model of OCR accessibility, we applied an approach based on the Bayesian information criterion (BIC). Here, it was tested, for each additional covariate, how many OCRs showed an improved BIC score minus how many showed a worse BIC score when the covariate was included in the linear regression model compared to when it wasn't. A covariate was then required to improve the mean BIC per OCR by at least 5 for it to be included in the final model. Initially, 53 numeric covariates were evaluated in this way. Compared to the base model, GC_coverage_20-39 (i.e., normalized coverage over each quintile of GC content ranging from 20 – 39), showed the largest and a very pronounced improvement in the fit of the model as it improved a net of 92.5% and 91.2% of the neuronal and non-neuronal OCRs. After this variable was added to the base model, the remaining covariates were tested again. While one more covariate was found for the non-neuronal dataset (FRiP, fraction of reads in peaks), no other covariate fulfilling the criteria for inclusion was found for the neuronal dataset. Subsequently, 15 categorical covariates were considered for inclusion due to the higher number of degrees of freedom of each covariate. None of these met the BIC inclusion criterion. Finally it was considered if the selected numeric covariates (GC_coverage_20-39 for both neuronal and non-neuronal dataset and FRiP only for non-neuronal dataset) affected chromatin accessibility as a quadratic term by testing the squared variable for inclusion. None meet the BIC criterium for inclusion so no additional covariates were added to the model. Thus, the definite models of chromatin accessibility included three (neuronal dataset) or four (non-neuronal dataset) variables: brain region by diagnosis (4 levels), sex (2 levels), GC_coverage_20-39 (numeric) and FRiP (numeric; just for non-neuronal dataset). These models jointly encompassed 6 (neuronal dataset) or 7 (non-neuronal dataset) degrees of freedom (fig. S12).

*Statistical Analysis of differences in chromatin accessibility*: The aforementioned normalized read counts were modeled with the `voomWithQualityWeights` function from the limma package (v.3.38.3) (89). This function utilizes both sample-level and observational-level weights. Voom first residualizes the read counts and fits a mean-variance function across all OCRs to properly address the fact that more accessible OCRs (e.g. those with higher log normalized read counts) show lower variance. The observation level weights are then set as the inverse of the estimated variance. Subsequently, the sample weights are similarly estimated and used to calculate a final set of weights. The normalized read count matrices from `voomWithQualityWeights` were then modeled for each cell type and phenotype (i.e., AD/Control, BBScore, CDR, and Plaque mean) by fitting weighted least-squares linear regression models estimating the effect of the right hand side variables on the accessibility of each OCR:

```
neuronal chromatin accessibility ~ brain region:<phenotype> + sex +
GC_coverage_20_39
non-neuronal chromatin accessibility ~ brain region:<phenotype> + sex +
GC_coverage_20_39 + FRiP
```

To define the sets of differentially accessible OCRs for all combinations of cell types and brain regions and AD-related phenotypes, we looked at contrast between Normal/Low and Severe categories as defined in Data S2. Since both neuronal and non-neuronal datasets often contained two samples from the same individual (one from each of the two brain regions), we ran the differential analysis using the "dream" method from variancePartition package (v1.17.9) (21,70). Dream properly accounts for correlation structures in repeated measures and, thus, avoids inflating the false discovery rate.

As an alternative approach to the differential analysis, we also assessed the difference in the accessibility of OCRs between cases and controls using $\pi_1$ metrics of statistical significance based on estimates of the proportion of true non-null tests, $\pi_1$ (90). The $\pi_1$ (which equals to $1 - \pi_0$) is an estimate of the fraction of OCRs that are differentially accessible between two groups; "1" corresponds to all OCRs estimated to have differential accessibility, whereas "0" corresponds to none of the OCRs having differential accessibility. To calculate this metric, we used *P*-values from differential analysis as an input of the 'propTrueNull' function of the limma package.

Next, adjusted matrices of chromatin accessibility were created for neuronal and non-neuronal samples where the effects of sex, GC content, and FRiP (in case of non-neuronal dataset) were removed. This residualization was done by subtracting the estimated effect of the aforementioned variables in the read count matrix and, thus, retaining just the effect of the brain region and AD case/control status.

Analysis of differentially expressed genes of AD-related phenotypes

The analysis of differential gene expression followed the same approach as the analysis of chromatin accessibility with the the following differences:

*Read count and OCR filtering*: The initial read count matrix of the MSBB RNA-seq dataset (77) consisted of 58,347 genes quantified for 940 samples. From this matrix, we removed (a) duplicated samples (preferentially keeping the samples with the highest RIN values), (b) samples with an rRNA rate > 0.5%, (c) samples with a RIN < 3, (d) samples with genotype mismatches. We also removed genes that were lowly expressed by only keeping genes with at least 1 count per million reads in at least 10% of the samples. These filters led to the exclusion of 107 samples and 37,271 genes. The final read count matrix of 833 samples by 20,709 genes was normalized using the trimmed mean of M-values (TMM) method (88). These genes are mostly protein-coding (74.2%), followed by antisense (7.1%), different types of pseudogenes (7.6%), lncRNA (5.4%), and various other biotypes (<6%).

*Deconvolution of RNA-seq cell type composition*: Cell type composition of RNA-seq samples was estimated using dTangle (v2.0.9) (91), a method built on the linear mixing model of linear-scale expressions of known marker genes. We derived marker genes and reference mixture from single-cell transcriptomics atlas of the human brain (92). Here, only frontal cortex cells were used and we defined five brain cell types: glutamatergic neurons, GABAergic neurons, astrocytes, oligodendrocytes and microglia. The ability of our reference panels to correct for differences in cell type composition was confirmed by comparing the predicted ratios of glutamatergic neurons in AD case and control samples. As expected, AD cases showed the most significant loss of glutamatergic neurons for EC, which is the most vulnerable human brain region in AD, followed by STG, IFG, and FP (figs. S17 and S18).

*Exploration of covariates and model selection*: The following covariates were selected by BIC method to be added to the base covariates, i.e. region by diagnosis status" (4x2=8 levels) and "sex" (2 levels): GC coverage 20-39%, GC coverage 40-59%, GC dropout, mean GC content, fraction of reads aligned to the multiple/too many loci, fraction of intronic reads, RIN, batch (6 levels). After selecting these technical covariates, we modeled the normalized read counts using voomWithQualityWeights and we estimated cell type composition per each sample upon the residualized normalized count matrix. Then, we launched the second round of BIC method that selected three out of five variables of estimated cell type ratios, i.e., fraction of astrocytes, microglia and glutamatergic neurons. In both runs of the BIC method, we required that at least 5% of the peaks showed a change of 4 in the BIC score, corresponding to "positive" evidence against the null hypothesis (93). The relatively high number of covariates selected here compared to the ATAC-seq analysis can, in part, be explained by the RNA-seq being derived from homogenate tissues as opposed to FANS-sorted nuclei. Therefore, the model needed to cope with (a) variability in neuronal / non-neuronal composition between individual dissections, (b) loss of glutamatergic neurons as a result of AD progression. The final model jointly encompassed 25 degrees of freedom (fig. S16).

Annotating ATAC-seq open chromatin regions

*Genes and genomic Context*: The Ensembl 95 genes were used for all analyses in this paper. Further, ChIPSeeker (94) (v.1.18.0) was used to assign genomic context and the closest gene for all ATAC-seq OCRs. For ChIPSeeker, a transcript database was created using GenomicFeatures (95) (v.1.14.8) and the Ensembl genes. Finally, the genomic contexts were defined as promoter (+/- 3kb of any TSS), 5'-UTR, 3'-UTR, exon, intron, and distal intergenic.

*Overlap with previously published epigenomic annotations*: To compare the ATAC-seq derived OCRs in this study to previously reported open chromatin (from REMC (18,96), Cancer Atlas (19), and the Brain Open Chromatin Atlas) the overlap was calculated using the Jaccard index. Here, the Jaccard index was taken as the intersection of base pairs divided by union of base pairs. For this, only the previously known OCRs were considered and not the novel peaks identified in this paper, although the results were similar. For REMC, the imputed datasets were used due to their broader scope and higher quality (96), and the samples were grouped as previously described (20) to reduce dimensionality. For the Cancer Atlas, the Glioblastoma Multiforme, and Low Grade Glioma were considered brain samples. For the Brain Open Chromatin Atlas, only cortical samples were considered by removing amygdala, insula, hippocampus, mediodorsal thalamus, nucleus accumbens, putamen, and the primary visual cortex.

*Overlap with previously published cell-marker genes*: To asses the cell specificity of neuronal and non-neuronal OCRs, we merged all our OCRs, and, among these, the OCRs overlapping only OCRs from one cell type were considered specific to that cell type. Subsequently, we tested the overlap of those peaks with cell specific marker genes (16,97) using the gene set enrichment approach detailed in the next section, but using a Fisher test.

Gene set enrichment analyses of open chromatin regions and gene *P*-values:

For gene sets, we used the MSigDB 7.0 gene sets (71) of sizes 10 to 1000 genes. For ATAC-seq, we used the GREAT approach to assign OCRs to genes (98,99). Using this approach, the gene sets were restructured into "peak sets" consisting of the union of OCRs inferred to regulate one or more genes in the respective gene sets. This was then used as an input to the CameraPR gene set analysis (72). For RNA-seq, the CameraPR function was used directly.

To conduct gene set enrichment analyses on the GWAS data, MAGMA v 1.07b was used on hg19, but, as is customary, excluding the broad MHC region due to its extensive linkage disequilibrium and complex haplotypes (*100*), as well as the ApoE region due to its extreme signal and complex haplotypes (*29*). Further, the genes were padded by 35kb upstream and 10kb downstream to also include genetic variants in the regulatory regions (*100*). Linkage disequilibrium (LD) was estimated from the European panel of 1000 Genome Project phase 3 (*101*).

To illustrate the top genes within each pathway, we needed a *P*-value for each gene in each assay. These were directly outputted for RNA-seq from *dream* and for GWAS from MAGMA. For ATAC-seq, we required direct overlap of the OCRs with one or more TSSs of the gene, to here only highlight OCRs that with fairly high confidence could be assigned to the gene.

Overlap of open chromatin regions with common disease and trait-associated genetic variants

To examine the role that OCRs identified in this paper might play in various diseases and traits, we tested if the OCRs were enriched in common trait associated genetic variants from a selection of GWAS studies. For this, LD-score partitioned heritability (v.1.0.0) (*30*) was used. In LD-score partitioned heritability, it is tested if common genetic variants located in genomic regions of interest explain more of the heritability than variants not in the regions of interest, while correcting for the number of variants in either category. Additionally, the LD-score approach enables one to correct for potential biases from the general genetic context of the genetic regions of interest. This is done using a baseline model of general genomic annotation (such as coding regions and conserved regions) and hence enables one to assess enrichment above and beyond what is expected from the general genetic context of the genomic regions in question. From this regression, a *P*-value as well as a regression coefficient is outputted. To enable comparisons of the regression coefficients across traits with a wide range of heritabilities, we chose to normalize it by the per-SNP heritability and named this adjusted metric the "heritability coefficient". This is not the same as the "enrichment" also outputted by the software, since the heritability coefficient takes the aforementioned baseline into account and the "enrichment" does not. We included the baseline model in the analyses of this paper and we used the approach on a selection of AD & AD-coheritable GWAS traits believed to involve brain function (102-103) and some well powered studies of immune traits (*110*). For the AD GWAS, we removed the APOE effect in the model by excluding SNPs around the APOE gene. For the traits, we used the European only version of the summary statistics when available. As a consequence all GWAS results were based on individuals of European ancestry. The broad MHC-region (hg19:chr6:25-35MB) was excluded due to its extensive and complex LD structure, but, otherwise, default parameters were used for the algorithm. We ran LD-score- analyses only with sets of OCRs covering 0.05% or more of the human genome.

Variance component modeling of gene expression

A variance component analysis was used to examine how much gene expression variability could be correlated to patterns of chromatin covariance (Fig. 2). To implement such a model, we followed an implementation suggested by a previous report (*69*). First, we modeled negative binomial distribution of RNA-seq count data by a variance stabilizing transformation (*111*) (vst;

varistran R package (v.1.0.4)). Then, for each gene represented by vst-normalized vector g, we considered the following variance component model:

$$Y_g' = N(0, P\sigma_p^2 + E\sigma_e^2 + I\sigma_i^2 + U\sigma_u^2)$$

where P and E are sample-sample covariance matrices of chromatin accessibility in promoter (OCRs overlapping region within 1kb from the transcription start site) and enhancer regions (OCRs overlapping region within 1-100kb from the transcription start site), $I\sigma_i^2$captures per-individual covariance matrix and $U\sigma_u^2$ is the noise term. The values of $\sigma_p^2$, $\sigma_e^2$, $\sigma_i^2$, and $\sigma_u^2$, were estimated by the average information restricted likelihood estimation (AIREML; gaston R package (v.1.5.5)). We used residualized count matrices of OCRs where the effect of technical covariates was regressed out. For clarification, this approach does not model the relationship of each gene to its own promoter/enhancer OCRs but to the overall status of all enhancers/promoter OCRs. The principle of this analysis is also summarized in fig. S6.

Prediction of enhancer-gene interactions

We used Activity-by-contact model (ABC, v.0.2) (25) to construct a comprehensive regulatory map of enhancer-promoter (E-P) interactions in neuronal and non-neuronal cell types of the two investigated brain regions (STG and EC). This model requires: (1) contact frequency between putative enhancers and promoters of regulated genes; and (2) enhancer activity data. Contact frequency matrices were generated from neuronal and non-neuronal Hi-C datasets composed of eight post-mortem human brains. Here different neocortical regions (dorsolateral prefrontal cortex, orbital frontal cortex, and anterior prefrontal cortex) were profiled across multiple donors aged 34-103 years. Enhancer activity data was represented by the cell type and brain region specific ATAC-seq signal (current study) and the H3K27ac ChIP-seq signal. ChIP-seq data was generated in a subset of ten controls (STG and EC; age of donors ranged between 61-103 years). In accordance with the authors' directions, we filtered out predictions for genes on chromosome Y and lowly expressed genes (genes that did not meet inclusion criteria in our RNA-seq dataset). We used the default threshold of ABC score (a minimum score of 0.02) and the default screening window (5MB around the TSS of each gene).

To perform an unbiased comparison of $OCR_{ABC}$ (i.e., OCRs with a predicted regulatory link to at least one gene by the ABC model) and $OCR_{other}$ (i.e., OCRs without any predicted regulatory link by the ABC model), we required comparable distributions of both (1) OCR width and (2) OCR distance to the nearest TSS. In particular, we permitted a difference in both metrics for each matching pair of $OCR_{ABC}$ - $OCR_{other}$ of 5%. Due to this restriction, about 2.3% and 6.5% of neuronal and non-neuronal $OCR_{ABC}$ remained unmatched and were therefore left out of the comparison.

*Overlap with existing epigenomic annotations*: We computed the overlap of $OCR_{ABC}$ and $OCR_{other}$ with chromatin states from Epigenomics Roadmap Project (18,112) using the scaled Jaccard index, obtained by calculating standard deviations after subtracting the mean of the sample (Jaccard index is an intersection of base pairs divided by the union of base pairs). We used the "expanded" chromHMM 18-state (6 histone marks, 98 epigenome model) for seven brain regions, i.e. angular gyrus, anterior caudate, cingulate gyrus, dorsolateral prefrontal cortex, hippocampus, inferior temporal lobe, and substantia nigra. To improve interpretability, we consolidated the 18 states into 9 states as follows: Promoter (TssA, TssFlnk, TssFlnkU, and TssFlnkD), Enhancer (EnhG1, EnhG2, EnhA1, EnhA2, EnhWk), Transcription (Tx, TxWk), Poised promoter (TssBiv), Repressed enhancer (EnhBiv), Repressed (ReprPC, ReprPCWk), Heterochromatin (Het), Repeats (ZNF/Rpts), Low (Quies).

*Enrichment in GTEX eQTL*: We computed the overlap of $OCR_{ABC}$ and $OCR_{other}$ with credible sets of SNPs for eGenes in GTEx brain samples (95% credible set interval) (*27*). We used a genome-wide profile as a background to evaluate whether variants in OCRs had a larger chance to affect gene expression. To rule out a potential bias from the genomic background, we also randomly assigned the positions of the annotated ATAC-seq peaks on chromosomes, and calculated the number of variants overlapping the set of aforementioned credible SNPs (50 permutations were evaluated).

Transcription factor analysis

To assess genome-wide putative chromatin occupancy by transcription factors, we performed footprinting analysis using TOBIAS (v.0.10.1) (*43*) as detailed in the following.

*Motif selection*: To build a broad collection of transcription factor binding motifs for the footprinting analyses, all human transcription factor binding motifs were downloaded from the CIS-BP 1.02 meta-database (*113*), which contained 3,059 motifs. As a lot of the transcription factors were represented by more than one motif in the CIS-BP database, and since the transcription factors within the same transcription factor family generally share binding motifs, we reduced the number by choosing only one motif per transcription factor. With a goal of choosing the best motif for each transcription factor, the motif deemed to be most similar to other motifs for that transcription factor was selected analogously to a majority vote: For each transcription factor, the candidate motifs for that given transcription factor were extracted and all pairwise similarities between these candidate motifs were assessed using TomTom (*114*) based on the following command line parameters (*115*): `tomtom -dist kullback -query-pseudo 0.1 -target-pseudo 0.1 -text -min overlap 0 -thresh 1`. These scores were log-transformed and summed for each candidate motif representing the transcription factor, and the motif showing the lowest score was chosen. This resulted in 431 motifs, representing 798 transcription factors.

*Estimation of genome-wide chromatin occupancy by transcription factors*: The footprinting analysis was performed using four merged BAM files consisting of samples representing the same cell type (neuronal / non-neuronal) & brain region (STG / EC). To enable comparisons between neuronal and non-neuronal binding events, we assessed chromatin occupancy by transcription factors in the merged set of neuronal and non-neuronal promoter OCRs, which contained 77,395 OCRs (1.7% of the genome). We ran the TOBIAS module ATACorrect to correct for Tn5 insertion bias in input BAM files, followed by TOBIAS ScoreBigwig to calculate footprinting scores across OCRs. Then, TOBIAS BINDetect combined footprinting scores with the information of transcription factor binding motifs to evaluate the individual binding positions of each transcription factor and determine whether a given position was bound by a given transcription factor or not for each condition, i.e. cell type and brain region. Finally, TOBIAS PlotAggregate was used to visually compare the aggregated footprints for select motifs.

*Calculation of TF cell type specificity score*: To identify regulatory differences between neuronal and non-neuronal samples, we calculated a cell type specificity score per motif *m*, cell type *c*, and brain region *b*, as follows:

$$S_{m,c,b} = \frac{number\ of\ bound\ TF\ sites\ per\ motif\ "m", cell\ type\ "c", brain\ region\ "b"\ that\ are\ not\ bound\ in\ the\ opposite\ cell\ type}{number\ of\ bound\ TF\ sites\ per\ motif\ "m"\ across\ all\ categories\ (neuron\ and\ non-neuron\ cell\ types\ \&\ STG\ and\ EC\ regions)}$$

To allow comparison with external dataset, we aggregate cell type and brain region score per motif into one value as follows:

$$S_m = (S_{m,neuron,STG} + S_{m,neuron,EC}) - (S_{m,non-neuron,STG} + S_{m,non-neuron,EC})$$

To calculate a concordance with Brain Open Chromatin Atlas (BOCA) (20), we retrieved their sets of motifs present in upregulated neuronal and non-neuronal OCRs together with their fold change (observed / expected ratio). For each motif *m*, we aggregated neuronal and non-neuronal fold change enrichment as follows:

$$BOCA\ S'_m = \frac{Fold\_enrichment_{neuron} + \frac{1}{Fold\_enrichment_{non-neuron}}}{2}$$

$$BOCA\ S_m = (BOCA\ S'_m - 1) \ldots if\ BOCA\ S'_m > 1$$
$$-(\frac{1}{BOCA\ S'_m} - 1) \ldots otherwise$$

*Calculation of TFRN networks and prioritization of TF motifs*: TF regulatory networks (TFRN) capturing TF-to-TF interactions among the 431 TF motifs (nodes) were assembled for each cell type and brain region. To define the directed connections (edges) between two TF motifs, we searched for actively bound transcription factor binding sites in the proximal regulatory regions (< 3 kb from TSS) of 798 TF genes. Due to high intracellular similarity of the TFRN, we merged brain region-specific TFRNs and ended up with 26,976 and 40,348 unique, directed TF-to-TF interactions for neuronal and non-neuronal TFRNs, respectively. Then, we used those networks as an input of the HotNet algorithm (73) to find altered subnetworks containing TF motifs that are highly dysregulated based on transcriptomics or on the GWAS level and are topologically close on an interaction network. We here utilized all four AD-related phenotypes (AD/Control, BBScore, CDR, Plaque mean) from the RNA-seq differential analysis results to derive four layers of transcriptomics weights for TF motifs. GWAS weights were extracted from gene-level *P*-values for Alzheimer's disease from GWAS summary statistics (29) and MAGMA as detailed above. For both transcriptomics and GWAS weights, we applied -log10(*P*-value) transformations. In the case of multiple TF genes per one TF motif, we set the weight to be the highest -log10(*P*-value) and applied the Sidak method for multiple testing corrections (116). Final lists of TF motifs with different TF binding in AD in neuronal and non-neuronal cells (as predicted by HotNet) were filtered out and only TF motifs that participated in at least 50 TF-to-TF interactions were kept.

*Prioritization of TF genes represented by the same TF motif*: Because each TF motif is usually represented by multiple TF genes, we needed to address the challenge of pinpointing phenotypically causal TF genes. For each TF motif, we first identified a set of genes that have its motif actively bounded in the proximal regulatory region (within 3 kb from TSS). We subsetted the expression matrix for those genes and calculated its principal components. Then, we calculated the Pearson correlation of the first principal component with the expression of TF genes belonging to the TF motif under inspection and we discarded TF genes with significance of Pearson correlation above Bonferroni adjusted *P*-value threshold of 0.05. For further prioritization of disease-associated-TFs, we also filtered out those TF genes that did not show differential gene expression in any of four AD-related phenotype comparisons. The whole process is summarized in fig. S26.

Experimental validation of USF2

*Cell line and generation of stable cell line:* Human neuroblastoma cell line SH-SY5Y was purchased from the ATCC (Cat# CRL-2266) and maintained using DMEM (Gibco, Cat# 11995073) supplemented with 10% fetal bovine serum (FBS)(Gibco, Cat# 16000069) and 1% penicillin-streptomycin (PS)(Gibco, Cat# 5140122). To generate stable cell lines overexpressing Usf2, Myc-DDK-Usf2 cDNA was transfected using Lipofectamine 2000 (Invitrogen, 11668019) according to the manufacturer's protocol. For the present study, a stable line for Usf2 was made by limiting dilution and were maintained on 500 μg/ml G418.

*siRNA transfection*: siRNA against human *USF2* (sc-36786) was purchased from Santa Cruz Biotechnology and negative control DsiRNA was purchased from Integrated DNA Technologies (IDT), and used as previously described (*117*). Cells were transfected using Lipofectamine RNAiMAX (Invitrogen, Cat# 13778150) according to the manufacturer's protocol and siRNA for *USF2* at a final concentration of 100 nM.

*Cell lysis*: Cells were washed with ice cold 1X PBS then harvested in ice cold RIPA buffer (20 mM Tris-HCL, pH 7.5, 150 mM NaCl, 1 mM EDTA, 1 mM EGTA, 1% NP-40, 1% sodium deoxycholate) with 1X protease and phosphatase inhibitor cocktail (Thermo, Cat# 1861284). After 30 min on ice, sonicated cells were then centrifuged at 16,000 g for 15 min. The supernatants were used for immunoblotting.

*Gel electrophoresis and immunoblotting*: Samples were mixed with 1X SDS sample buffer (62.5 mM Tris, pH 6.8, 10% Glycerol, 1% SDS, 20 mM DTT, 5% β-mercaptoethanol, and 0.005% Bromophenol blue [BPB]) and incubated 5 min at 100 ˚C, otherwise samples were mixed with 2X urea sample buffer (9.6% SDS, 4 M Urea, 16% Sucrose, 0.005% BPB, and 4.6% β-mercaptoethanol) and incubated 15 min at 55 ˚C for V0 subunits of v-ATPase followed by electrophoresis on Novex™ 4-20% Tris-Glycine gradient gels (Invitrogen, Cat#s WXP42020BOX; WXP42026BOX). Proteins were transferred onto 0.2 μm nitrocellulose membranes (Pall Laboratory, Cat# 66485) and the membrane was incubated overnight in primary antibody (**Data S16**) then incubated with HRP conjugated secondary antibody. The blot was developed using ECL-kits (Invitrogen, Cat# WP20005; Millipore, Cat# WBKLS0500).

*Lysosomal isolation:* Cells were incubated in growth medium containing 10% Dextran conjugated magnetite (Liquid Research LLC, DexoMAG™ 40) for 24 hr, then chased in normal growth media for 24 hr. Cells were washed with 1X PBS then harvested in 4 ml of ice-cold Buffer A (1 mM HEPES, pH 7.2, 15 mM KCl, 1.5 mM MgAc, 1 mM DTT, and 1X PIC). Cells were then homogenized with 40 strokes of a loose-fitting pestle in a Dounce homogenizer then passed through a 23 G needle 5 times. After homogenization, 500 μl of ice-cold Buffer B (220 mM HEPES, pH 7.2, 375 mM KCl, 22.5 mM MgAc, 1 mM DTT, and 20 μM DNase I) was added and samples were then centrifuged at 750 xg for 10 min. The supernatant was then decanted over a QuadroMACS™ LS column (Miltenyi Biotec, 130-042-976) that had previously been equilibrated with 0.5 % BSA in PBS, and then collected non-lysosomal fraction (Flow) to flow through via gravity. The pellet was subjected to re-addition of 4 ml ice cold Buffer A, 500 µl ice cold Buffer B and then re-suspended and re-centrifuged. This second supernatant was also passed over the column and allowed to flow through via gravity. DNase I (10 µl/ml in PBS) was added and the column was then incubated for 10 min and then washed with 1 ml ice cold PBS. Lysosomes were eluted by removing the column from the magnetic assembly, adding 100 µl of PBS for immunoblotting / M1 buffer (10 mM Tris, pH 7.5, 250 mM Sucrose, 150 mM KCl, 3 mM β-mercaptoethanol, and 20 mM $CaCl_2$) for v-ATPase activity and Proton Translocation assay and forced through the column using a plunger.

*v-ATPase activity assay:* Lysosome-enriched fractions (4 µg) were mixed with 0.052% $NaN_3$ for blocking the mitochondrial ATPase activity. The v-ATPase activity measured using ATPase assay kit (Innova Biosciences, 601-0120) according to the manufacturer's protocol. Control samples were measured in the presence of the v-ATPase inhibitor ConA (1 µM) and the experimental values were subtracted accordingly. Absorbance was measured at 650 nm and solutions of $P_i$ were used to generate a standard curve.

*Lysosomal pH measurement:* Procedures were performed as previously described (14). Following the addition of 250 µg LysoSensor$^{TM}$ Yellow/Blue dextran (Molecular probes, L22460) treatments, cells were incubated for 24 hr. The samples were then read in a Wallac Victor 2 fluorimeter (Perkin Elmer) with excitation at 355 nm. The ratio of emission 440 nm/535 nm was then calculated for each sample. The pH values were determined from the standard curve generated via pH calibration samples.

Three-dimensional chromatin interactions from population-level maps of chromatin accessibility

To determine three-dimensional (3D) chromatin interactions and evaluate differential accessibility in 3D interactions associated with AD-related phenotypes, we integrated the decorate pipeline (59) and applied additional statistical tests. Here, we explain the systematic workflow of our pipeline (**fig. S29**).

*Data preprocessing and cis-regulatory domain (CRD) calling*: CRD calling was performed separately on $m_{OCR} \times n_{samples}$ covariate-corrected ATAC-seq count matrices of STG neurons, EC neurons, STG non-neurons, and EC non-neurons, obtained from single-OCR analysis. Before CRD calling, PEER (probabilistic estimation of expression residuals) (118) residualization on each matrix was performed to remove the global effects of covariates, hence, to retain the local correlation structure. PEER-corrected matrices $m_{OCR\_PEERj} \times n_{samples}$ ($j = \{1, 5, 10, 15, 20, 25\}$) from each dataset makes a total of 24 input matrices (6 PEER-corrected STG neurons, 6 PEER-corrected EC neurons, 6 PEER-corrected STG non-neurons, and 6 PEER-corrected EC non-neurons) for CRD calling. CRDs were called on 24 matrices individually using the following R functions from the *decorate* package.

```
CRDlist=runOrderedClusteringGenome(
      mOCR_PEERj × nsamples,OCRgenomic_coords,method.corr="spearman")
CRDClusters = createClusters(CRDlist, method = "meanClusterSize",
      meanClusterSize=c(10, 25, 50, 80,100))
CRDScore=scoreClusters(CRDlist,CRDClusters)
```

*CRD filtering and merging*: From the above step, we obtained 24 *CRDScore* lists containing names of OCRs that are assigned to CRDs, their mean correlation, and Lead eigen factor (LEF). LEF of a CRD is a fraction of variance explained by the first eigenvalue of the correlation matrix *[m × m]* of OCRs located within a CRD. Larger LEF values (i.e. >10%) can be interpreted as strongly correlated OCRs, whereas smaller values correspond to weaker correlations of OCRs located within a CRD. Filtering out the CRDs with weaker correlations is an important step as it substantially reduces the burden of multiple testing in differential CRD analysis.

The optimal value of $LEF_{cutoff}$ was calculated in three steps. In the first step, OCR genomic locations were shuffled in a chromosome-wise manner in $m_{OCR\_PEERj} \times n_{samples}$ matrix to create permuted matrices $m_{Permutation\_i\_OCR\_PEERj} \times n_{samples}$ (where $j = \{1, 5, 10, 15, 20, 25\}$ and $i = <1, 10>$). This step breaks the local correlation structure and retains the global correlation structure. In the second step, *CRD calling* was implemented using the same parameters as explained in the previous section for all 240 matrices (10 permutations × 6 PEER-corrected matrices × 4 datasets; $m_{Permutation\_i\_OCR\_PEERj} \times n_{samples}$). In the third step, $LEF_{cutoff}$ was defined based on the 10% quantile of $LEF_{Permuted}$ vector after combining 10 $CRDScore$ lists obtained from CRD calling on $m_{Permutation\_i\_OCR\_PEERj} \times n_{samples}$ ($i = <1\text{-}10>$)

$$LEF_{Permuted} = [LEF_1, LEF_2, LEF_3, ..., LEF_{10}], where\ LEF_i = N \times 1$$
$$LEF_{cutoff} = Pr(LEF_{Permuted} = threshold), where\ threshold = 1 - 0.10$$

The final table of CRDs was obtained by keeping all CRDs with $LEF_{measured} > LEF_{cutoff}$ in $CRDScore$ list of $m_{OCR\_PEERj} \times n_{samples}$ (**fig. S30A**). Next, overlapping CRDs of different sizes were merged to obtain discrete CRDs for downstream analysis. To decide the optimal number of PEER factors, we measured the number of CRDs called on the input matrix ATAC-seq count matrix residualized by various numbers of PEER factors; we tested {*1, 5, 10, 15, 20, 25*} PEER factors (**fig. S31**). The distributions of LEF, mean absolute correlation of CRDs, and the number of OCRs located within CRDs are shown in **fig. S30B-D**, while the final lists of coordinates of CRDs of STG neurons, EC neurons, STG non-neurons, and EC non-neurons are provided in **Data S17**.

*CRD biological validation*: To validate 3D interactions captured by CRDs with the Hi-C dataset, we used CTCF ChIP-seq peak list from ENCODE human neural cells (*119*). We quantified the density of CTCF sites in 200 bins (each bin size equals to 1kb) around CRD boundaries (**Fig 7D**). In order to quantify how many *in situ* 3D interactions captured by CRDs are within the 3D interactions measured as Topologically Associated Domains (TADs) from cell specific Hi-C experiments (section "Hi-C data analysis"), we measured the number of CRDs overlapping with cell specific Hi-C TADs stratified by number of TADs ($N = \{0,1,2,3,>=4\}$). Next, we measured how many cell-specific Hi-C TADs are within the CRDs stratified by the number of CRDs ($N = \{0,1,2,3,>=4\}$). We measured the correlation of OCRs inside the Hi-C loops and outside the Hi-C loops. Genome wide enrichment of CRDs in A and B compartments was performed using Fisher exact test. Cell specific A and B compartment coordinates (section "Hi-C data analysis") were obtained from Hi-C datasets.

*Differential CRD analysis*: To perform differential CRD analysis for each AD-related phenotype, we used per-OCR differential analysis results (**Data S8**). Here, we show the calculation of *P*-value and log(fold change) for one CRD ($CRD_x$) that is linked to *k* OCRs.

$$CRD_x = [OCR_1, OCR_2, OCR_3, ..., OCR_k], where\ CRD_x\ has\ k\ OCRs$$

$$p.min = min\{Pvalue_{OCR=i};\ i = 1,2,3,\ldots,k\}$$
$$Pvalue_{CRD_x} = 1 - (1 - p.min^k)$$
$$logFC_{CRD_x} = \frac{1}{k}\sum_{i=1}^{k} logFC\ (OCR_i)$$

*Differential gene-CRD analysis*: To map genes to CRDs, we utilized the promoter/enhancer annotations of their respective OCRs. Enhancer OCR-gene links were predicted by "activity-by-contact" model (ABC score ≥ 0.2, **Data S5**). Promoter OCR-gene links were determined based on the distance to the nearest TSS (promoter OCRs need to overlap the genomic region of 2 kb around gene's TSS). To perform differential gene-CRD analysis for each AD-related phenotype, we used per-gene differential analysis results (**Data S11**). Here, we show the calculation of *P*-value and log(fold change) for one CRD (*$CRD_x$*) that is linked to *k* genes.

$$ABC_{Mapped\ CRD_x} = [Gene_1, Gene_2\ , Gene_3, \ldots, Gene_k], where\ CRD_x\ has\ k\ genes$$
$$p.min = min\{Pvalue_{Gene_i};\ i = 1,2,3,\ldots,k\}$$
$$Pvalue_{ABC_{Mapped\ CRD_x}} = 1 - (1 - p.min^k)$$
$$logFC_{ABC_{Mapped\ CRD_x}} = \frac{1}{k}\sum_{i=1}^{k} logFC\ (OCR_i)$$

**Data S1. (separate Excel file)**
Summary characteristics of the ATAC-seq and RNA-seq datasets.

**Data S2. (separate Excel file)**
Classification of samples from ATAC-seq and RNA-seq datasets by various clinical and pathological traits.

**Data S3. (separate Excel file)**
Summary of quality control metrics of the ATAC-seq dataset.

**Data S4. (separate Excel file)**
Gene expression explained by variance component decomposition model that discretize explanatory power of promoters and enhancers OCRs.

**Data S5. (separate Excel file)**
Enhancer-gene links as predicted by Activity-By-Contact (ABC) methods.

**Data S6. (separate Excel file)**
Enrichment of "Enhancer OCRs" and "Other non-promoter OCRs" in eQTLs. "# of variants" indicates how many variants overlapped the annotation. "# of credible" indicates how many credible SNPs overlapped the annotation. "%" indicates the percentage over all SNPs overlapping the annotation that were credible SNPs. *P*-value is for enrichment against the background (where 1% of SNPs were credible) using a Fisher exact test. 50 permutations were done to evaluate potential biases.

**Data S7. (separate Excel file)**
Numbers of differentially accessible OCRs for combinations of cell type & brain regions and AD-related phenotypes.

**Data S8. (separate Excel file)**
Differential ATAC-seq OCRs.

**Data S9. (separate Excel file)**
Fraction of differentially accessible OCRs based on proportion of true tests.

**Data S10. (separate Excel file)**
Overlap between disease-associated OCRs and common genetic variants associated with brain and non-brain related traits using LD-sc. Only sets of OCRs covering at least 0.05% of the genome were tested.

**Data S11. (separate Excel file)**
Differential RNA-seq genes.

**Data S12. (separate Excel file)**
Gene *P*-values across ATAC-seq, RNA-seq, and GWAS

**Data S13. (separate Excel file)**
Cell type specificity score for sets of analyzed transcription factors.

**Data S14. (separate Excel file)**
Reference for experimental validation or computational prediction of cell type specificity of highlighted transcription factors.

**Data S15. (separate Excel file)**
Sets of genes with exclusive transcriptional regulation coming either from neuron or non-neuron TFs.

**Data S16. (separate Excel file)**
Summary information about primary antibodies used for experimental validation of the effect of USF2 knockout and over-expression.

**Data S17. (separate Excel file)**
Genomic coordinates of CRDs with their respective OCRs.

**Data S18. (separate Excel file)**
Differential CRDs and differential genes inside CRDs.

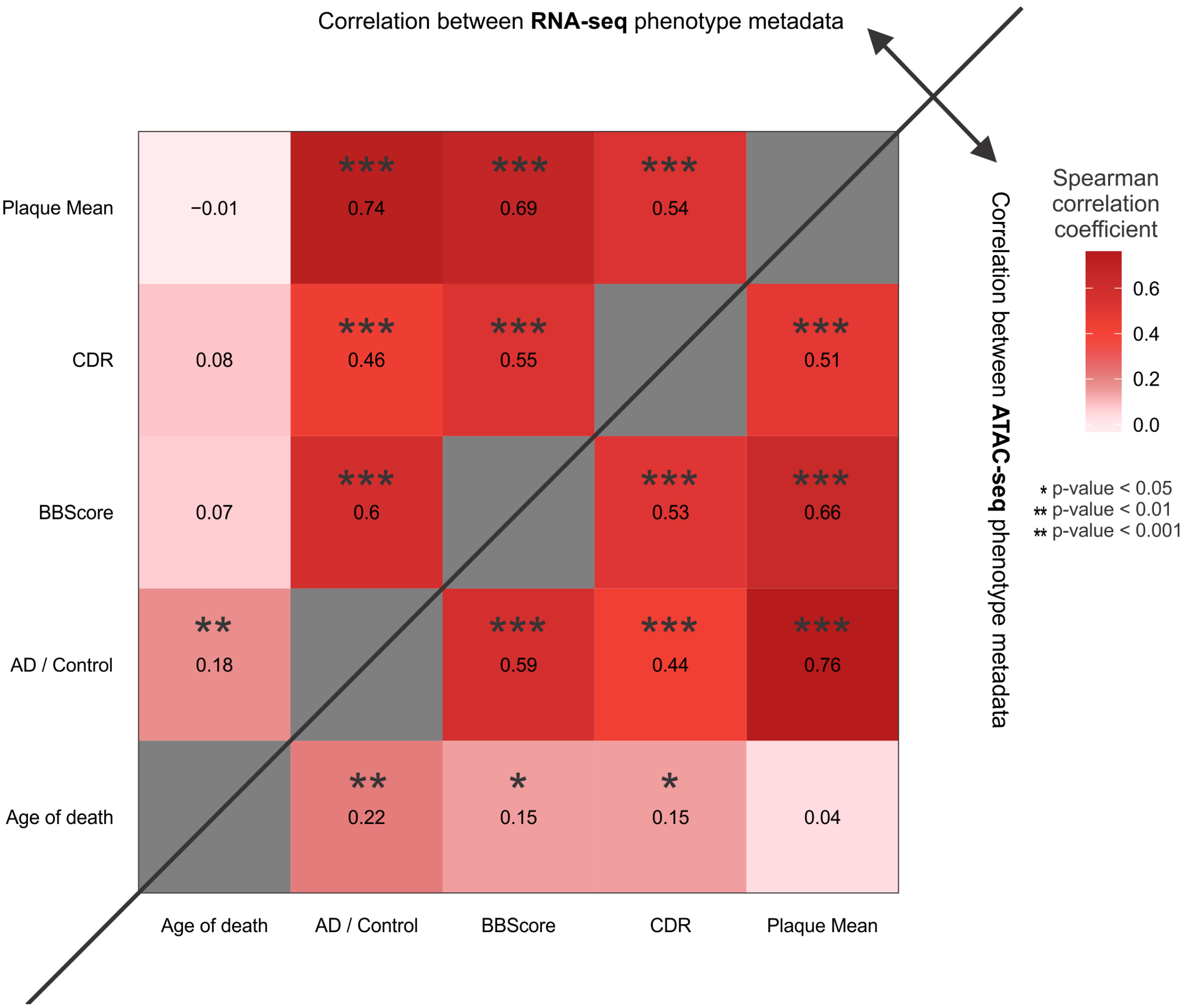


**Fig. S1.**
Correlation among phenotypic variables within RNA-seq (left upper triangle) and ATAC-seq (right lower triangle) cohort. The numbers in each cell indicate the Spearman's rank correlation coefficient between row and column phenotypic attributes. Those correlations slightly differ between ATAC-seq and RNA-seq cohort due to different sets of assayed individual (3% of individuals from ATAC-seq cohort are not present in RNA-seq cohort and, conversely, 34% of individuals from RNA-seq cohort are not present in ATAC-seq).

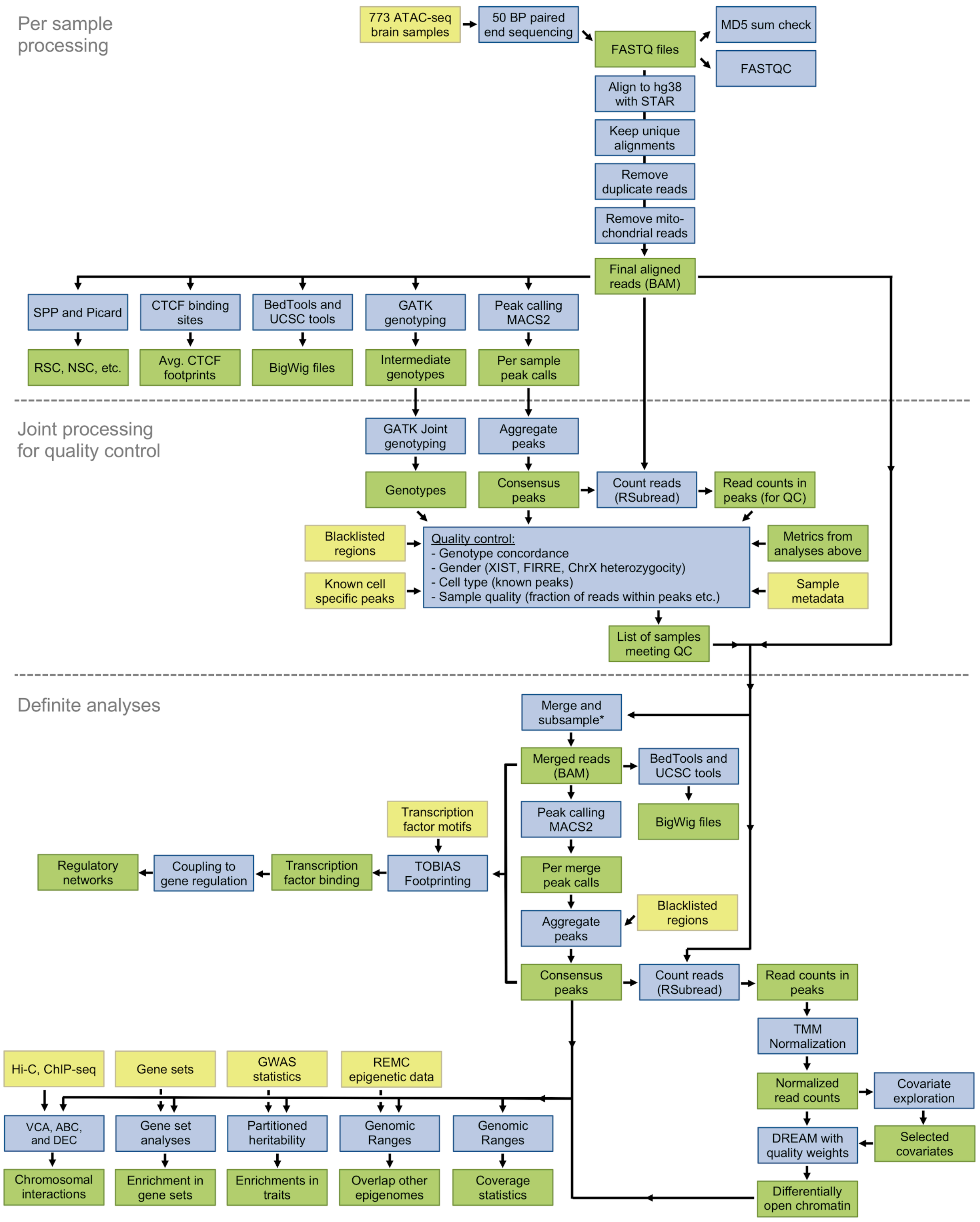


**Fig. S2.**
Flowchart of data processing. There are three different phases to the pipeline: 1) per sample processing, 2) joint processing for quality control, and 3) processing and analyses of samples

surviving quality control. Yellow: input data. Blue: analyses. Green: processed data. REMC: Roadmap epigenomic mapping consortium, VCA: variance component analysis, ABC: activity by contact, DEC: differential epigenetic correlation test.

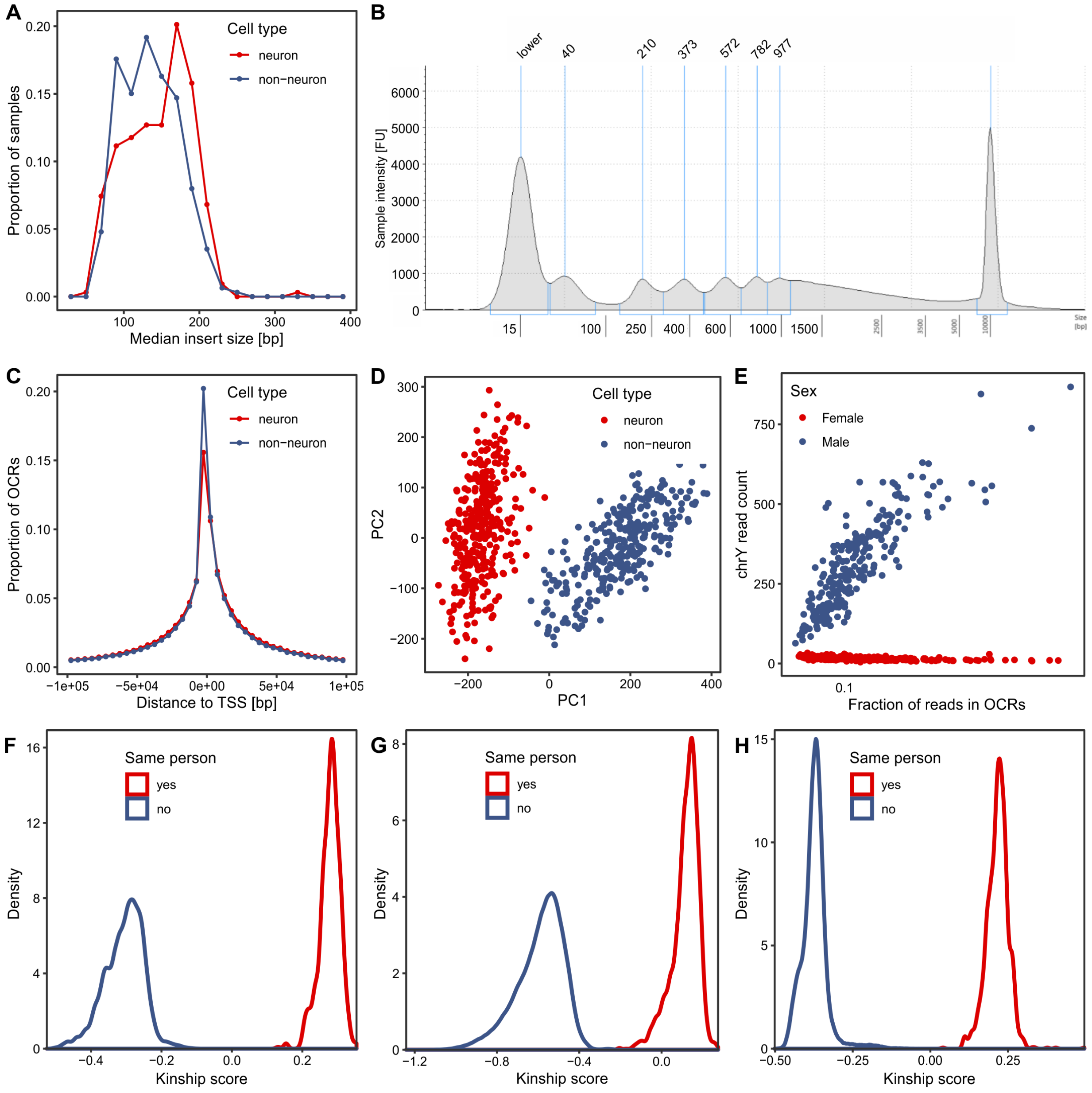


**Fig. S3.**
Quality control metrics. (**A**) Median read insert size distribution of ATAC-seq samples; (**B**) Tapestation profile of typical ATAC-seq library; (**C**) Distance to TSS distribution of OCRs; (**D**) Principal component analysis of chromatin accessibility levels in OCRs; (**E**) Sex check based on the number of reads mapped to chromosome Y; (**F-H**) Genotype checks based on pairwise comparisons of genotypes from ATAC-seq with genotypes from whole-genome sequencing, RNA-seq, and the ATAC-seq samples themselves.

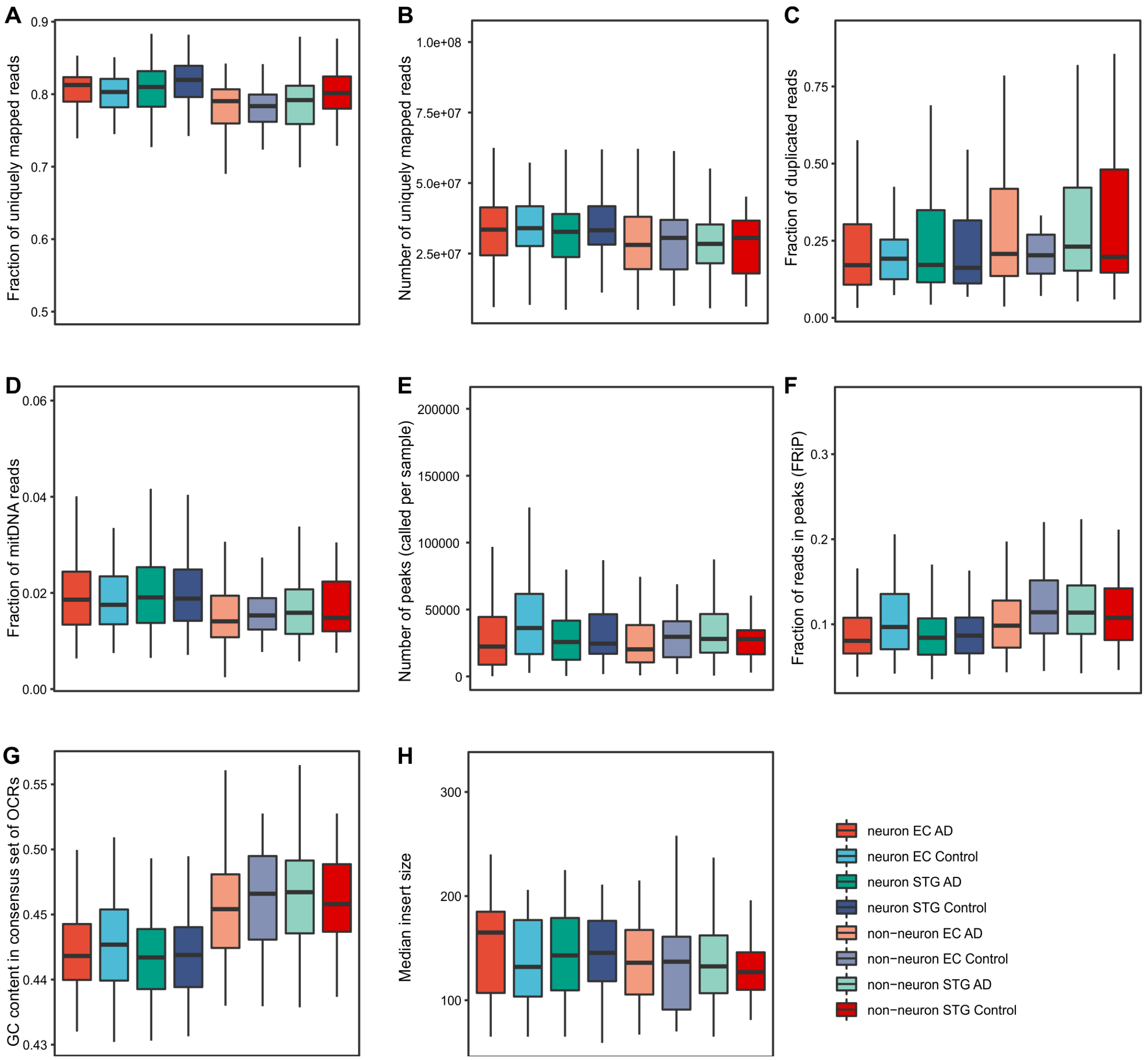


**Fig. S4.**
Quality control metrics stratified by cell type, brain region and diagnosis. (**A**) Fraction of uniquely mapped, non-duplicated, non-chrM paired-end reads compared to all reads in raw sequencing files; (**B**) Number of uniquely mapped, non-duplicated, non-chrM paired-end reads; (**C**) Fraction of duplicated to uniquely mapped paired-end reads; (**D**) Fraction of mitochondrial DNA reads to uniquely mapped, non-duplicated paired-end reads; (**E**) Number of peaks (called per sample); (**F**) Fraction of reads in peaks (FRiP); (**G**) GC-content in consensus set of OCRs; (**H**) Median insert size. The center line indicates the median, the box shows the interquartile range, whiskers indicate the highest/lowest values within 1.5x the interquartile range.

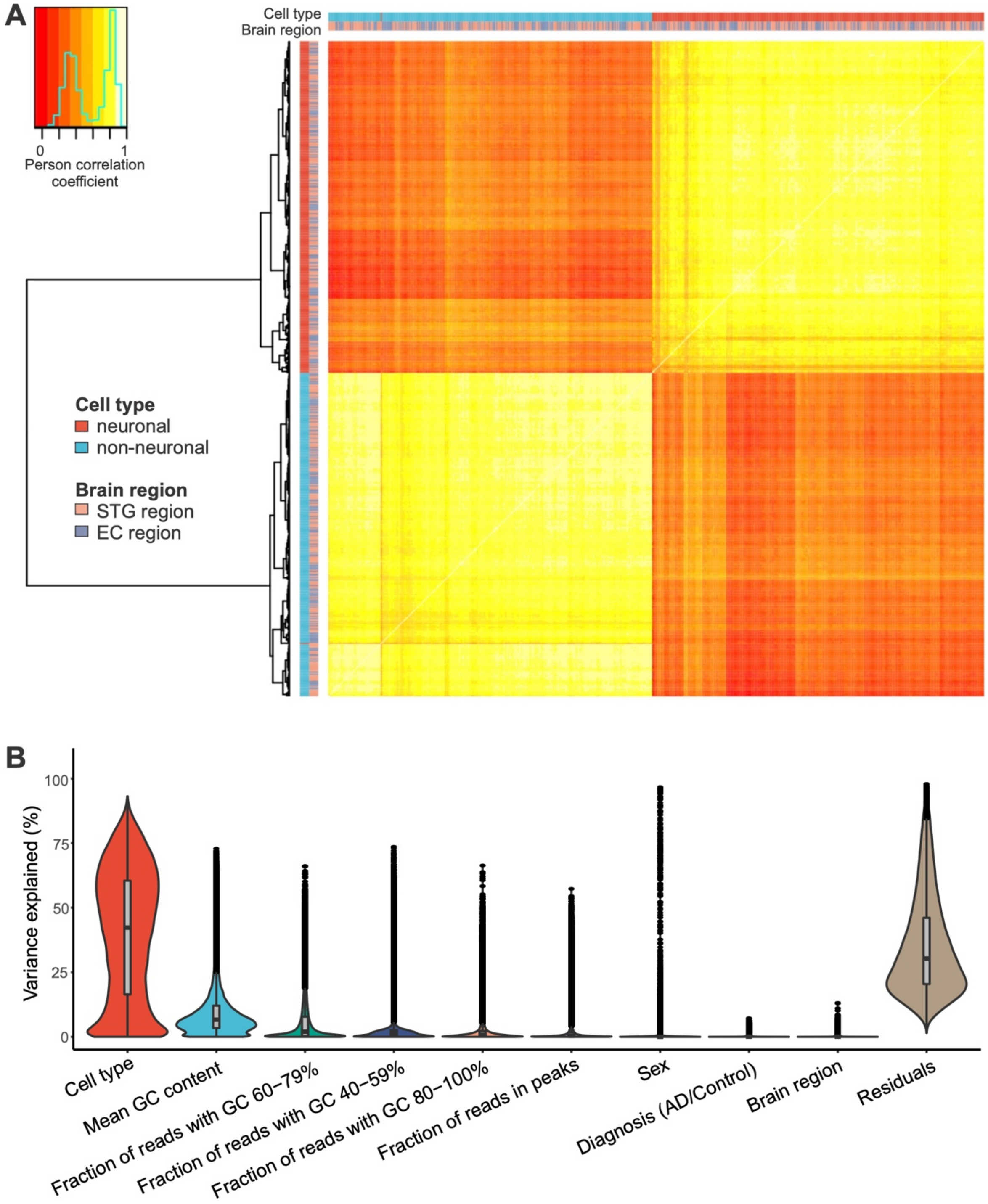


**Fig. S5.**
Cell type is a major source of variation in the ATAC-seq dataset. (**A**) Heatmap of the correlation among ATAC-seq samples, based on the Pearson correlation from OCR read counts. (**B**) Distribution of the fraction of variance explained by each covariate plus residuals across all OCRs prior adjustment for covariates. Covariates were selected by repeated BIC procedure.

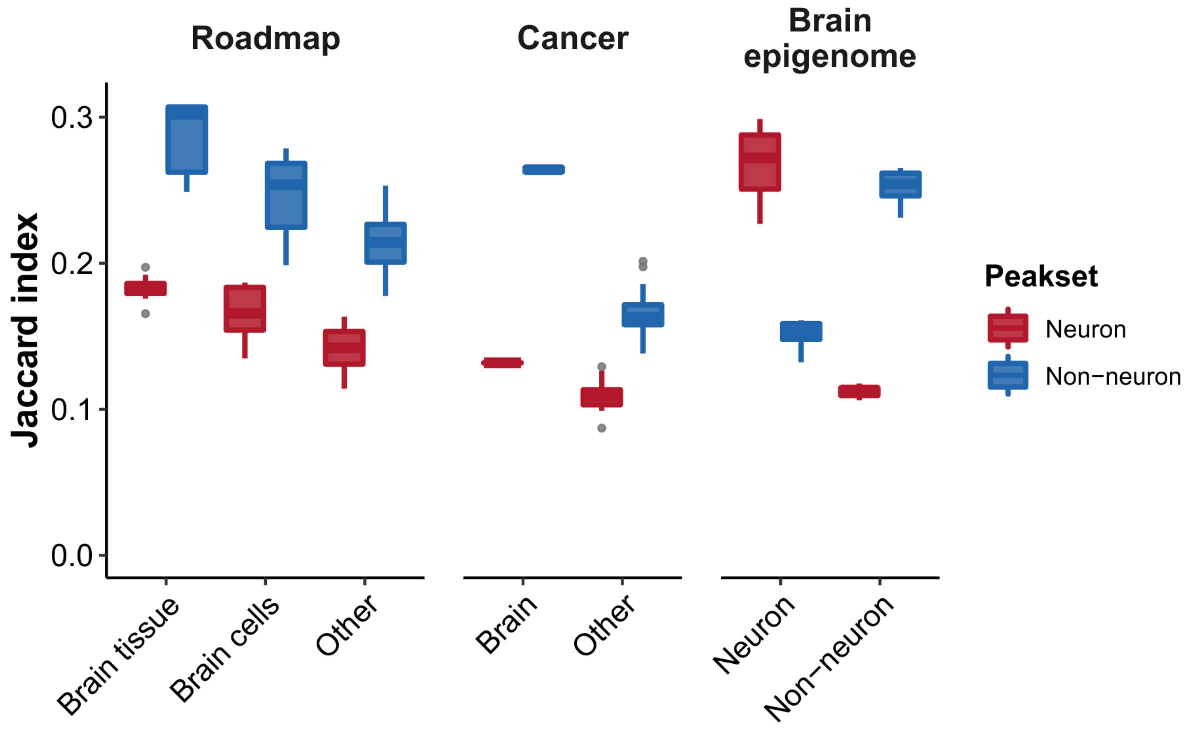


**Fig. S6.**
Jaccard overlap between OCRs identified in this paper with Roadmap Epigenomics Consortium, The Cancer Genome Atlas, and a brain epigenome atlas. For this, we considered only the non-novel OCRs in this study, although results were comparable considering all OCRs aside from the average Jaccard index being smaller. The center line indicates the median, the box shows the interquartile range, whiskers indicate the highest/lowest values within 1.5x of the interquartile range, and potential outliers from this are shown as dots.

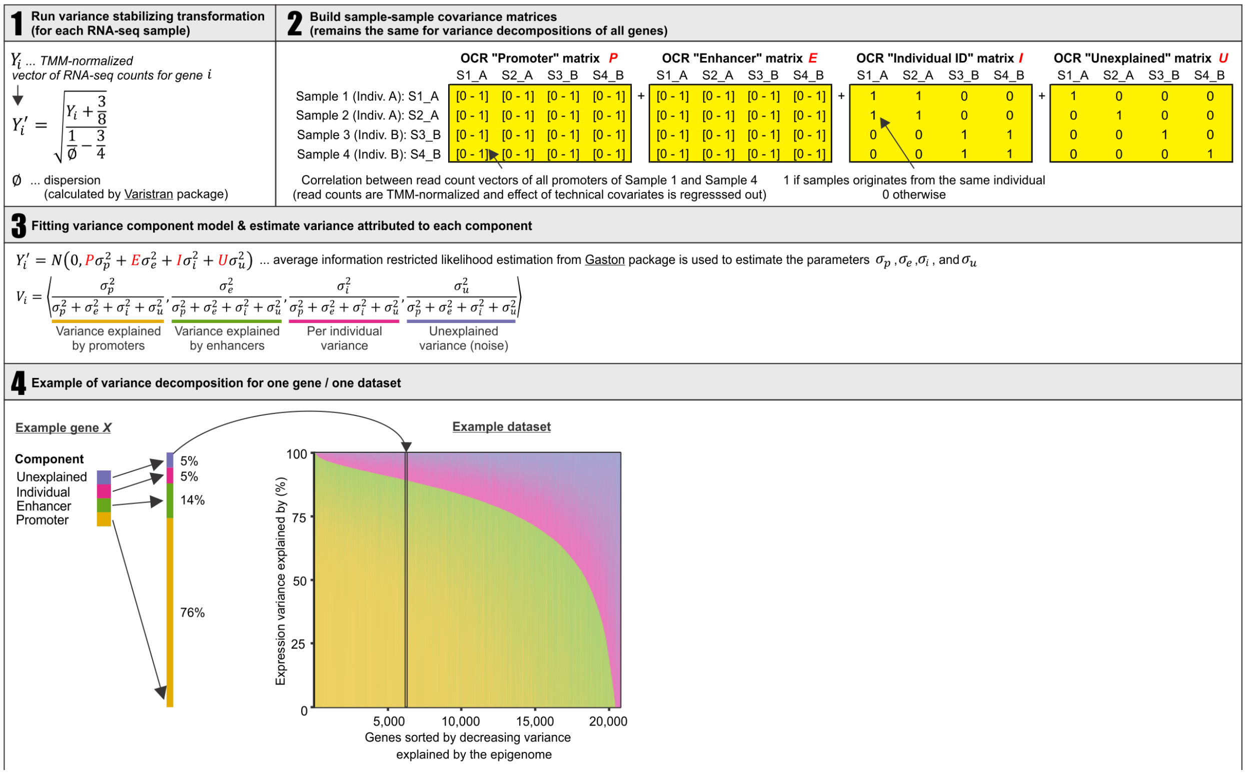


**Fig. S7.**

Workflow of variance component analysis used to explain gene expression variability by the patterns of chromatin covariance. Panel 1 describes the variance stabilizing transformation of RNA-seq count data. Panel 2 depicts ATAC-seq sample-sample correlation matrices computed separately for promoter OCRs and enhancer OCRs, supplemented by “Individual ID” matrix (for capturing inter-individual variance) and Identity matrix (for capturing unexplained variance). Panel 3 describes a variance component model based on Average Information Restricted Likelihood Estimation (AIREML). Panel 4 shows an example of one gene with expression variance explained by chromatin covariance patterns in promoter (76%), enhancer (14%), inter-individual (5%), and noise (5%).

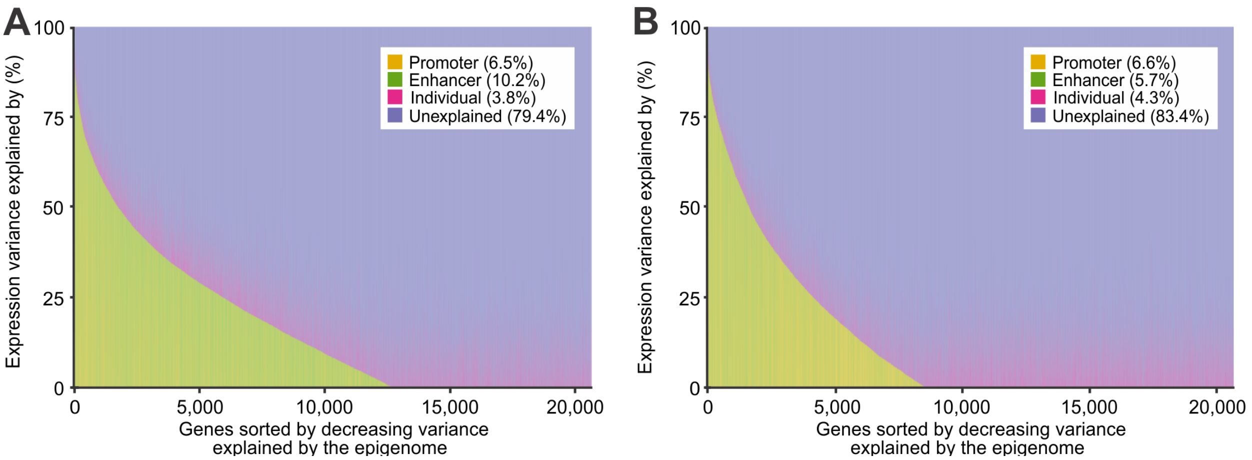


**Fig. S8.**
Variance component analysis of gene expression for permuted dataset, i.e. samples in RNA-seq matrix were shuffled. Neuronal (**A**) and non-neuronal (**B**) results of variance component model indicates that only a minimal gene expression was estimated to be explained by promoter-, enhancer- or individual-level variance with a shuffled dataset (mean variance explained by each component shown in parentheses).

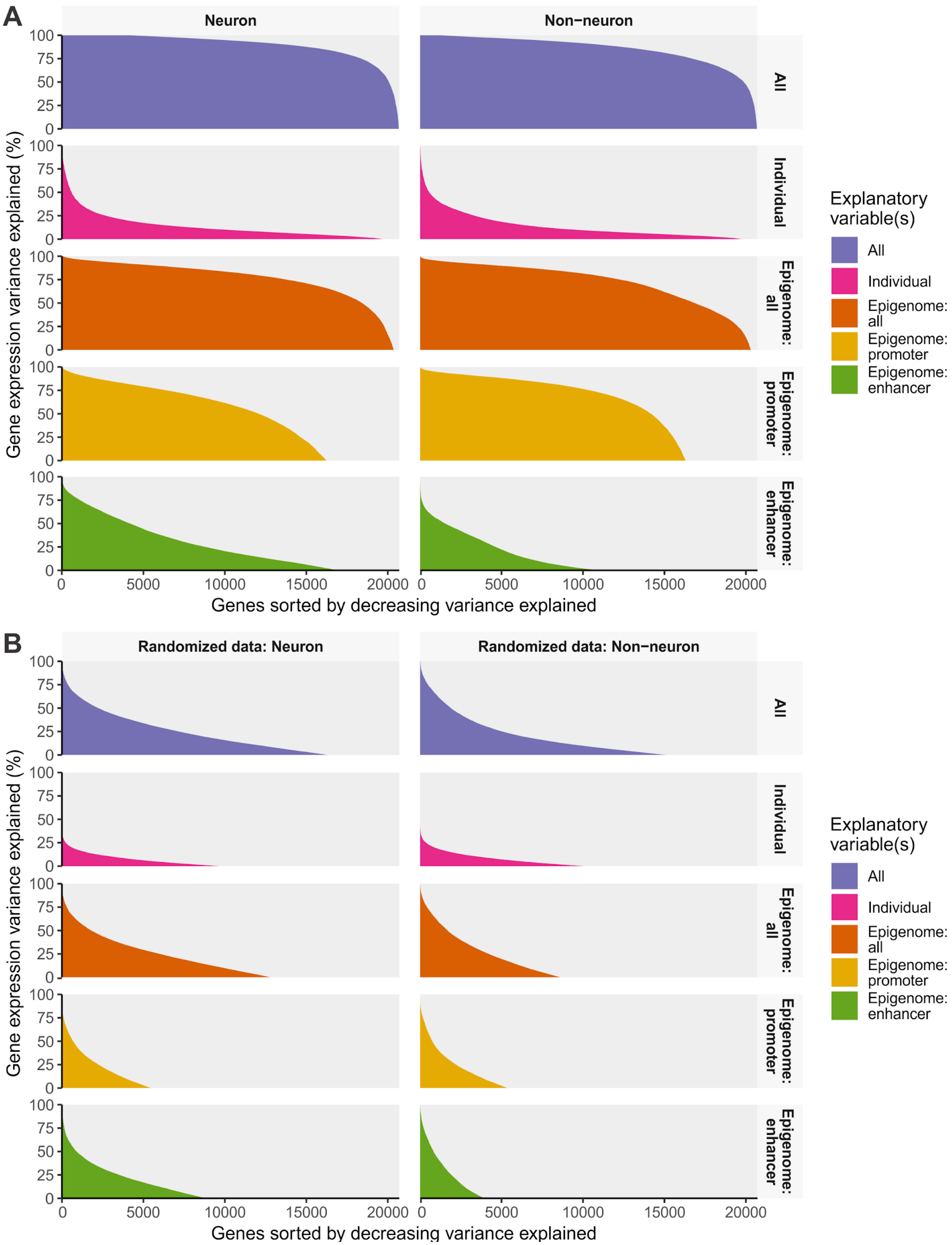
A
Neuron
Non−neuron
All
Individual
Epigenome: all
Epigenome: promoter
Epigenome: enhancer
Gene expression variance explained (%)
Genes sorted by decreasing variance explained
Explanatory variable(s)
B
Randomized data: Neuron
Randomized data: Non−neuron

**Fig. S9.**

Variance in gene expression estimated to be explained by the neuronal or non-neuronal epigenome as well as the individual brain using (**A**) actual data (**B**) shuffled data. The genes in each subpanel are sorted by the amount of variance explained by the variable(s) in question, and are thus different amongst the subpanels. E.g. a gene might have a high degree of variance explained by the epigenome but a small amount explained by the individual. “All” is the sum of individual, promoter and enhancer. “Epigenome: all” is the sum of promoter and enhancer.

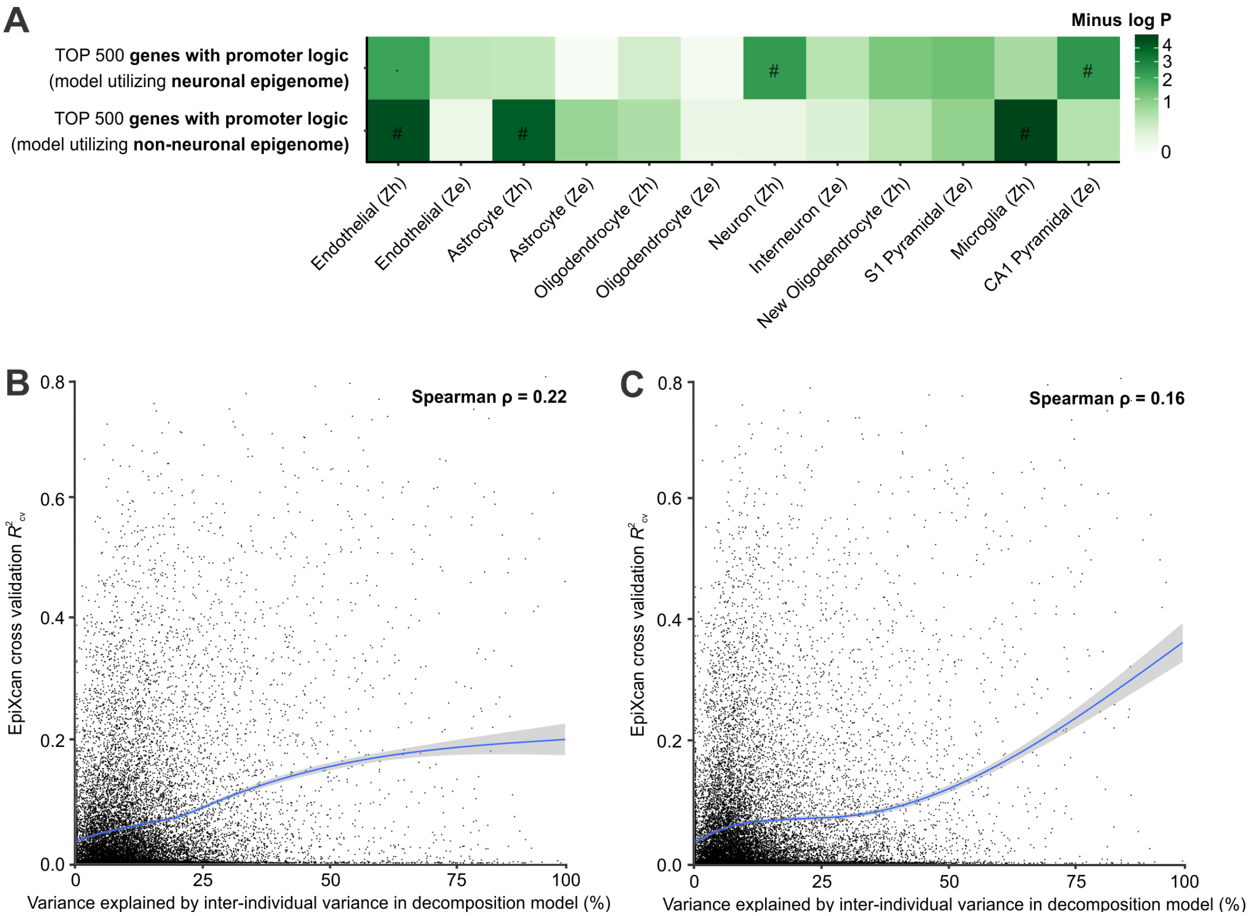


**Fig. S10.**
Additional analysis of the results of variance decomposition model. (**A**) Overlap between the top 500 genes with the highest proportion of variance explained by promoters/enhancers and cell-specific marker genes from (16,97). "#": FDR < 0.05 " · ": nominal *P*-value < 0.05. (**B-C**) Per gene relationship between variance explained by inter-individual variance in (B) neuronal and (C) non-neuronal ATAC-seq dataset and EpiXcan cross validation R-squared ($R^2_{CV}$). The solid smooth blue lines are LOESS (local regression) fits.

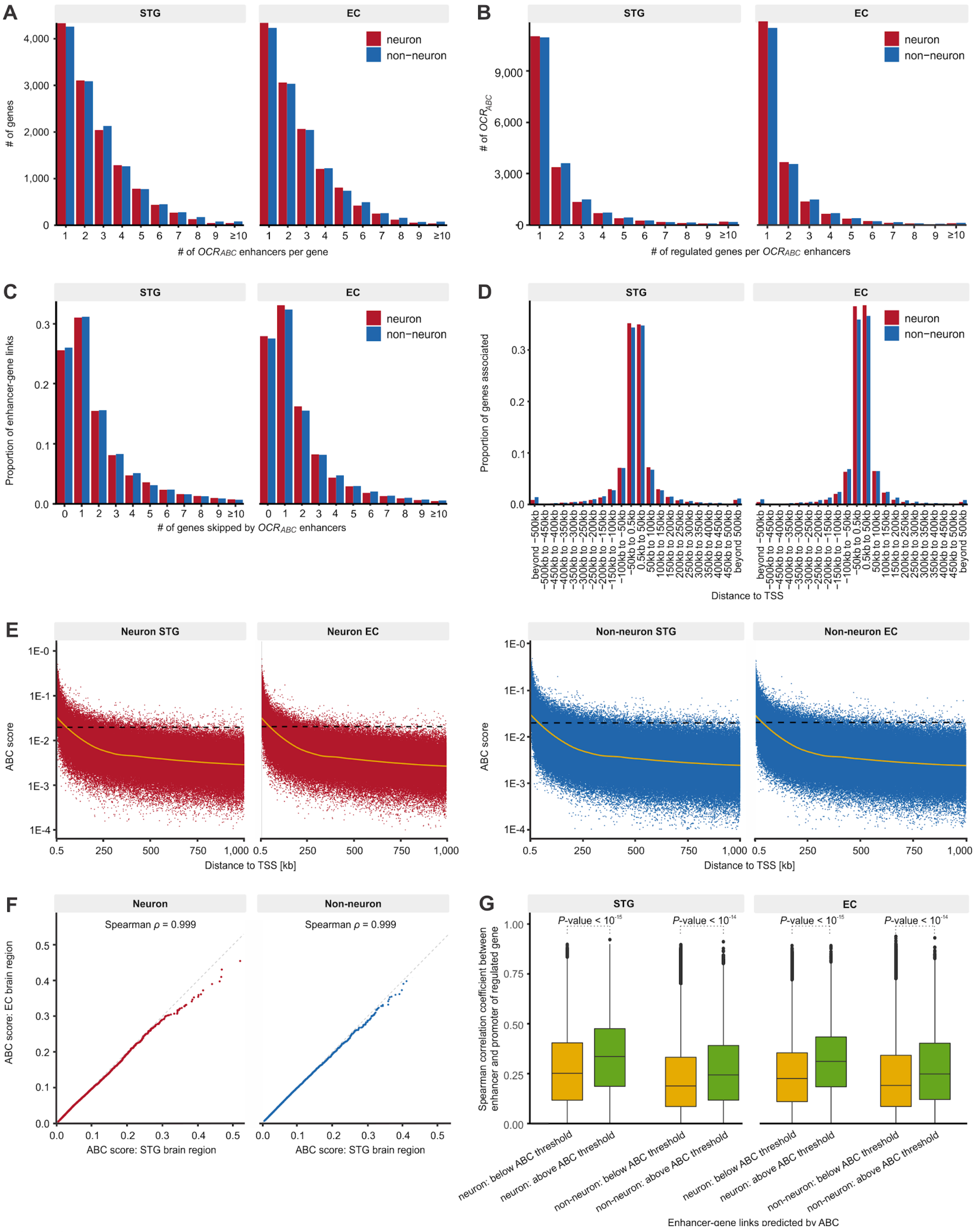


**Fig. S11.**
Linking distal regulatory OCRs ($OCR_{ABC}$) to genes using the Activity-By-Contact (ABC) method. (**A**) Histogram of the number of $OCR_{ABC}$ linked per gene. (**B**) Histogram of the number

of genes linked per $OCR_{ABC}$. (**C**) Histogram of the number of genes "skipped" by an $OCR_{ABC}$ to reach their linked genes. (**D**) Histogram of the distance of $OCR_{ABC}$ to the TSS of regulated genes. (**E**) Scatterplot of genomic distance vs ABC score of enhancer-gene links. The dashed black line denotes the minimum ABC score for an enhancer-gene link to be reported as a valid association. The solid yellow line is LOESS (local regression) fit. (**F**) Correlations between ABC score calculated for enhancer-gene links in STG and EC brain region. The correlation is calculated for all enhancer-gene links that exceed minimum ABC threshold in at least one brain region. (**G**) Correlation between enhancer OCRs and promoter OCRs of linked genes from the ABC method, for links that did not meet the cut-off (low scores, $OCR_{other}$) versus those that did (high scores, $OCR_{ABC}$). *P*-values were calculated by t-test from correlation values converted to Z-scores. The center line indicates the median, the box shows the interquartile range, whiskers indicate the highest/lowest values within 1.5x of the interquartile range, and potential outliers from this are shown as dots.

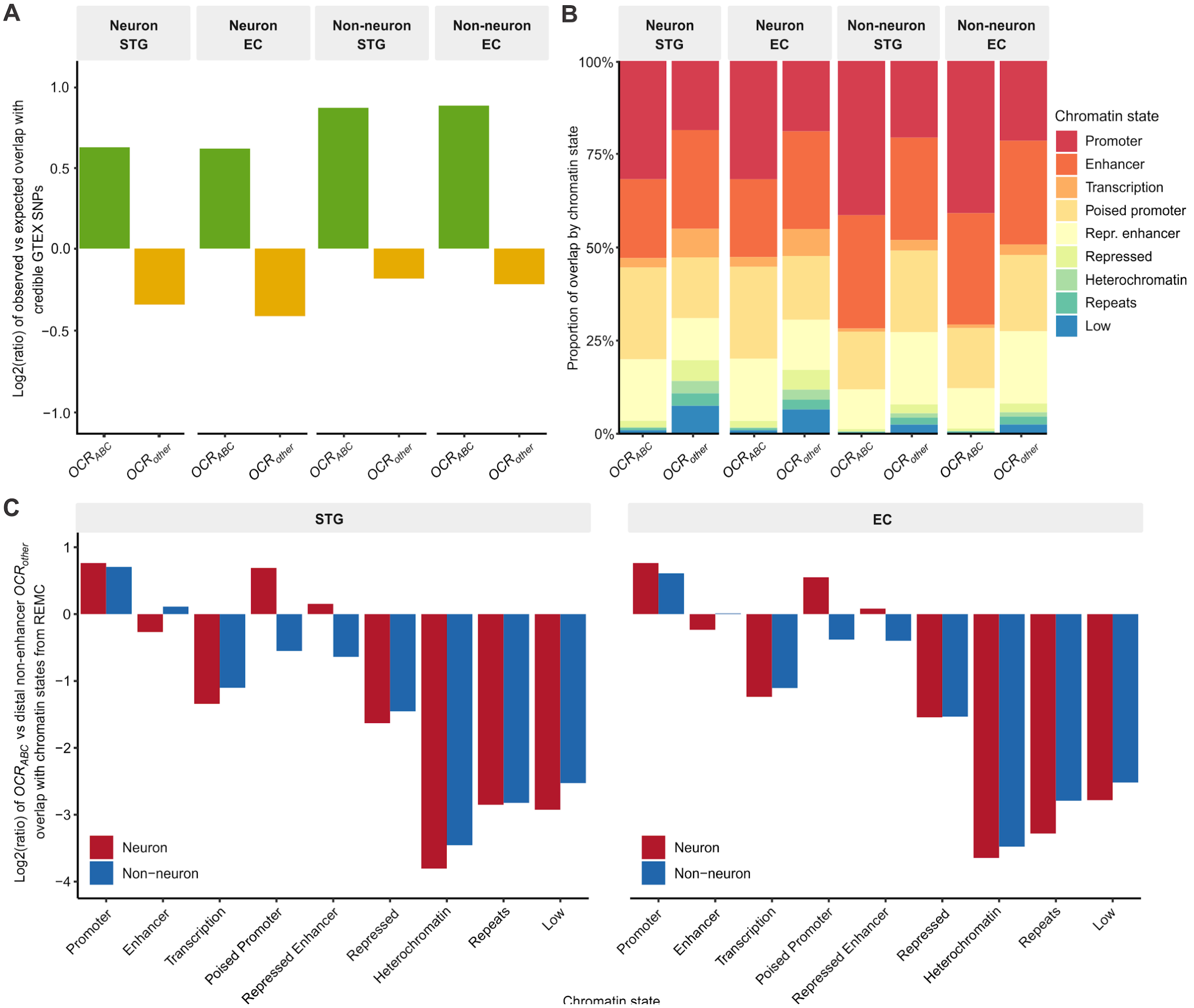


**Fig. S12.**
Comparison of distal OCRs that were predicted to participate in E-P interactions ($OCR_{ABC}$) with distal OCRs that did not regulate genes ($OCR_{other}$). (**A**) Log2 ratio of relative frequency (observed/expected) of overlap of GTEX credible SNPs with open chromatin (Data S5, Materials and Methods). (**B**) Relative frequency (observed/expected overlap in base pairs) of overlaps for each chromatin state with $OCR_{ABC}$ and $OCR_{other}$. Chromatin states were obtained from brain regions included in the Roadmap Epigenomics Project (Materials and Methods). (**C**) Log2 ratio of relative frequency of overlap with chromatin states for distal OCRs meeting the ABC cutoff ($OCR_{ABC}$) versus not meeting the ABC cutoff ($OCR_{other}$).

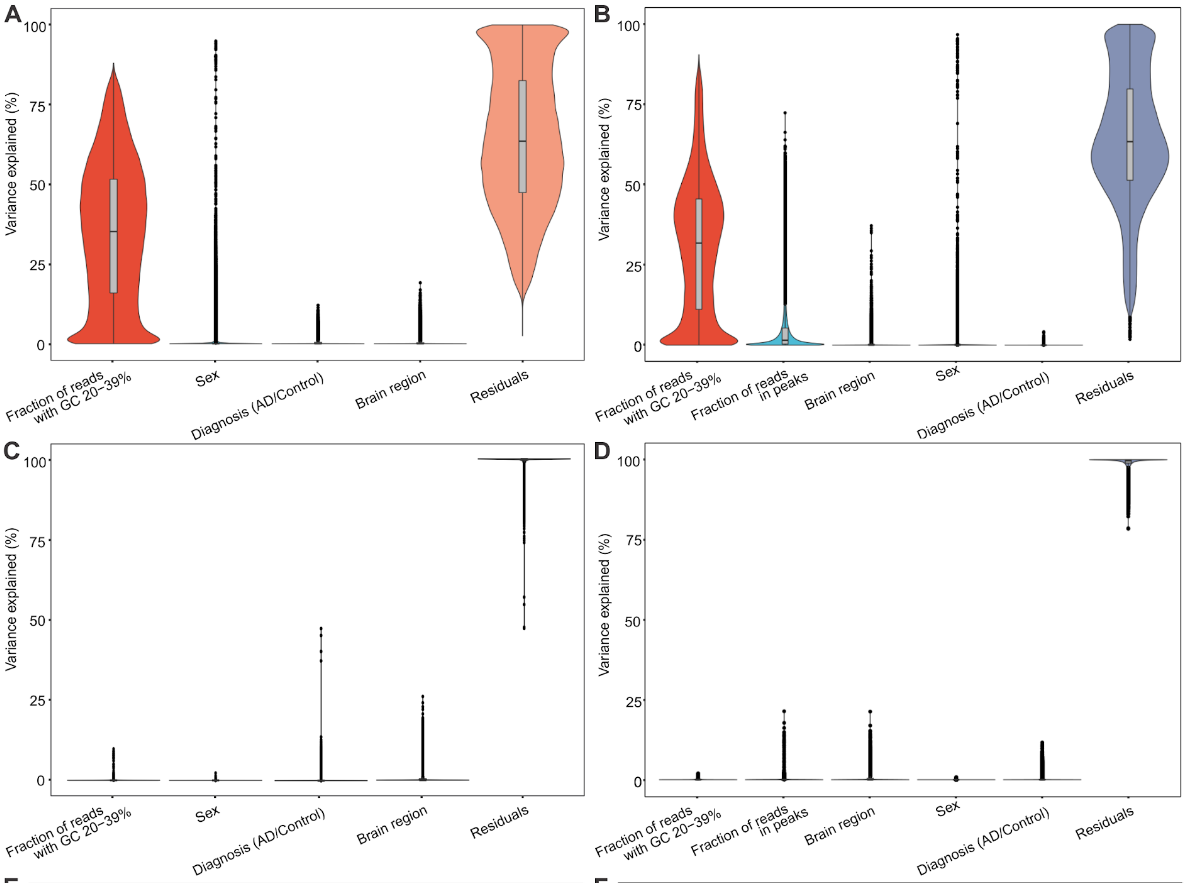


E

| Variable | Variance explained [%] Pre-covariate adjustment | Variance explained [%] Post-covariate adjustment |
|---|---|---|
| Brain region | 0.36 | 0.45 |
| Diagnosis (AD/Control) | 0.37 | 0.06 |
| Sex | 0.47 | 0.00 |
| Fraction of reads with GC 20-39% | 34.68 | 0.04 |
| Residuals | 64.11 | 99.45 |

F

| Variable | Variance explained [%] Pre-covariate adjustment | Variance explained [%] Post-covariate adjustment |
|---|---|---|
| Brain region | 0.29 | 0.44 |
| Diagnosis (AD/Control) | 0.00 | 0.27 |
| Sex | 0.29 | 0.00 |
| Fraction of reads with GC 20-39% | 30.69 | 0.10 |
| Fraction of reads in peaks | 4.66 | 0.06 |
| Residuals | 64.08 | 99.14 |

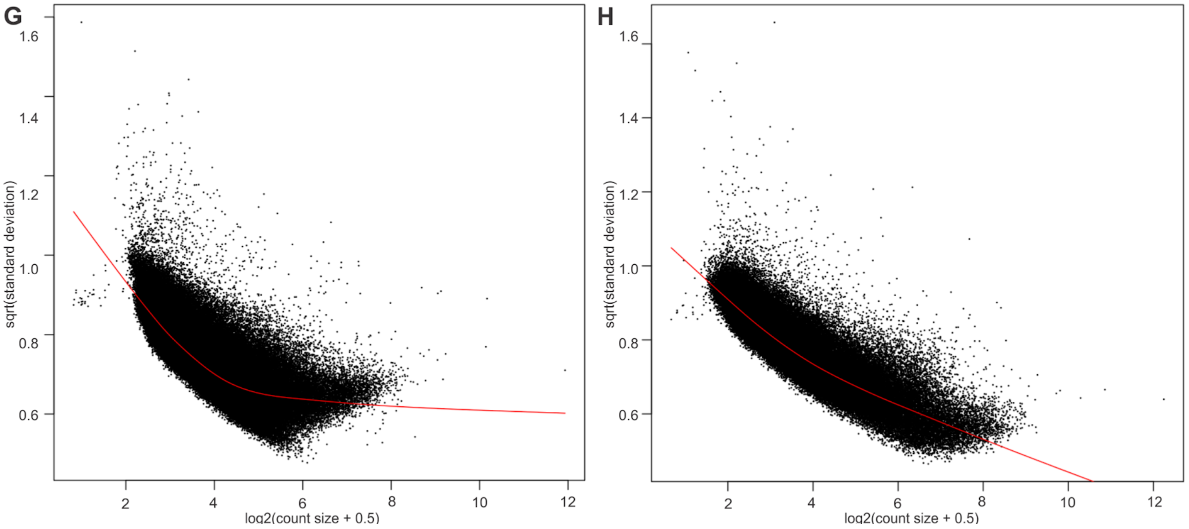


**Fig. S13.**
Variance explained by covariates for (**A**) neurons before adjustment for covariates, (**B**) non-neurons before adjustment for covariates, (**C**) neurons after adjustment for covariates, (**D**) non-neurons after adjustment for covariates. (**E**) Statistics for the variance explained in neurons, (**F**)

statistics for variance explained in non-neurons. (**G**) Variance as a function of read counts in a given OCR for neurons. (**H**) Variance as a function of read counts in a given OCR for non-neurons. In the last two plots, OCRs are represented by black points with LOWESS trends shown in red.

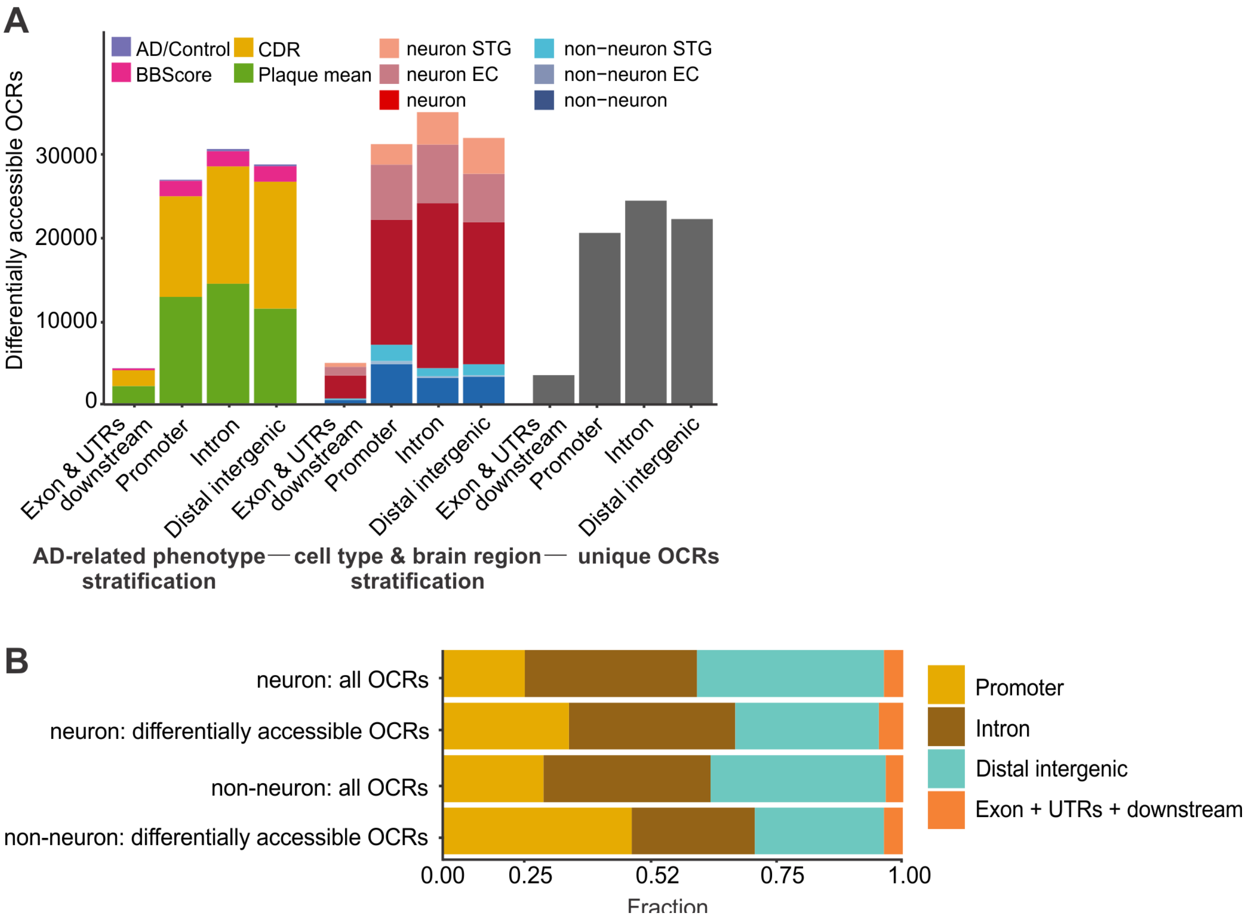


**Fig. S14.**
Analysis of disease-associated OCRs. (**A**) Numbers of differentially accessible OCRs stratified by genic annotations. (**B**) Comparison of relative proportions of genic annotations within a set of differentially accessible OCRs and all OCRs.

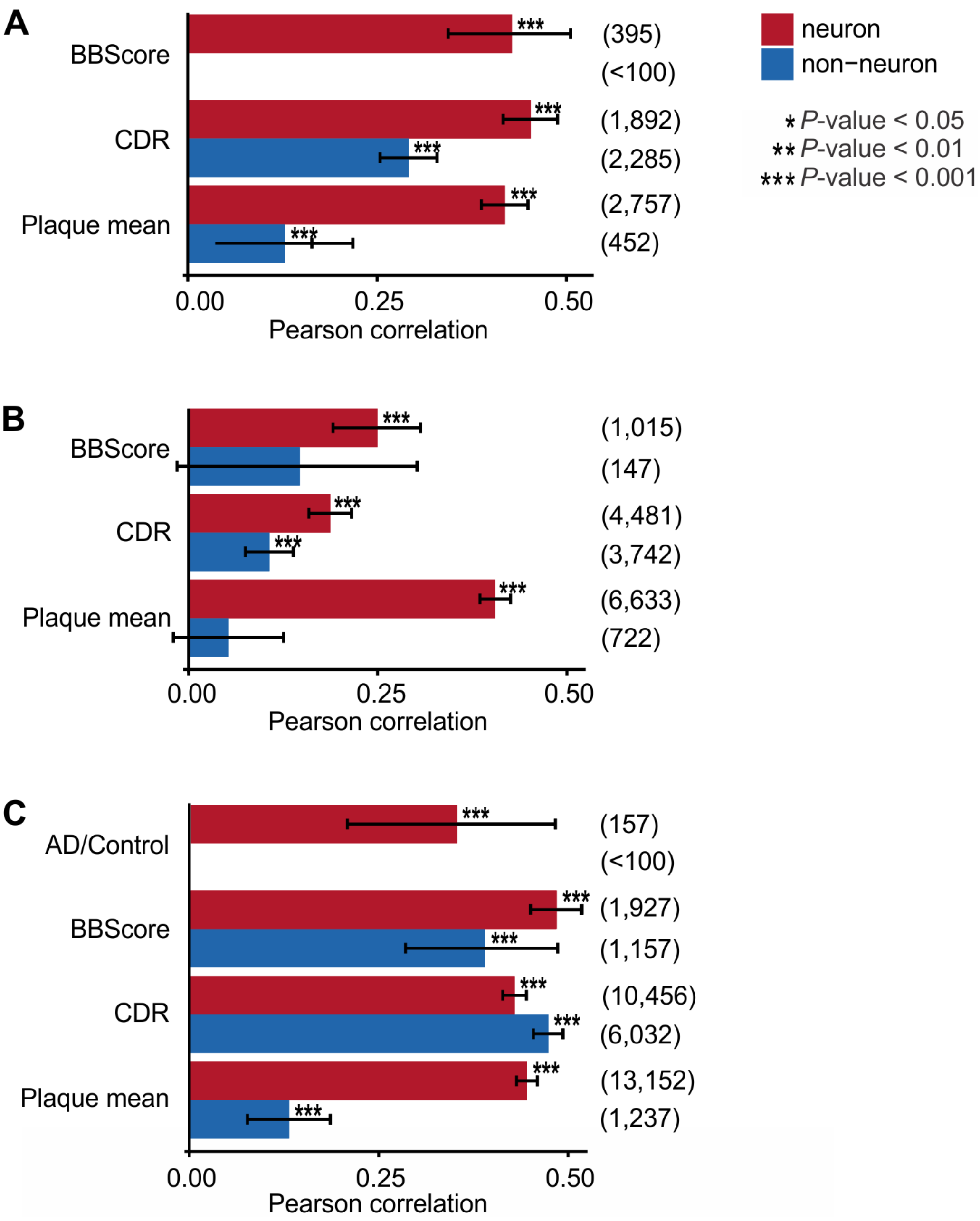


**Fig. S15.**
Concordance between sets of differentially accessible OCRs from this study and other epigenetic studies of AD. The Pearson correlation coefficient was calculated based on log(fold change) of accessibility in our ATAC-seq OCRs and log(fold change) (alternatively correlation to Tau protein load for (A)) of overlapping OCRs from an external dataset. To consider OCRs as overlapping, there need to be at least 25% overlap of OCRs from an external dataset by OCRs from our dataset. The numbers on the right side of the plot show the number of overlapping OCRs used for calculation of correlation coefficient. (**A**) Comparison to the Pearson correlation coefficients of H3K9ac ChIP-seq peaks and Tau protein load (*3*). (**B**) Comparison to the log(fold change) of ATAC-seq OCRs in iPSC-derived forebrain neurons overexpressing MAPT gene that

encodes the Tau protein (*3*). (**C**) Comparison to log(fold change) of ATAC-seq OCRs in iPSC-derived neurons from 3 AD cases and 3 controls (*28*).

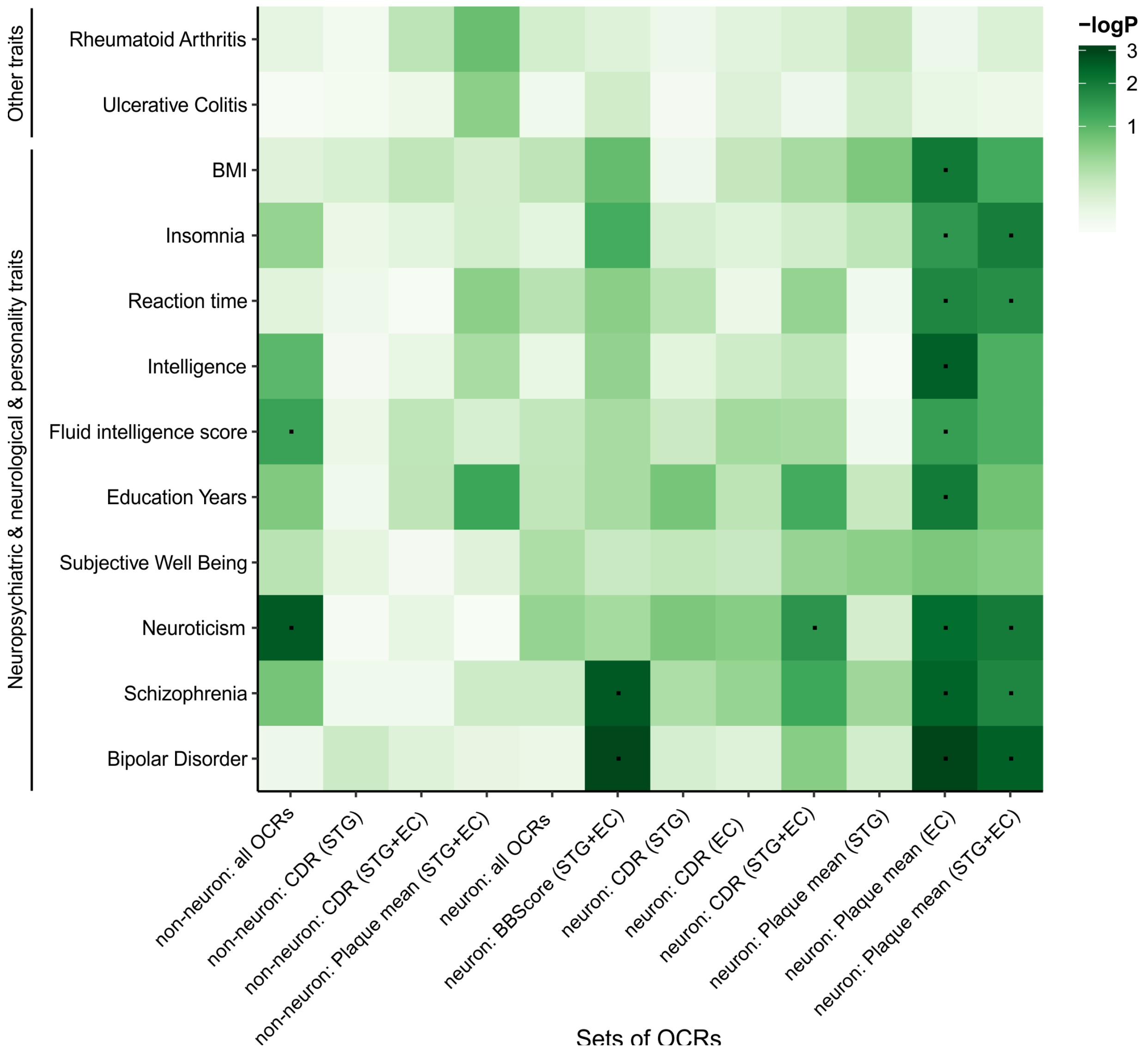


**Fig. S16.**

Overlap between genetic variants associated with brain and non-brain related traits and AD associated OCRs from this study. The analysis was performed using LD-score partitioned heritability. Ulcerative colitis and rheumatoid arthritis were meant as negative controls, since they are not thought to be brain-related traits. None of the tested traits and OCRs overlaps showed an FDR-significant enrichment (correcting for all 144 combinations). " · ": nominally significant at $P < 0.05$.

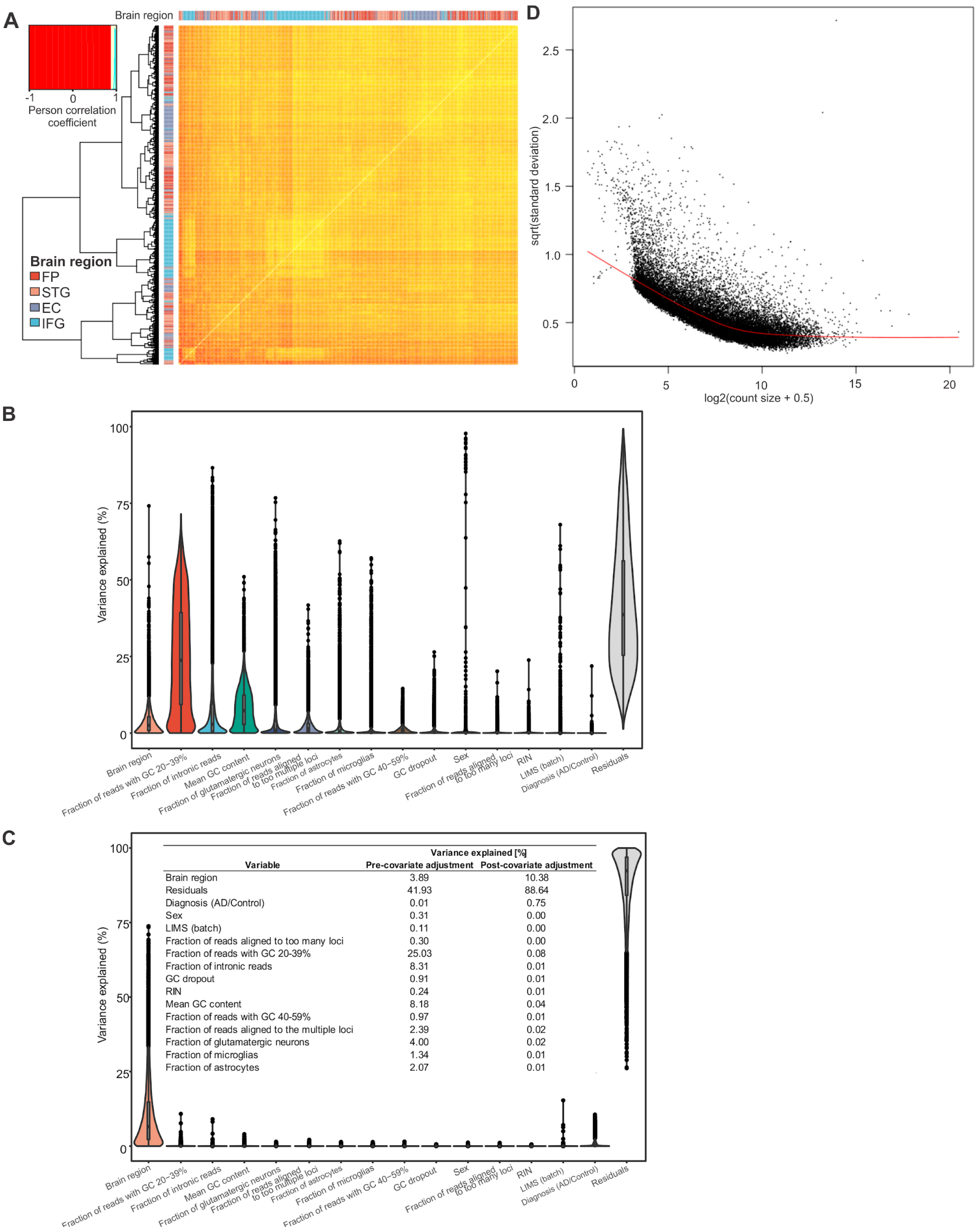


**Fig. S17.**
Analysis of RNA-seq dataset. (**A**) Pearson correlation heatmaps of expression of RNA-seq genes between all samples. (**B**) Distribution of variance explained by covariates before adjusting for

covariates. (**C**) Distribution of variance explained by covariates after adjusting for covariates. Covariates were selected by repeated BIC procedure (Materials and Methods). (**D**) Variance as a function of read counts in a given gene. OCRs are represented by black points with LOWESS trends shown in red that indicate low - moderate biological variation.

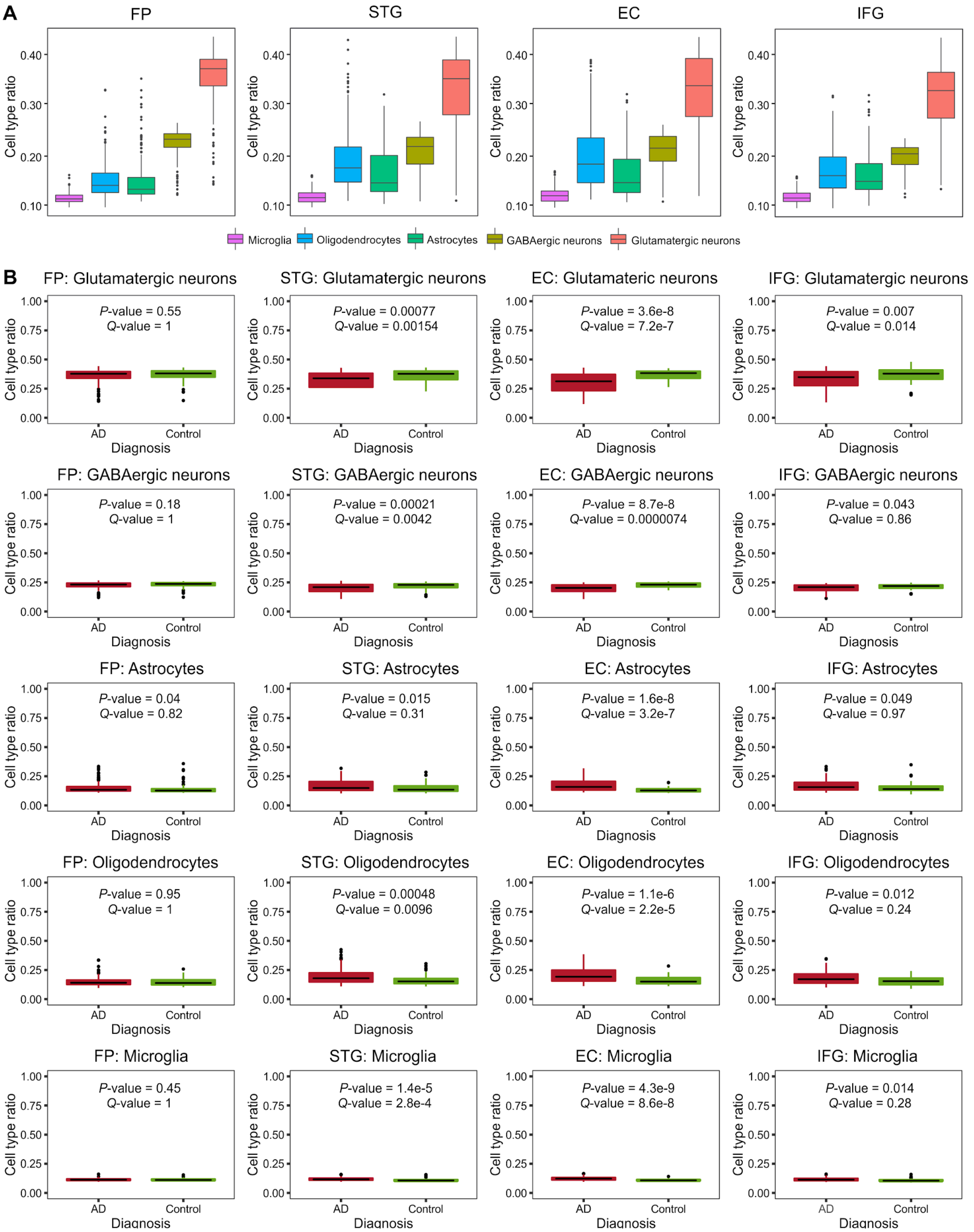


**Fig. S18.**
Cell type deconvolution of RNA-seq datasets. (**A**) Distribution of predicted cell type composition for each brain region. (**B**) Distribution of predicted cell type composition stratified

by AD case/control diagnosis status. The differences were tested by a Wilcoxon test and adjusted for multiple testing. The center line indicates the median, the box shows the interquartile range, whiskers indicate the highest/lowest values within 1.5x of the interquartile range, and potential outliers from this are shown as dots.

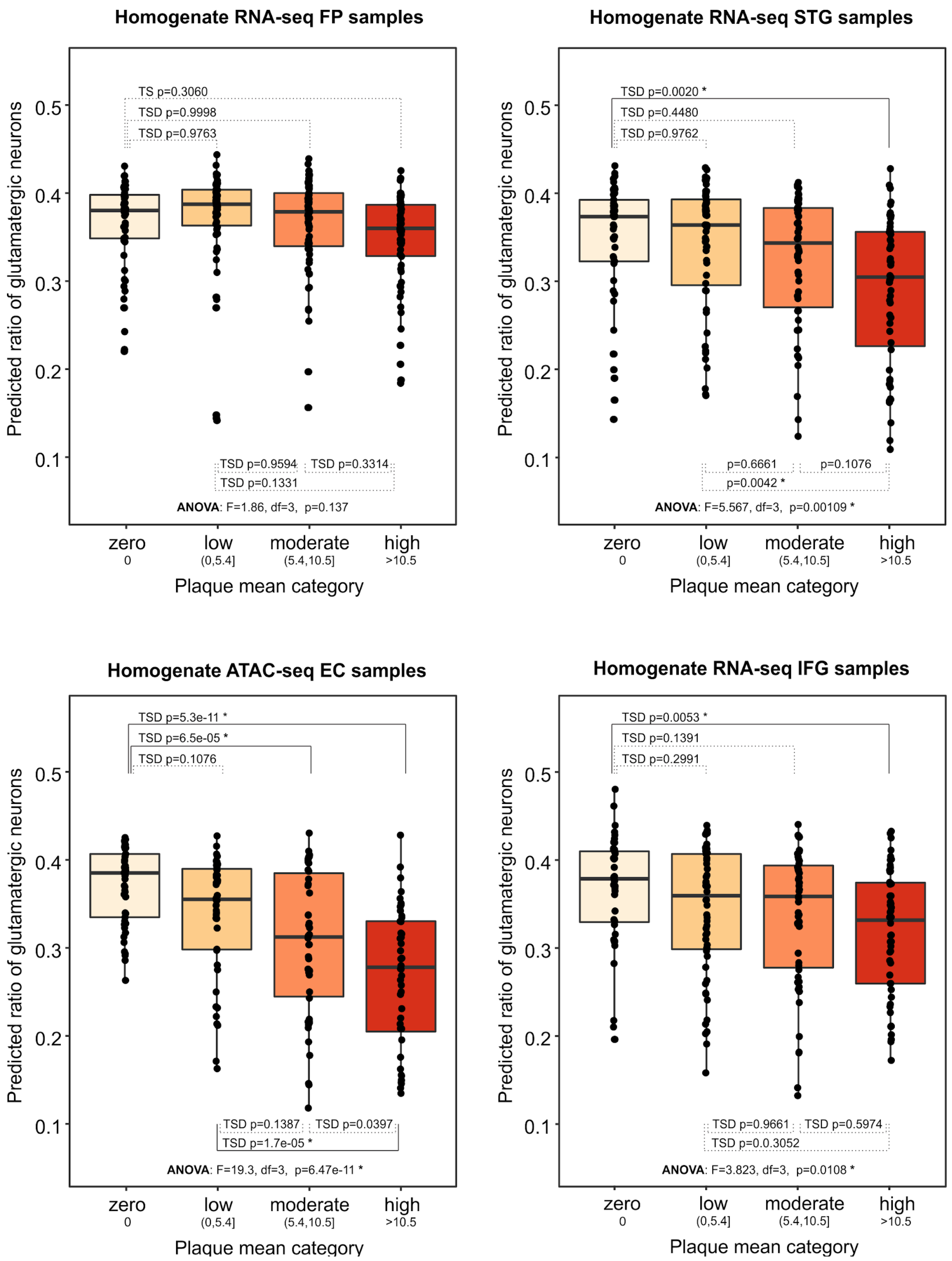


**Fig. S19.**
The relative loss of glutamatergic neurons as a consequence of disease progression is the most prominent for EC region bulk RNA-seq samples, followed by STG, IFG, and FP. We binned RNA-seq samples in 4 categories according to their plaque mean values. Then, we plotted the predicted ratios of glutamatergic neurons per category and ran ANOVA followed by Tukey's

honest significance test (TSD) to determine the significance of differences of glutamatergic ratios between plaque mean categories. The center line indicates the median, the box shows the interquartile range, whiskers indicate the highest/lowest values within 1.5x of the interquartile range. Dots indicate the individual data points.

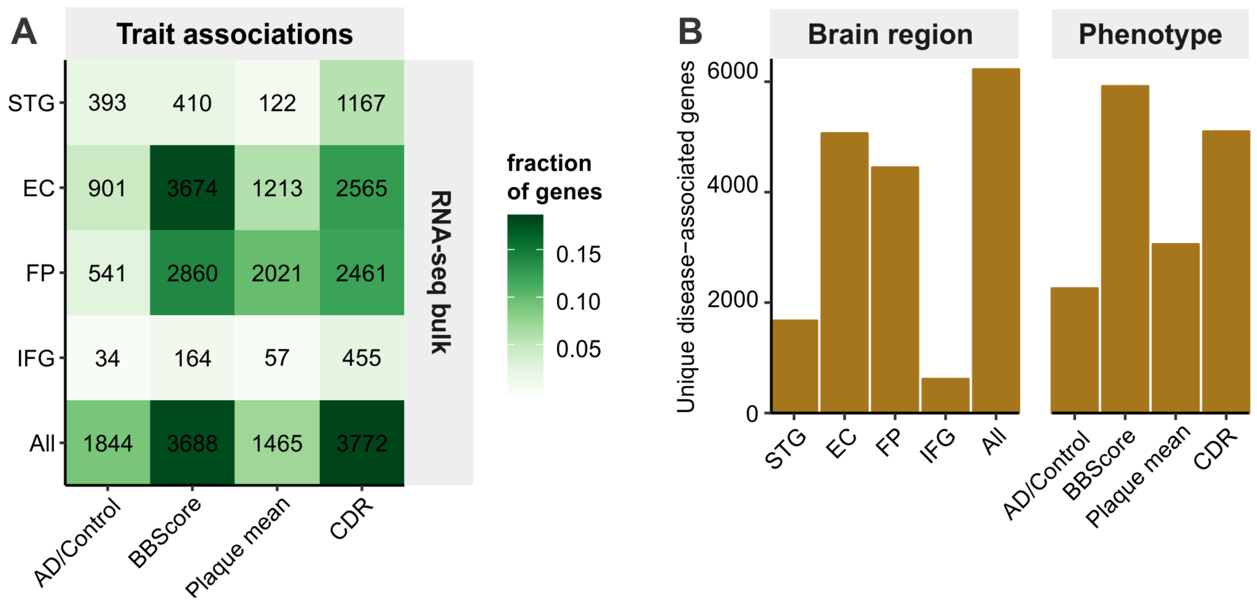


**Fig. S20.**
Number of genes showing significant expressional changes in AD and related phenotypes. (**A**) Heatmap of significant associations by brain region and phenotype. (**B**) Aggregated numbers of unique genes showing associations in a certain brain region or phenotype.

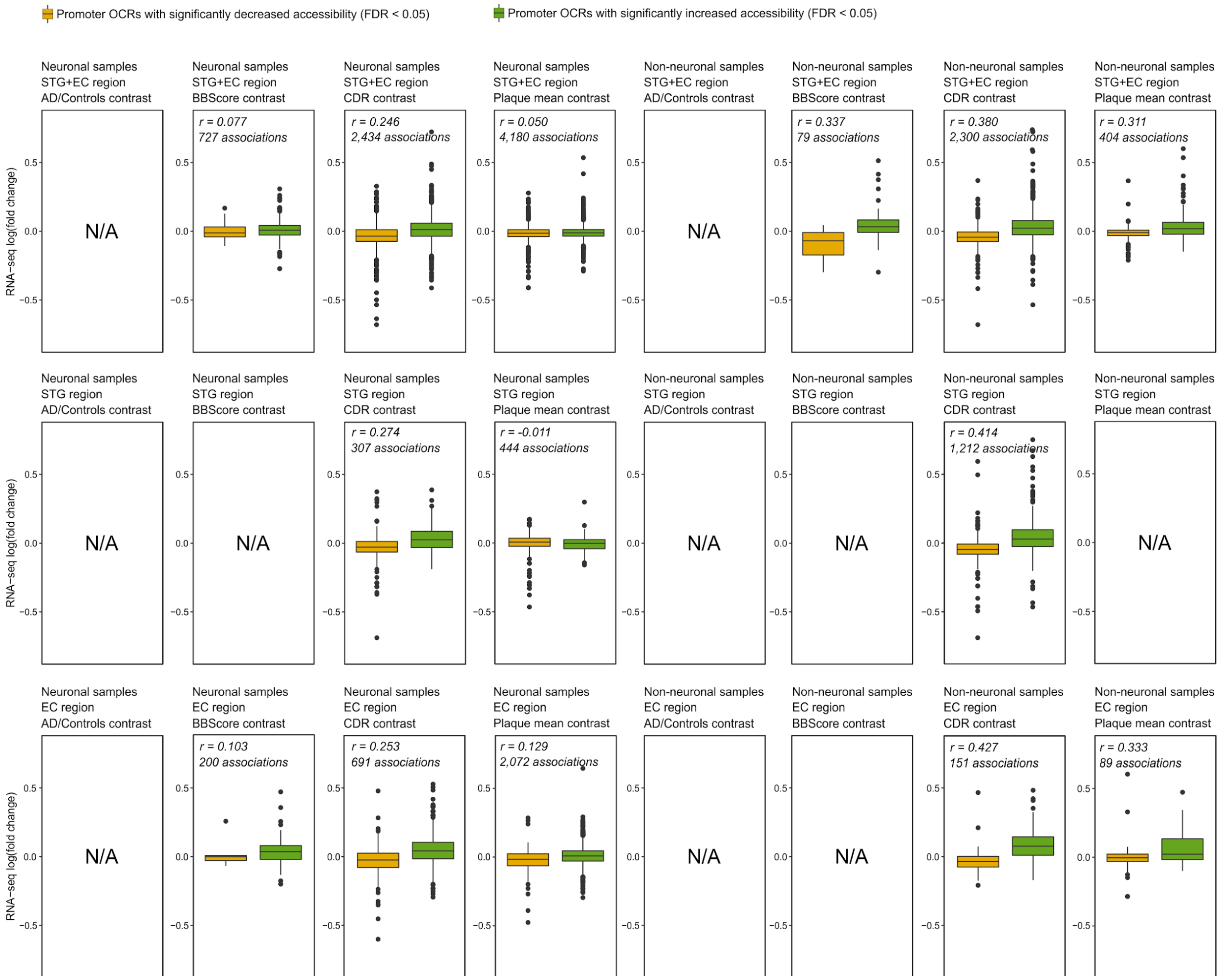


**Fig. S21.**
Correlation between AD associated changes in transcription (RNA-seq) and AD associated changes in the epigenome (ATAC-seq). The boxplots show the change of transcription of genes with significant increase/decrease (FDR<0.05) of accessibility of their respective promoter OCRs (as defined by multiple AD-related phenotypes). Pearson correlation of ATAC-seq and RNA-seq log(fold change) is depicted together with the number of associations (only scenarios with more than 50 associations are shown; only protein-coding genes are considered). The center line indicates the median, the box shows the interquartile range, whiskers indicate the highest/lowest values within 1.5x of the interquartile range, and potential outliers from this are shown as dots.

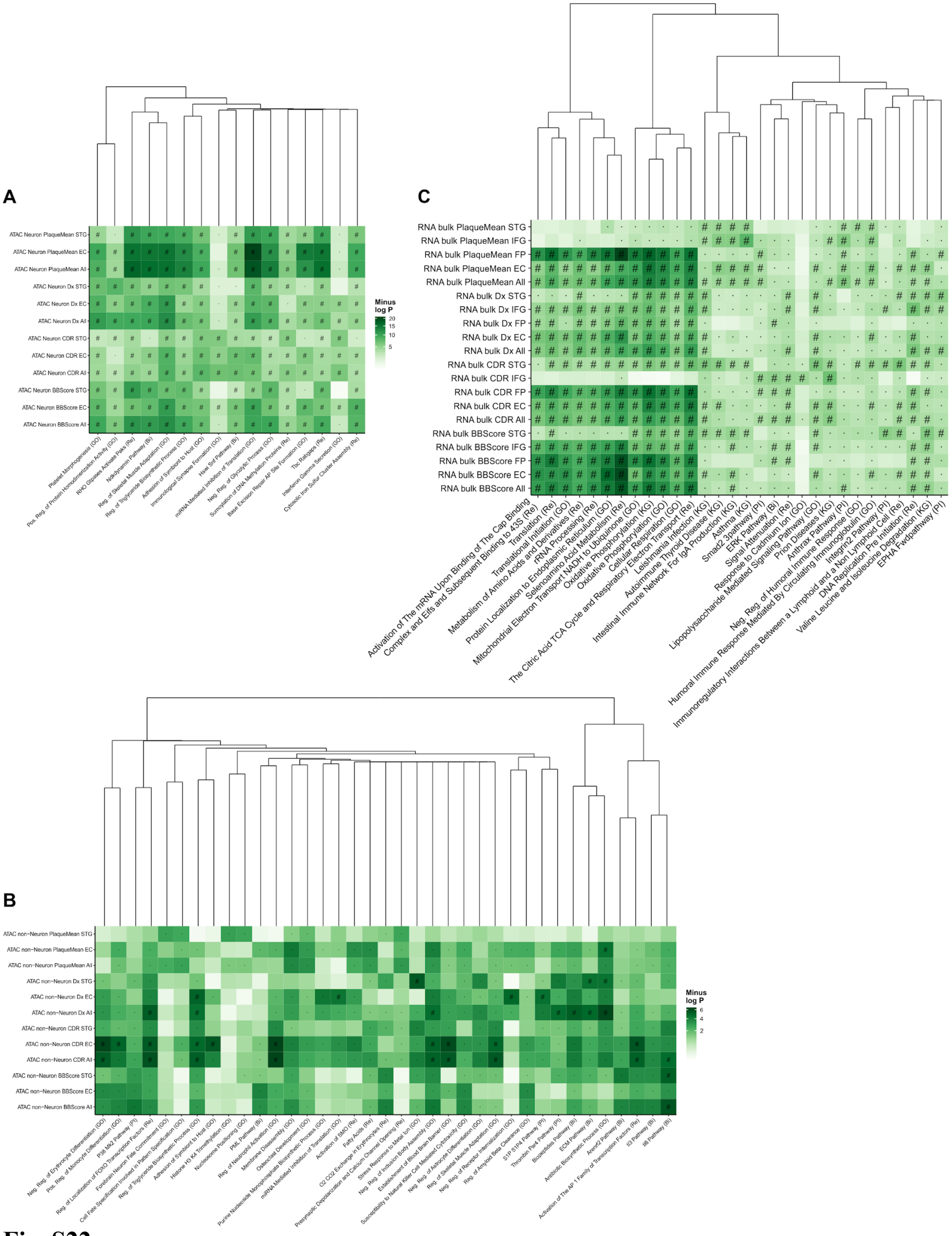


**Fig. S22.**
Gene set enrichment analysis using general gene sets. Top five gene sets of each contrast tested in each assay. The gene sets are clustered based on the constituent member genes. FDR throughout is calculated for all tests within one assay (e.g. all contrasts analyzed in ATAC Neuron times the number of gene sets tested). (**A**) ATAC-seq neuron (**B**) ATAC-seq non-neuron

(**C**) RNA-seq bulk. “#” Significance at FDR<5%. “ · ”: Nominal significance. “Bi”: Biocarta. “GO”: Gene Ontology. “KG”: Kegg. “PI”: protein interaction database. “Re”: Reactome. The number of gene sets varies by assay as the same gene set would often be among the top gene sets across multiple contrasts.

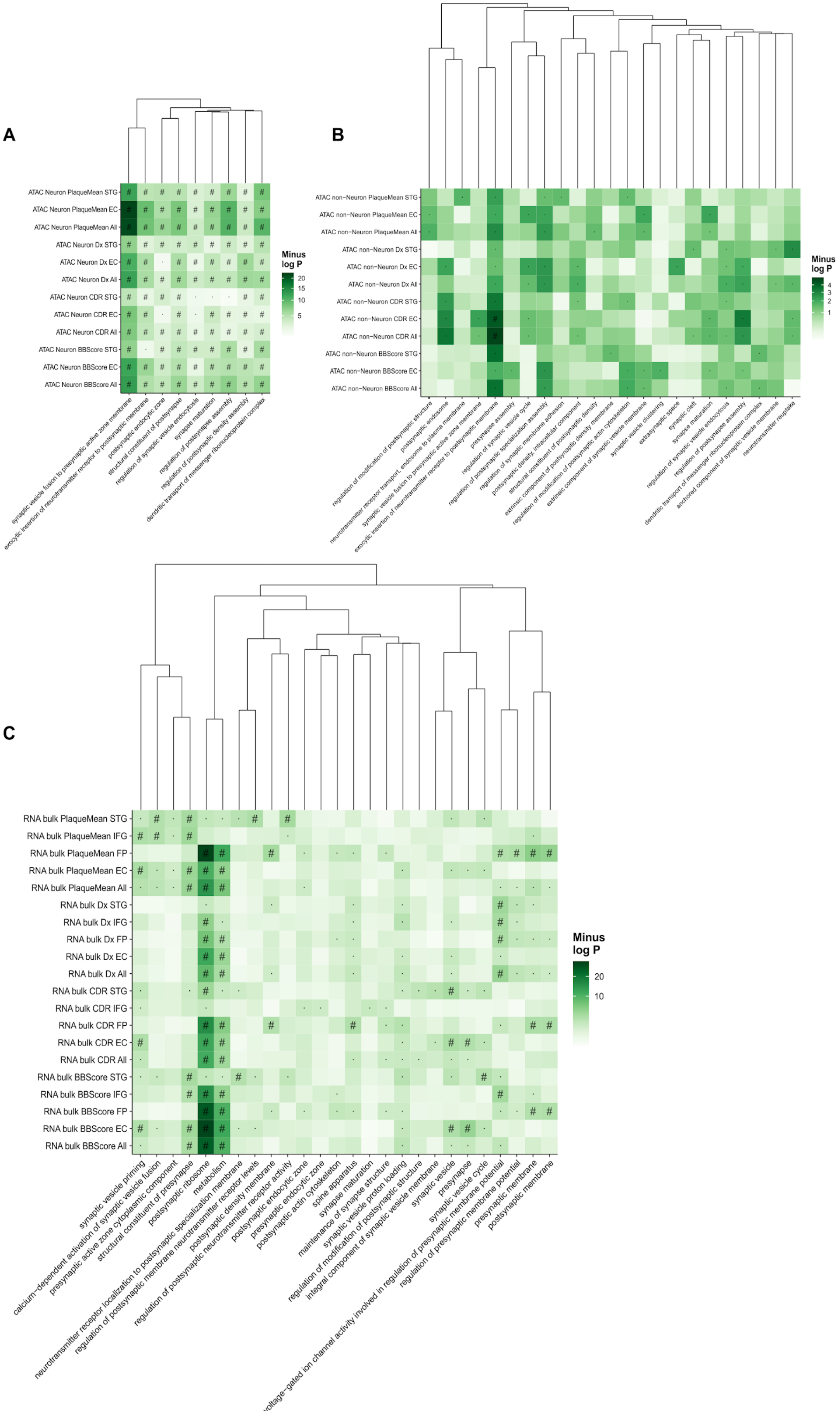
A
ATAC Neuron PlaqueMean STG
ATAC Neuron PlaqueMean EC
ATAC Neuron PlaqueMean All
ATAC Neuron Dx STG
ATAC Neuron Dx EC
ATAC Neuron Dx All
ATAC Neuron CDR STG
ATAC Neuron CDR EC
ATAC Neuron CDR All
ATAC Neuron BBScore STG
ATAC Neuron BBScore EC
ATAC Neuron BBScore All
Minus log P
20
15
10
5
synaptic vesicle fusion to presynaptic active zone membrane
exocytic insertion of neurotransmitter receptor to postsynaptic membrane
postsynaptic endocytic zone
structural constituent of postsynapse
regulation of synaptic vesicle endocytosis
synapse maturation
regulation of postsynapse assembly
regulation of postsynaptic density assembly
dendritic transport of messenger ribonucleoprotein complex
B
ATAC non−Neuron PlaqueMean STG
ATAC non−Neuron PlaqueMean EC
ATAC non−Neuron PlaqueMean All
ATAC non−Neuron Dx STG
ATAC non−Neuron Dx EC
ATAC non−Neuron Dx All
ATAC non−Neuron CDR STG
ATAC non−Neuron CDR EC
ATAC non−Neuron CDR All
ATAC non−Neuron BBScore STG
ATAC non−Neuron BBScore EC
ATAC non−Neuron BBScore All
Minus log P
4
3
2
1
regulation of modification of postsynaptic structure
postsynaptic endosome
neurotransmitter receptor transport, endosome to plasma membrane
synaptic vesicle fusion to presynaptic active zone membrane
exocytic insertion of neurotransmitter receptor to postsynaptic membrane
presynapse assembly
regulation of synaptic vesicle cycle
regulation of postsynaptic specialization assembly
regulation of synaptic membrane adhesion
postsynaptic density, intracellular component
structural constituent of postsynaptic density
extrinsic component of postsynaptic density membrane
regulation of modification of postsynaptic actin cytoskeleton
extrinsic component of synaptic vesicle membrane
synaptic vesicle clustering
extrasynaptic space
synaptic cleft
synapse maturation
regulation of synaptic vesicle endocytosis
regulation of postsynapse assembly
dendritic transport of messenger ribonucleoprotein complex
anchored component of synaptic vesicle membrane
neurotransmitter reuptake
C
RNA bulk PlaqueMean STG
RNA bulk PlaqueMean IFG
RNA bulk PlaqueMean FP
RNA bulk PlaqueMean EC
RNA bulk PlaqueMean All
RNA bulk Dx STG
RNA bulk Dx IFG
RNA bulk Dx FP
RNA bulk Dx EC
RNA bulk Dx All
RNA bulk CDR STG
RNA bulk CDR IFG
RNA bulk CDR FP
RNA bulk CDR EC
RNA bulk CDR All
RNA bulk BBScore STG
RNA bulk BBScore IFG
RNA bulk BBScore FP
RNA bulk BBScore EC
RNA bulk BBScore All
Minus log P
20
10
synaptic vesicle priming
calcium−dependent activation of synaptic vesicle fusion
presynaptic active zone cytoplasmic component
structural constituent of presynapse
postsynaptic ribosome
metabolism
neurotransmitter receptor localization to postsynaptic specialization membrane
regulation of postsynaptic membrane neurotransmitter receptor levels
postsynaptic density membrane
regulation of postsynaptic neurotransmitter receptor activity
postsynaptic endocytic zone
presynaptic endocytic zone
postsynaptic actin cytoskeleton
spine apparatus
synapse maturation
maintenance of synapse structure
synaptic vesicle proton loading
regulation of modification of postsynaptic structure
integral component of synaptic vesicle membrane
voltage−gated ion channel activity involved in regulation of presynaptic membrane potential
synaptic vesicle
presynapse
synaptic vesicle cycle
regulation of presynaptic membrane potential
presynaptic membrane
postsynaptic membrane

**Fig. S23.**
Gene set enrichment analysis using Syngo gene sets. Top five gene sets of each contrast tested in each assay. The gene sets are clustered based on the constituent member genes. FDR throughout is calculated for all tests within one assay (e.g. all contrasts analyzed in ATAC Neuron times the number of gene sets tested). (**A**) ATAC-seq neuron (**B**) ATAC-seq non-neuron (**C**) RNA-seq bulk. “#” Significance at FDR<5%. “ · ”: Nominal significance. The number of gene sets varies by assay as the same gene set would often be among the top gene sets across multiple contrasts.

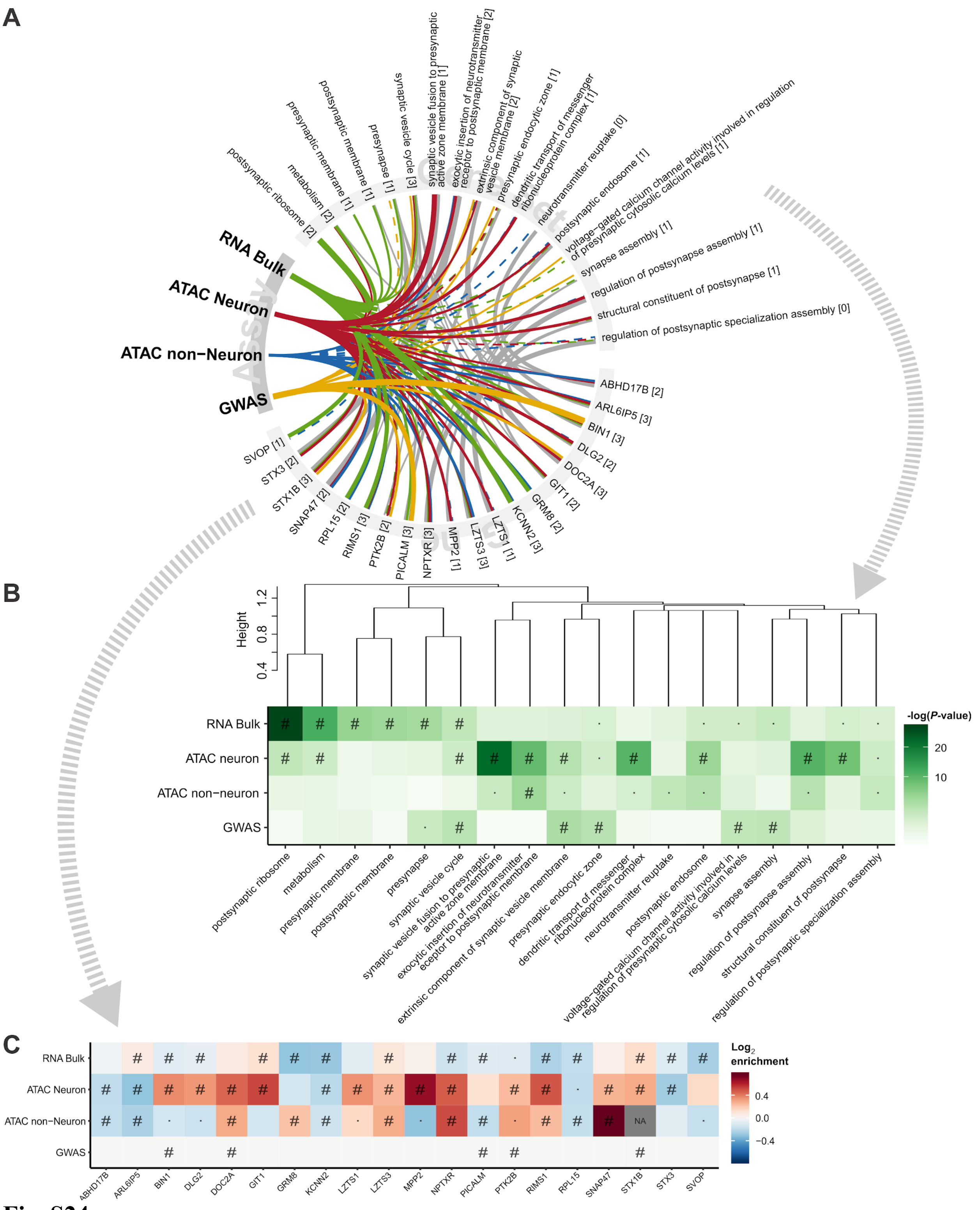


**Fig. S24.**
Gene set enrichment analysis using synaptic gene ontology gene sets (Syngo). (**A**) Top five gene sets of the four overall assays and the top genes within these. Thickness indicates the strength of association. A full line indicates significance at FDR<5%, whereas a dashed line indicates

nominal significance. Grey lines between genes and gene sets indicate gene set membership and the thickness is inversely proportional to the size of the gene set. The numbers in square brackets indicate the number of assays in which the gene or gene set was found significant at FDR<5%. The gene sets are clustered based on the constituent member genes. FDR throughout is calculated for all tests within one assay (e.g. all contrasts analyzed in ATAC Neuron times the number of gene sets tested). (**B**) heatmap of *P*-values for the associated gene sets. (**C**) Heatmap of fold changes in chromatin accessibility/gene expression and significance of change. The GWAS associations are genetic variants which might increase, decrease or otherwise alter gene function, thus, no colors are applied to these. "#" Significance at FDR<5%. " · ": Nominal significance. "NA": not available. The ApoE- and MHC-loci were excluded from the GWAS analysis, as is customary. Not all genes have an OCR directly linked to it in neuron and non-neuron ATAC-seq.

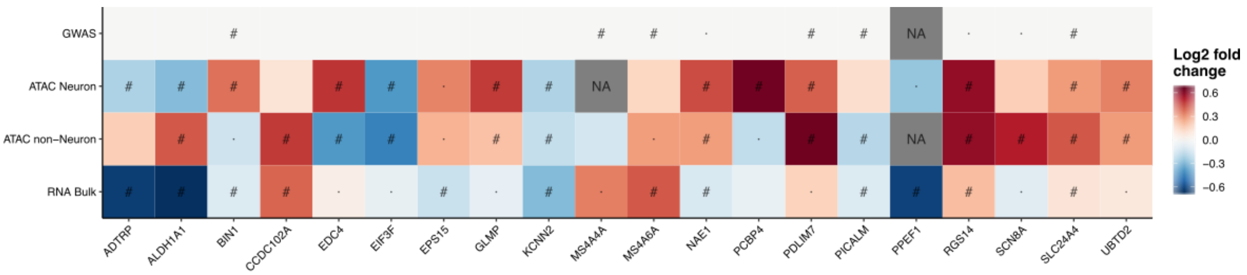


**Fig. S25.**
Heatmap of top five genes across the four overall assays with fold changes in chromatin accessibility/gene expression and significance. FDR throughout is calculated for all tests within one assay (e.g. all contrasts analyzed in ATAC Neuron times the number of gene sets tested). "#" Significance at FDR<5%. " · ": Nominal significance. "NA": not available. The ApoE- and MHC-loci were excluded from the GWAS analysis, as is customary. Not all genes have an OCR directly linked to it in neuron and non-neuron ATAC-seq.

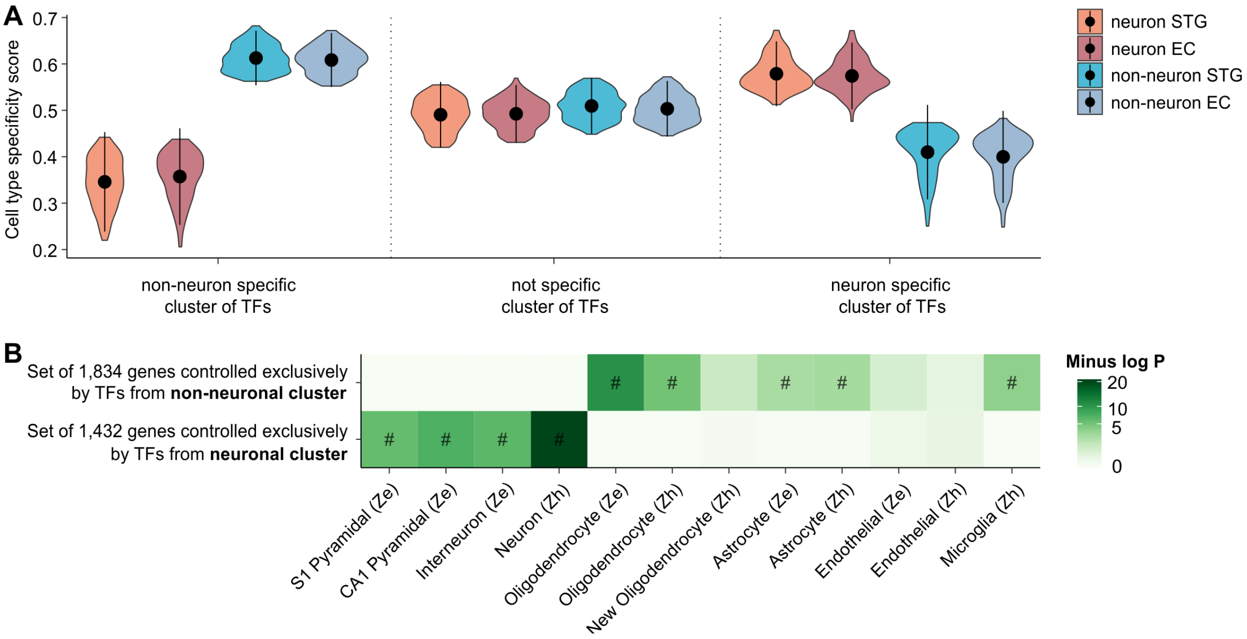


**Fig. S26.**
Cell type specificity of transcription factors and their regulatory targets. (**A**) Distribution of the cell type specificity score across cell types (neuron & non-neuron) and brain regions (STG & EC) for TFs belonging to the same cluster as assigned by hierarchical clustering. Black dots indicate means. Vertical lines denote the mean plus / minus two times the standard deviations. (**B**) Overlap between the sets of genes exclusively regulated by TFs from only neuronal / non-neuronal TF clusters and cell-specific marker genes from (*16,97*). "#": FDR < 0.05 " · ": nominal *P*-value < 0.05. "TF": transcription factor.

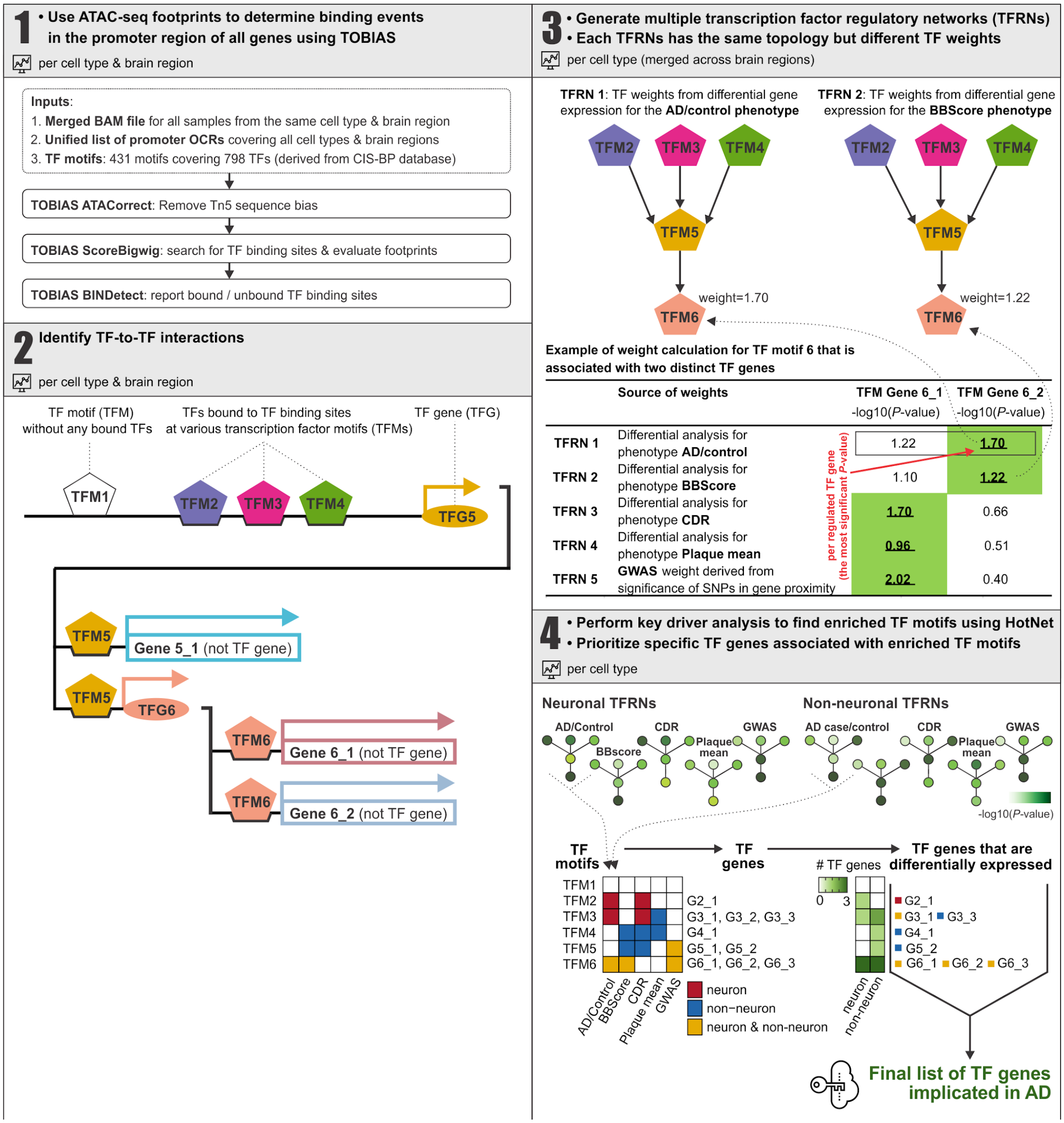


**Fig. S27.**

Workflow to identify transcription factors implicated in epigenetic reprogramming in AD. Panel 2 shows an example of TF to TF interactions, where transcription factor gene 5 (TFG5 is regulated by three transcription factors and, in part, regulates another non transcription factor gene (gene 5_1) as well as a transcription factor gene (TFG6). Some transcription factor motifs represent multiple genes, as the transcription factors recognize similar or identical motifs and are, thus, bioinformatically indistinguishable in the transcription factor footprinting analysis TF: transcription factor; TFM: transcription factor motif; TFG: Transcription factor gene; TFRN transcription factor regulatory network.

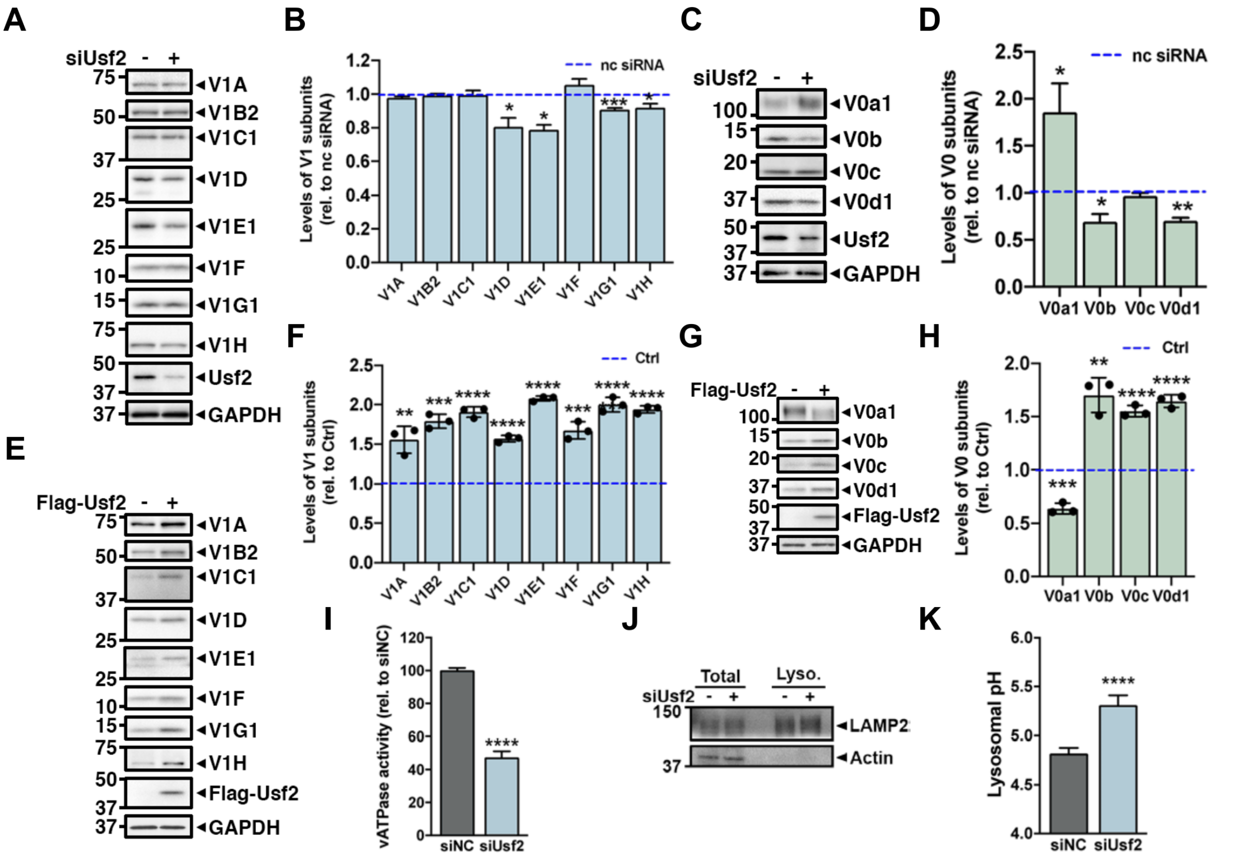


**Fig. S28.**

TF USF2 knockdown and overexpression in the human neuroblastoma cell line, SH-SY5Y. (**A-D**) Immunoblot analysis of v-ATPase V1 (A) and V0 (C) subunits in cell lysates transfected with negative control siRNA (nc siRNA; -) or siUSF2 (+) for 72 hr. (**E-H**) Immunoblot analysis of v-ATPase V1 (E) and V0 (G) subunits in lysates of USF2 overexpressing stable SH-SY5Y neuroblastoma cells. The barplots (B, D, F and H) show the band intensity of each protein. GAPDH served as a loading control. (**I**) Lysosomal v-ATPase activity measured colorimetrically ATP hydrolysis with and without a v-ATPase inhibitor (Concanamycin A; ConA). Activity assay performed on lysosomal fractions pre-treated with inhibitors of P- and F-type ATPases (*o*-vanadate) to minimize nonspecific ATPase activity. (**J**) The immunoblot represents purity of lysosomal enriched fraction. LAMP2 served as a marker for lysosome and Actin served as a loading control. (**K**) Lysosomal pH values were measured ratiometrically using LysoSensor Yellow/Blue (Y/B) dextran. All quantitative data were subjected to two-tailed unpaired Student's *t*-test "*": *P*-value<0.05; "**": *P*-value<0.005; "***": *P*-value<0.0005; "****": *P*-value<0.0001.

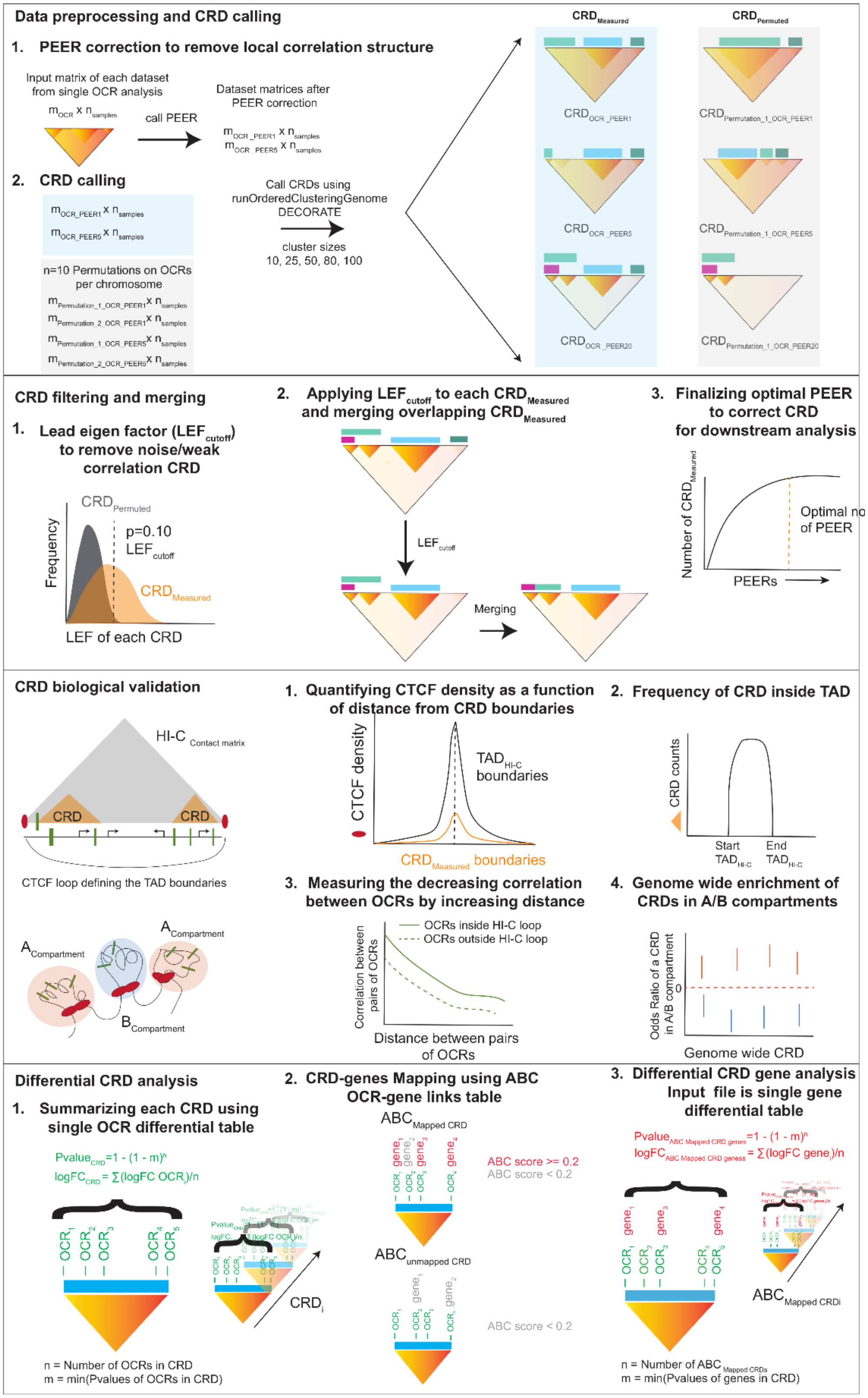


**Fig. S29.** Graphical illustration of the various steps in the CRD analyses.

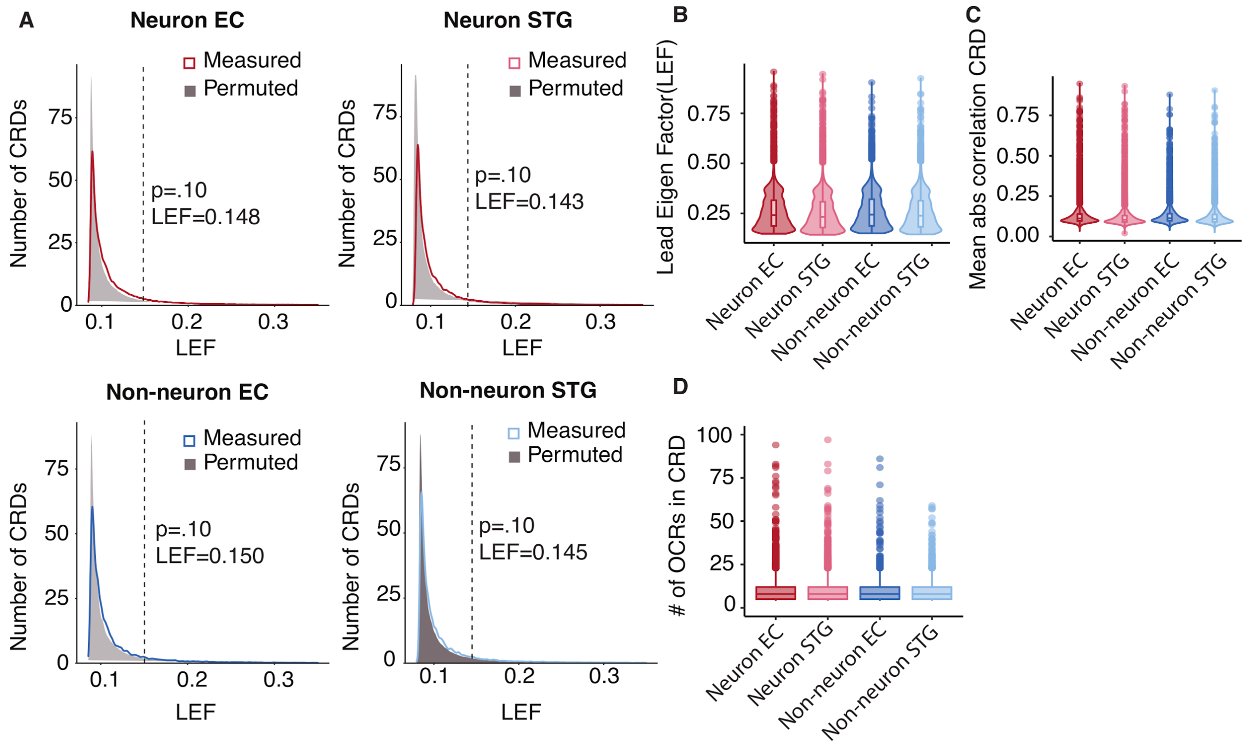


**Fig. S30.**

CRD filtering and merging. (**A**) An example illustrating filtering step using distribution of Lead Eigen Factor (LEF) of CRDs called on $m_{OCR\_PEER} \times n_{samples}$ and $m_{Permutation\_i\_OCR\_PEER} \times n$ samples ($i=10$ permutations were performed). Final $LEF_{cutoff}$ was obtained at $P$-value=0.10 from the distribution of LEF of permuted CRD. (**B-C**) Distribution of (B) Lead eigen factor and (C) mean absolute correlation of CRD called on $m_{OCR\_PEER} \times n_{samples}$ after filtering using $LEF_{cutoff}$ obtained from the previous step (A), stratified by cell type and brain region specific CRDs. (**D**) Boxplot depicting the number of OCRs in CRDs called on $m_{OCR\_PEER25} \times n_{samples}$ of neuronal EC and STG matrices, and $m_{OCR\_PEER15} \times n_{samples}$ non-neuronal EC and STG matrices.

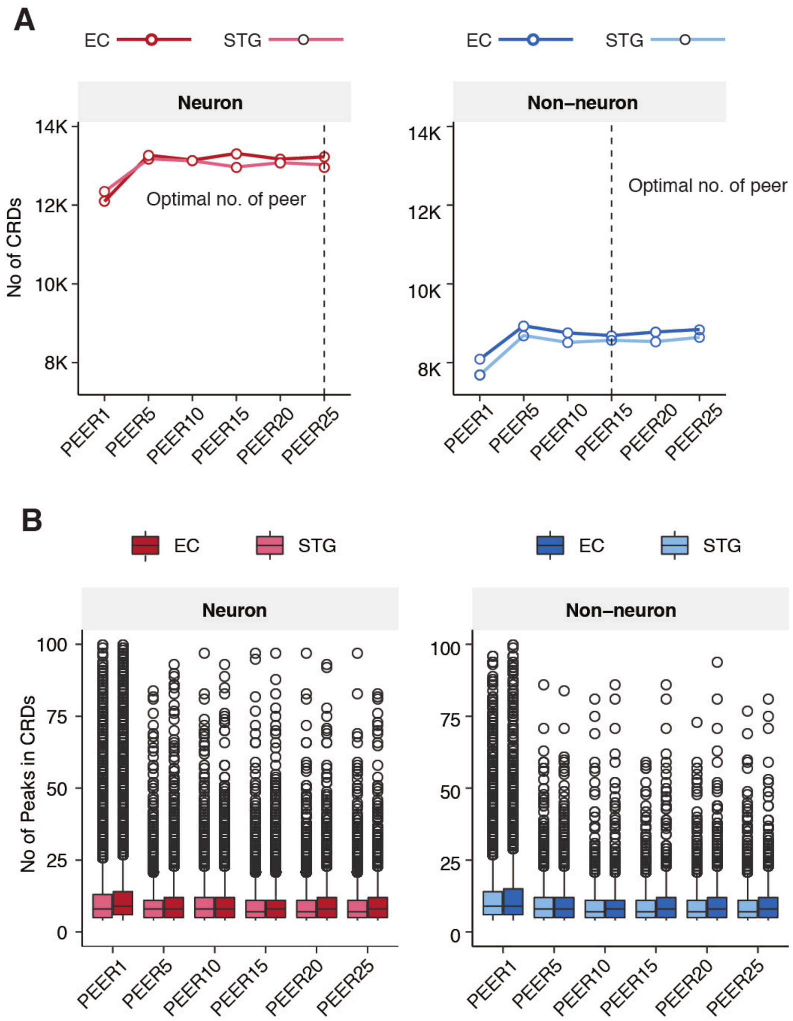


**Fig. S31.**
Selection of optimal number of PEER factors. (**A**) Number of CRDs and (**B**) boxplot depicting the number of OCRs in CRDs obtained after applying $\mathrm{LEF}_{\mathrm{cutoff}}$ and merging the overlapping CRDs called on $m_{OCR\_PEER\_j} \times n_{samples}$, where $j=\{1,5,10,15,20,25\}$.

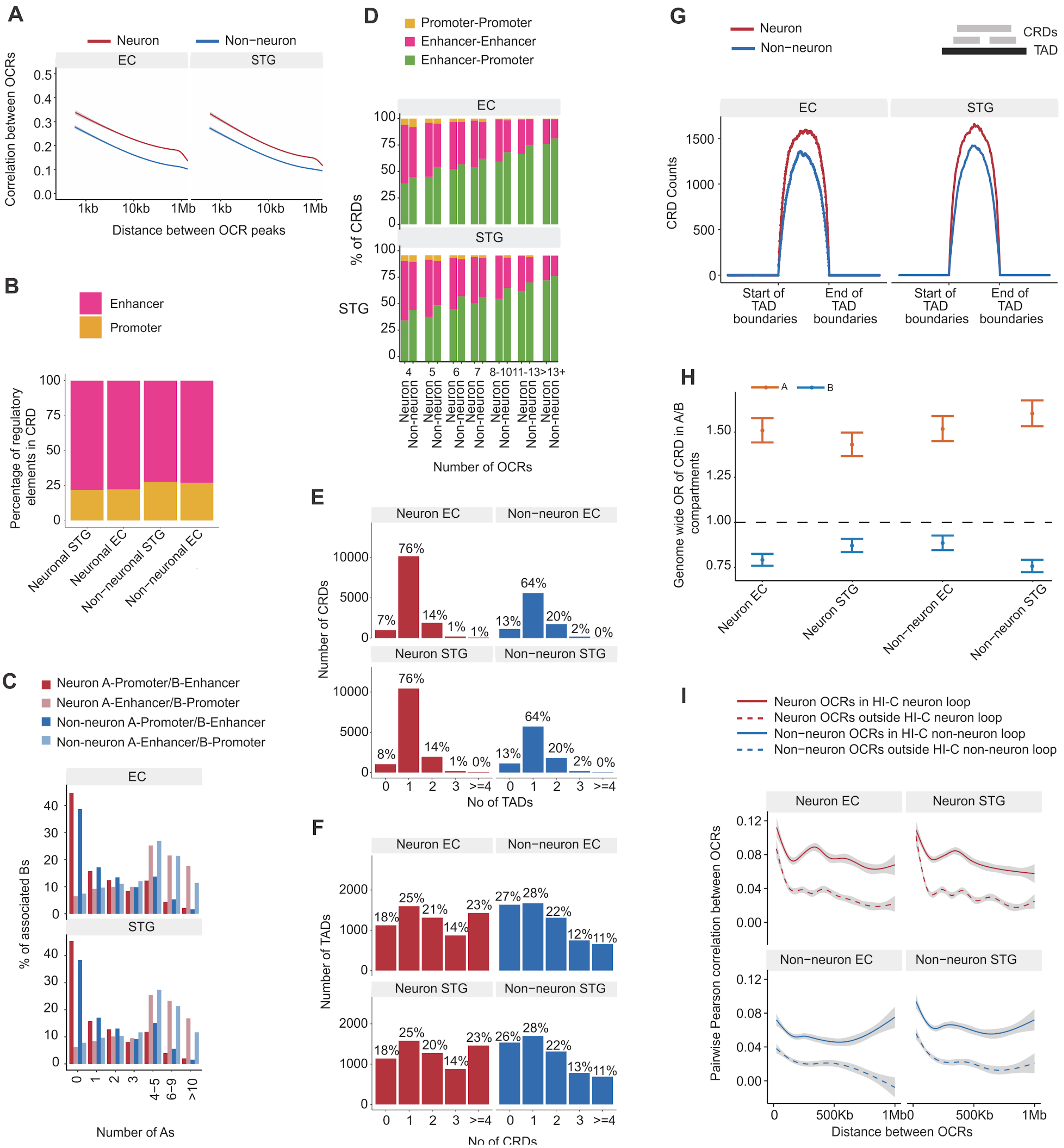


**Fig. S32.**

CRD biological validation. **(A)** An example of pairwise correlation of cell type and brain region specific OCRs in chromosome 22 as a function of distance (within 1Mb). The x-axis is on a log scale. **(B)** Composition of OCRs within CRDs stratified by cell type and brain region. OCRs were considered to be either promoters or enhancers based on the proximity to the nearest TSS. **(C)** Percentage of enhancer OCRs in CRDs stratified by the number of promoter OCRs in CRDs (neuron: dark red, non-neuron: dark blue) and percentage of promoter OCRs in CRDs stratified by the number of enhancer OCRs in CRDs (neuron: light red, non-neuron: light blue). For example, the second dark red bar in the top panel (Number of A's =1) shows the set of EC neuron CRDs with 1 promoter OCR associated with ~15% of enhancers. **(D)** Proportions of cell type and brain region specific CRDs containing only promoter-promoter, enhancer-enhancer, or a mixture of enhancer-promoter as a function of the number of OCRs contained in cell type and region specific CRDs. **(E)** Percentage of CRDs overlapping with TADs stratified by the number of overlapped TADs. (**F**) Fraction of TADs overlapping with CRDs stratified by the number of overlapped CRDs. (**G**) Density of cell type and region specific CRDs that are within one TAD boundary. The distances between TAD boundaries have been normalized between 0 and 1 for each possible interval. The density of CRDs has been computed within 100 bins in each interval. (**H**) Odds ratio of CRDs in A/B compartments with the whole genome as background. Error bars indicate 95% confidence intervals. *P*-values were estimated based on two-sided Fisher's exact. **(I**) Pairwise correlation of cell type and brain region specific OCRs within Hi-C loops as a function of distance shown in solid lines. Pairwise correlations of OCRs outside Hi-C loops are indicated by dashed lines.

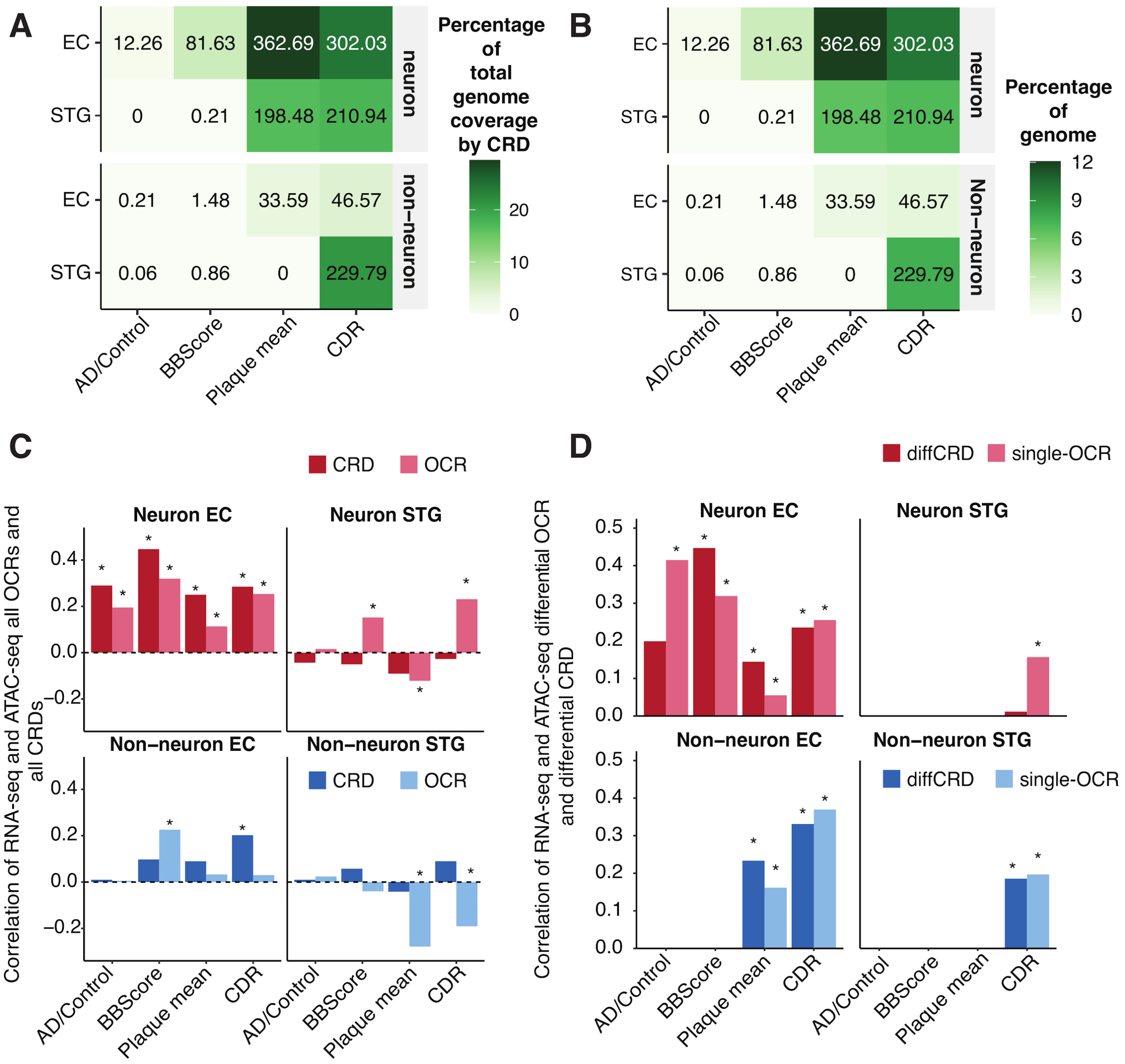


**Fig. S33.**
Differential CRD analysis. **(A)** Percentage of total genomic coverage by CRDs and **(B)** percentage of whole genome coverage by CRDs that are associated with AD-related phenotypes stratified by cell type and brain region. **(C)** Pearson correlation stratified by cell type and brain region between logFC of ABC-mapped genes to CRDs and mean logFC of OCRs in CRDs (dark red/blue), and between logFC of ABC-mapped genes to OCRs and logFC of OCRs (light red/blue). Significant correlations at *P*-value<0.05 are marked as * above the bar. **(D)** Pearson correlation stratified by cell type and brain region between logFC of ABC-mapped genes to CRDs and mean logFC of OCRs in CRDs that were differentially dysregulated at FDR 5% shown (dark red/blue), and between logFC of ABC-mapped genes to OCRs and logFC of OCRs that were differentially accessible at FDR 5% (light red/blue). Significant correlations at *P*-value<0.05 are marked as * above the bar.

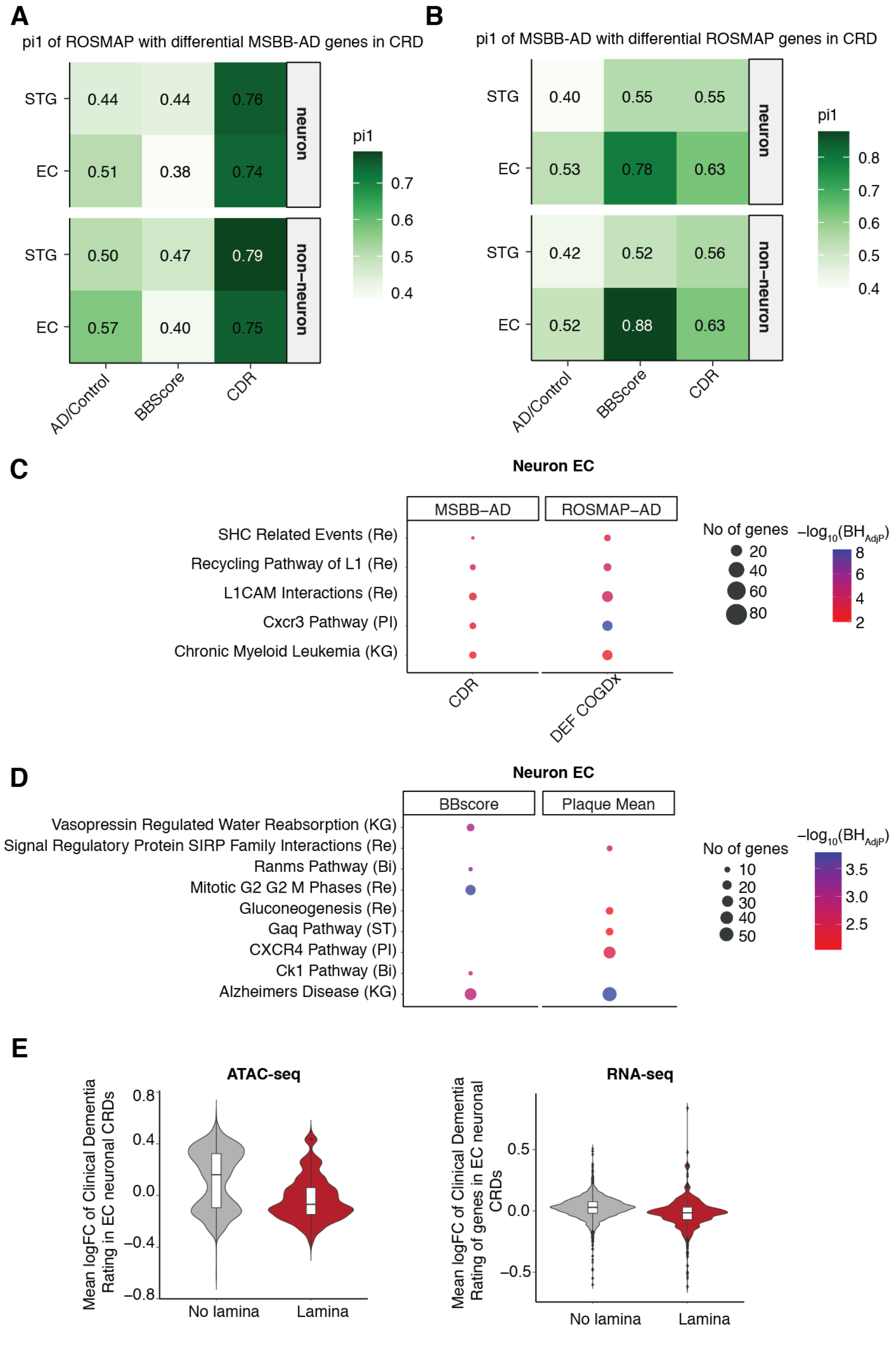


**Fig. S34.**
Differential CRD and differential gene-CRD analysis. *pi1* statistics stratified by cell type and brain regions and estimated as the proportion of non-null tests performed on a vector of

differential $ABC_{mapped}$ genes CRD test nominal *P*-values in the **(A)** replication ROSMAP dataset discovered in MSBB-AD dataset **(B)** replication MSBB-AD dataset discovered in ROSMAP dataset. **(C)** Dot-plot showing enrichment of top five pathways in $ABC_{mapped}$ genes in EC neuronal differential CRDs associated with CDR in MSBB-AD and definite cognition decline in ROSMAP datasets. These sets of differential CRDs were obtained by taking intersection of results from differential CRD test and differential $ABC_{mapped}$ genes CRD test at FDR 5%. **(D)** Dot-plot showing enrichment of top five pathways in $ABC_{mapped}$ genes in EC neuronal differential CRDs associated with BBscore and Plaque mean in MSBB-AD dataset. These sets of differential CRDs were obtained by taking intersection of results from differential CRD test and differential $ABC_{mapped}$ genes CRD test at FDR 5%. **(E)** Violin plots depicting the distribution of mean logFC related to clinical dementia in OCRs inside CRDs and $ABC_{mapped}$ genes to CRDs stratified by their overlap status as “no lamina” and “Lamina” for CRDs that have no overlap and overlap and with lamina segments from ESC cells respectively.
